# Spatial transcriptomics using combinatorial fluorescence spectral and lifetime encoding, imaging and analysis

**DOI:** 10.1101/2021.06.22.449468

**Authors:** Tam Vu, Alexander Vallmitjana, Joshua Gu, Kieu La, Qi Xu, Jesus Flores, Jan Zimak, Jessica Shiu, Linzi Hosohama, Jie Wu, Christopher Douglas, Marian Waterman, Anand Ganesan, Per Niklas Hedde, Enrico Gratton, Weian Zhao

**Author notes:** These authors contributed equally to this work (Tam Vu, Alexander Vallmitjana, Joshua Gu).

## Abstract

Multiplexed mRNA profiling in the spatial context provides important new information enabling basic research and clinical applications. Unfortunately, most existing spatial transcriptomics methods are limited due to either low multiplexing or assay complexity. Here, we introduce a new spatialomics technology, termed Multi Omic Single-scan Assay with Integrated Combinatorial Analysis (MOSAICA), that integrates *in situ* labeling of mRNA and protein markers in cells or tissues with combinatorial fluorescence spectral and lifetime encoded probes, spectral and time-resolved fluorescence imaging, and machine learning-based target decoding. This technology is the first application combining the biophotonic techniques, Spectral and Fluorescence Lifetime Imaging and Microscopy (FLIM), to the field of spatial transcriptomics. By integrating the time dimension with conventional spectrum-based measurements, MOSAICA enables direct and highly-multiplexed *in situ* spatial biomarker profiling in a single round of staining and imaging while providing error correction removal of background autofluorescence. We demonstrate mRNA encoding using combinatorial spectral and lifetime labeling and target decoding and quantification using a phasor-based image segmentation and machine learning clustering technique. We then showcase MOSAICA’s multiplexing scalability in detecting 10-plex targets in fixed colorectal cancer cells using combinatorial labeling of only five fluorophores with facile error-correction and removal of autofluorescent moieties. MOSAICA’s analysis is strongly correlated with sequencing data (Pearson’s r = 0.9) and was further benchmarked using RNAscope™and LGC Stellaris™. We further apply MOSAICA for multiplexed analysis of clinical melanoma Formalin-Fixed Paraffin-Embedded (FFPE) tissues that have a high degree of tissue scattering and intrinsic autofluorescence, demonstrating the robustness of the approach. We then demonstrate simultaneous co-detection of protein and mRNA in colorectal cancer cells. MOSAICA represents a simple, versatile, and scalable tool for targeted spatial transcriptomics analysis that can find broad utility in constructing human cell atlases, elucidating biological and disease processes in the spatial context, and serving as companion diagnostics for stratified patient care.

## Introduction

Cell fate and cell-cell, cell-niche interactions are tightly regulated in space at both genetic and tissue and system level to mediate organ development, tissue homeostasis and repair, and disease appearance and progression. Therefore, spatial transcriptomics that profile gene expression landscape at the single-cell level in tissues in a 3D spatial context as shown in this work represents a new frontier in biological research and precision medicine^1-8^. For instance, spatial transcriptomics techniques can (a) help realize the vision of the human cell atlas in generating “high-resolution and comprehensive, three-dimensional reference maps of all human cells in the body”, (b) determine molecular mechanisms that govern cell fate, state, lineage and cell cooperation in tissue formation in developmental biology and regenerative medicine, (c) investigate the biological changes associated with different diseases in a spatial-dynamic fashion and to uncover disease molecular mechanisms and discover disease biomarkers, and (d) characterize the complexities of tissue biopsy (e.g. tumor) in clinical pathology to inform personalized disease diagnosis and therapeutic intervention in the era of precision medicine. Spatial transcriptomics tools need to be able to assess multiple transcripts within the same cell and sample in a highly multiplexed fashion due to the heterogeneous gene expression and many different cell identities/states exist in a particular tissue. Furthermore, patient derived materials are often available in limited quantity and generating many sections to test for different markers separately is tedious and non-feasible.

A major bottleneck in spatial transcriptomics is the lack of tools that can be both easy-of-use and highly multiplexing^7-11^. Conventional tools for in situ analysis including fluorescence in situ hybridization (FISH) (e.g., LGC Stellaris™) can only detect 3-4 targets at a time because of the limited number of spectral channels in fluorescence microscopes^12-14^. Conventional methods for *in situ* profiling of transcripts are further confounded by the autofluorescent moieties in tissue preparations including clinical biopsies. Recent single-cell RNA sequencing (scRNAseq) methods provide information on the presence and identity of transcripts in single cells but lack the critical spatial context needed to understand complex heterogeneous tissue ^15-17^. Spatial transcriptomic methods that are based on sequential labeling, stripping, and imaging (e.g. seqFISH, MERFISH), branched amplification (e.g. RNAscope™, SABER), or in situ sequencing (e.g. GeoMx™, Slide-seq) are often too complicated, error-prone, time-consuming, laborious and costly to scale up, limiting their broad usage ^18-23^. Furthermore, repeated processing of the same sample can damage tissue structural integrity and target molecules and are often not feasible for precious samples such as clinical biopsies.

In this work we report a new spatial-omics technology termed MOSAICA (Multi Omic Single-scan Assay with Integrated Combinatorial Analysis) that enables direct, highly multiplexed biomarker profiling in the 3D spatial context in a single round of staining and imaging. MOSAICA employs *in situ* staining with combinatorial fluorescence spectral and lifetime encoded probes, spectral- and time-resolved fluorescence imaging, and AI-based target decoding pipeline (Fig. 1). Fluorescence lifetime is a measure of the time a fluorophore spends in the excited state before returning to the ground state and is an inherent characteristic of the fluorophore and its surrounding environment^24^. By utilizing both time and intensity domains for labeling and imaging, we were able to discriminate a large repertoire of spectral and lifetime components simultaneously within the same sample to enable increased multiplexing capabilities with standard optical systems.

**Figure 1.**
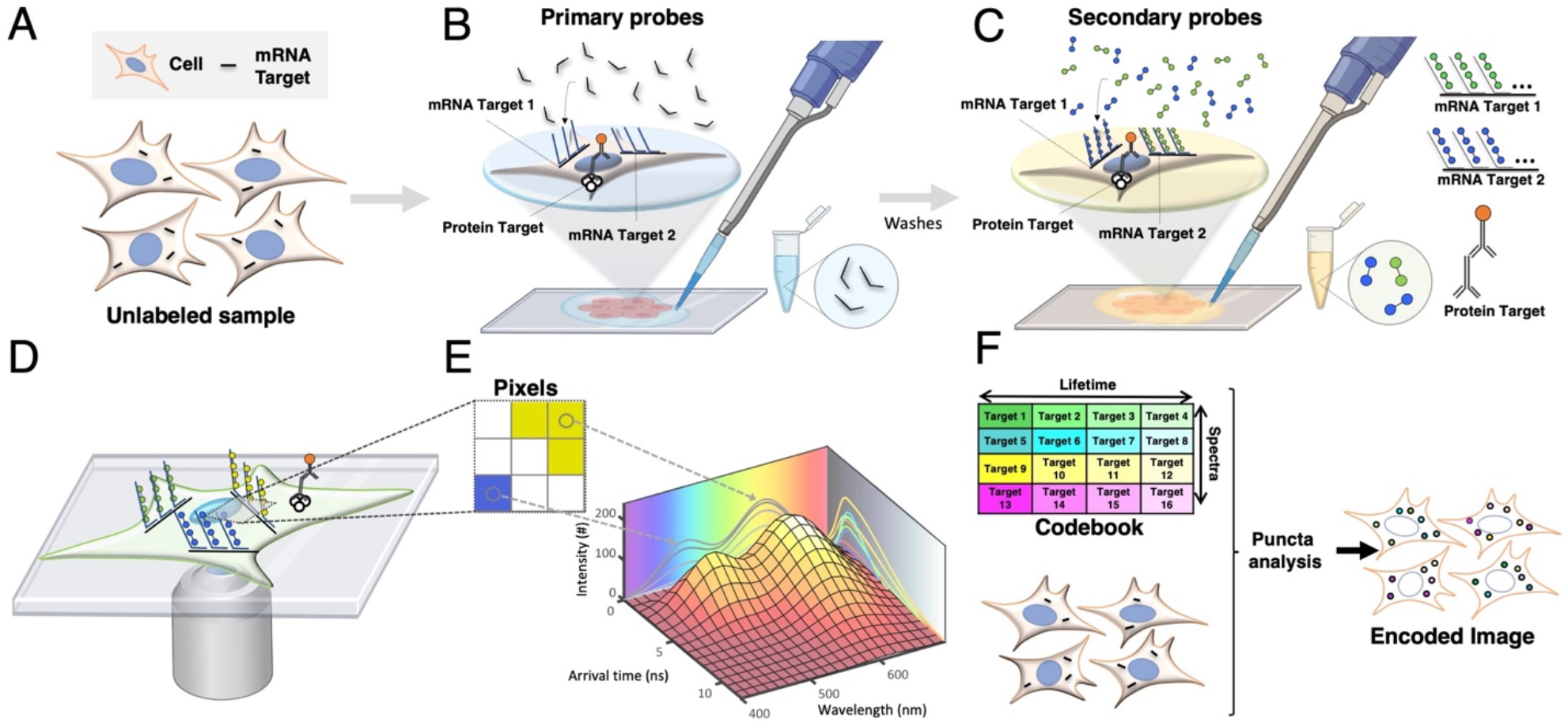
Schematic of the MOSAICA approach for labeling and analysis of spectral and time-resolved components. **A)** Sample(s) can be fixed cells or tissues. RNA transcripts from genes of interest are targeted for detection. Protein targets can be stained too in mRNA and protein codetection. **B)** Primary labeling probes are designed to include two functional regions: a target region which is complementary and can bind to the mRNA target and an adjacent readout region which can subsequently bind to fluorescently labeled oligonucleotides. **C)** Secondary fluorescent probes are added to bind to the primary probes to form different combinations (combinatorial labeling) through a “readout” domain. **D)** Labeled targets are measured under a fluorescent microscope to interrogate the spectral and lifetime characteristics of the labeled moieties. **E)** Phasor analysis is used to identify which fluorophore labels are present in each pixel and puncta. **F)** Labeled targets eliciting the encoded intensity-based or time-based signature are decoded to reveal the locations, identities, counts, and distributions of the present mRNA targets in a multiplexed fashion.

In this study, we described the MOSAICA pipeline including automated probe design algorithm, probe hybridization optimization and validation, combinatorial spectral and lifetime labeling and analysis for target encoding and decoding. Particularly, we developed an automated machine learning-powered spectral and lifetime phasor segmentation software that has been developed to spatially reveal and visualize the presence, identity, expression level, location, distribution, and heterogeneity of each target mRNA in the 3D context. We showcased MOSAICA in analyzing a 10-plex gene panel in colorectal SW480 cells based on combinatorial spectral and lifetime barcoding of only five generic commercial fluorophores. Using this model, we illustrated the multiplexing scalability and MOSAICA’s ability to correct for stochastic nonbinding artifacts present within the sample. We further demonstrated MOSAICA’s utility in improved multiplexing, error-correction, and autofluorescence removal in highly scattering and autofluorescent clinical melanoma FFPE tissues, demonstrating its potential use in tissue for cancer diagnosis and prognosis. To further reveal the potential of MOSAICA, we demonstrated its multi omics capability with simultaneous co-detection of protein and mRNA in colorectal SW480 cells. MOSAICA is rapid, cost-effective, and easy-to-use and can fill a critical gap between conventional FISH and sequential- and sequencing-based techniques for targeted and multiplexed spatial transcriptomics.

## Results

### MOSAICA Workflow

In a typical MOSAICA workflow (Figure 1), primary oligonucleotide probes designed to specifically bind to mRNA targets with a complementary target region (25 to 30 base long) are incubated with fixed cell or tissue samples (Fig. 1A,B). These primary probes also contain an adjacent adaptor region consisting of two readout sequences for modular secondary probe binding. In this study, double ended secondary probes with fluorophores on each end are hybridized to the readout region on the primary probes (Fig. 1C). Through combinatorial labeling, each target is encoded with a dye with a distinct spectrum and lifetime signature. The labeled samples are then imaged using a confocal microscope (Fig. 1D) that is equipped with both spectral and lifetime imaging capabilities. Both spectral and fluorescence lifetime data will be captured and visualized using phasor plots (Fig. 1E). Our automated machine learning algorithm and a codebook then reveals the locations, identities, counts, and distributions of the present mRNA targets in a 3D context (Fig. 1F).

### Probe design pipeline

To rapidly design oligonucleotide probes for each gene, we modified the python platform, OligoMiner^25^, a validated pipeline for rapid design of oligonucleotide FISH probes. Briefly, as shown in Supplementary Figure 1A, using the mRNA or coding sequence (CDS) file of the target gene, the blockParse.py script will screen the input sequence and output a file with candidate probes while allowing us to maintain consistent and customized length, GC, melting temperature, spacing, and prohibited sequences. Using Bowtie2, the candidate probes are rapidly aligned to the genome to provide specificity information that is used by the outputClean.py script to generate a file of unique candidates only. The primary probes comprise complementary sequence of typically 27-30 nucleotides (nt) and are designed mostly within the CDS region, which has fewer variation than the untranslated region (UTR)^20^. We wrote a script, seqAnalyzer.py, to automate the alignment of primary probes to sequencing data (Fig. S1B) so that probes that aligned to regions of lower read counts would be discarded. Furthermore, primary probe “read-out” domains and secondary probes (typically 15 to 20 nt long) are designed to be orthogonal to each other to avoid off-target binding. Libraries and databases of over 200,000 orthogonal sequences are available online and we have simply used those that have been previously validated^26^. Fluorophores exhibiting distinct spectrum (typically with excitation/emission spectra in the 400-700 nm range) and lifetimes (typically in the 0.3-10 ns range) can be conjugated to oligos which were obtained through commercial vendors (see Methods).

### Probe labeling validation and optimization

We first investigated the specificity of our labeling condition using a simple cell mixture model comprising wild-type HEK293T-X cells and HEK293T-X cells engineered with mNeonGreen (Fig. S2A) by detecting mNeonGreen mRNA as the target gene. Since only fluorescent mNeonGreen positive cells can express the corresponding mRNA transcripts, this cell mixture model provides a straightforward tool to assess specificity and nonspecific binding. Using a Nikon epifluorescence microscope to image the samples following staining with primary and secondary probes (all probe sequences used in this study are provided in Table S3), we detected on average 43.5 puncta per mNeonGreen positive cell (n = 76 cells) and 0.25 puncta per wild-type cell (n = 164) (Fig. S2B,C), indicating minimal nonspecific binding with our probe labeling strategy. To further validate the baseline level of nonspecific binding, we included a negative control with the primary probe designed towards dopachrome tautomerase (DCT), a gene in the mouse genome that is not expressed in our HEK293T-X model system, along with a condition with secondary probes only. Similarly, an average of 43.5 puncta per cell was detected for the mNeonGreen cells while the wild type and negative controls a mean of 2.5 puncta per cell was detected with a lower signal to noise (Fig. S2B,C). We next optimized labeling efficiency by testing the number of primary probes and incubation times of primary probes and secondary probes (summarized in Fig. S3). We determined our optimal condition to comprise a minimal of least 12 primary probes for each target mRNA (Fig. S3A,B) (in practice, we always maximize the number of primary probes per mRNA depending on the size of mRNA. Indeed, 40 primary probes per channel per mRNA were subsequently used in this study) with incubation time of 16 hours for primary probe hybridization and 1 hour for secondary probe hybridization, respectively (Fig. S3C,D), which were used in subsequent experiments.

### Imaging and phasor analysis

Lifetime imaging is a tool that measures the spatial distribution of probes with different fluorescence lifetime. Samples are stimulated with modulated or pulsed lasers at a particular frequency, typically around the 40 to 80 MHz, which allows the fluorescence to decay within the stimulated period, typically in the ns range. After acquiring for sufficient time, i.e., after enough laser pulses or periods, one can construct a histogram of photon arrival times at each pixel. The shape of this histogram has a rapid rise, followed by a faster or slower decay which is characteristic of the fluorescent molecule(s) present in the pixel. To model this decay data, an exponential decay model can be fitted or alternatively one can make use of the fit-free phasor approach^27,28^. We used this second approach because it requires no a priori knowledge of an underlying model (i.e. number of fluorescent species at the pixel) and it is computationally inexpensive in virtue of the Fast Fourier Transform algorithm. The phasor transform extracts two values from the decay curve that characterize the shape (and importantly not the size, so that the transform is independent of the amount of photons) and these two values, namely S and G, correspond to the two coordinates of the pixel on the phasor plot (see equations in the supplemental material). The values are obtained by an integral of the product of the decay of the two trigonometric functions, sine and cosine, fit in the stimulation period, and they correspond to the first order terms of the Fourier Series decomposition of the decay curve.

Similarly, if one uses a spectral detector, i.e., a separate detector for different spectral bands, then for each pixel, one can obtain another histogram, in this case with the number of photons arriving in each channel, i.e., at each wavelength. This curve can also be transformed to an analogous spectral phasor space to map the recorded spectra at each pixel onto the 2D spectral phasor space^29^. Combining the lifetime measurement with a spectral detector, one effectively has a 5-dimensional space in which to characterize each pixel. On top of the spatio-temporal coordinates (x,y,z,t), each pixel now carries information in 5 additional coordinates: its intensity value (however many photons arrived at that pixel), the two phasor coordinates for the lifetime phasor transform, and the two phasor coordinates for the spectral phasor transform. A typical image, on the order of 10^6^ pixels, obtained with this method provides 10^6^ points in this 5D space^30^. If the sample presents different populations of fluorescent molecules at different locations, the pixel phasor data at these different locations map to different positions in this phasor space and a clustering technique can be used to resolve each population.

Figure 2 depicts the spectral/lifetime analogy for fluorescence microscopy using the phasor approach. As an example, it uses a hypothetical experiment where 4 different target genes are targeted with 4 fluorescent species. Of the 4 species, we construct the example so that two fluorescent species emit in one color and the other two in another color. At the same time, within each color, one has a short lifetime and the other has a long lifetime. This hypothetical sample is excited, and the individual photons are detected at each pixel (Fig. 2A). In each pixel, we accumulate enough photons to build a spectral histogram and a lifetime histogram (Fig. 2B). These curves are phasor-transformed to reveal two distinct populations in the phasor space, corresponding to the two colors and the two lifetimes. By means of our previously published automatic clustering using machine learning^31^, we identify these populations and return to the image space to label each pixel depending on the group it belongs to in the phasor space (Fig. 2C). By combining the spectral and lifetime information, we have automatically segmented the image into regions, i.e., identified the pixels that belong to the different species (Fig. 2D). Again, note that in this example in Figure 2, we have chosen the probes to be the most convoluted case possible; one couple shares a similar spectrum and the other couple shares another spectrum. At the same time, one of the members of either couple share a similar lifetime and the other two members of either couple share another lifetime. This is the reason why even if there are four distinct fluorescent probes, only two spectral populations are detected both in the spectral and lifetime phasor space, and the combinations of these two populations yield to the four distinct groups. The four probes cannot be resolved unless both the lifetime and spectral information are accessed.

**Figure 2.**
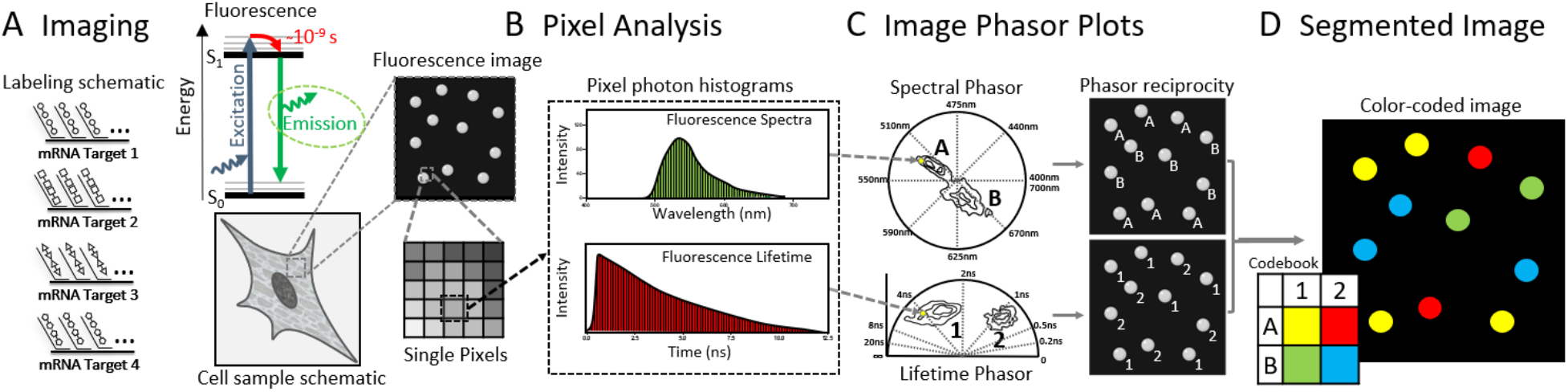
Image and phasor analysis with spectrum and lifetime analysis in MOSAICA. **A)** As an example, four different probes are used to target four different genes. The fluorescence is collected using the spectral and FLIM instrument to form images where each pixel carries information of the spectra and lifetime. **B)** At each pixel we compute the photon distribution in the spectral and temporal dimension. The phasor transform maps these distributions in each pixel to a position on the phasor space. **C)** The phasor plots reveal the presence of different populations. These populations are identified and then mapped back to the original image. **D)** We color code the pixels based on the combination of the two properties. This allows us to separate by lifetime probes that were emitting with similar spectra and vice-versa, separate by spectra probes that fluoresce with similar lifetimes.

### Combinatorial target spectral and lifetime encoding and decoding

In the previous section, we showed how by combining the time dimension with the spectral dimension, we can increase the number of possibilities and therefore enhance the multiplexing capabilities squaring the number of targets that can be resolved. To further increase multiplexing and improve detection efficiency, we employ combinatorial labeling, a method in which targets are labeled with two or more unique fluorophores, to greatly increase the base number of targets we can label with a given number of fluorophores/probes. To illustrate this concept, here we demonstrate a minimal exemplary working example of combinatorial labeling where two probes are used to label three targets. In this situation, each probe labels one target and the third target is labeled with both probes simultaneously. Figure 3 shows a real case with such configuration, both for spectra and for lifetime. In Figure 3A, the cartoon represents the case of using two probes with distinct spectra. When imaging this sample, we can use two spectral channels, Figure 3B/C, where some targets appear in only one channel, other targets appear in only the other channel and the target that is labeled with both probes appears in both channels. All targets are then detected and color-coded depending on their presence in one channel, the other or the two simultaneously (Fig. 3D) and the overall counts of each combination in the field of view can be provided (Fig. 3E).

**Figure 3.**
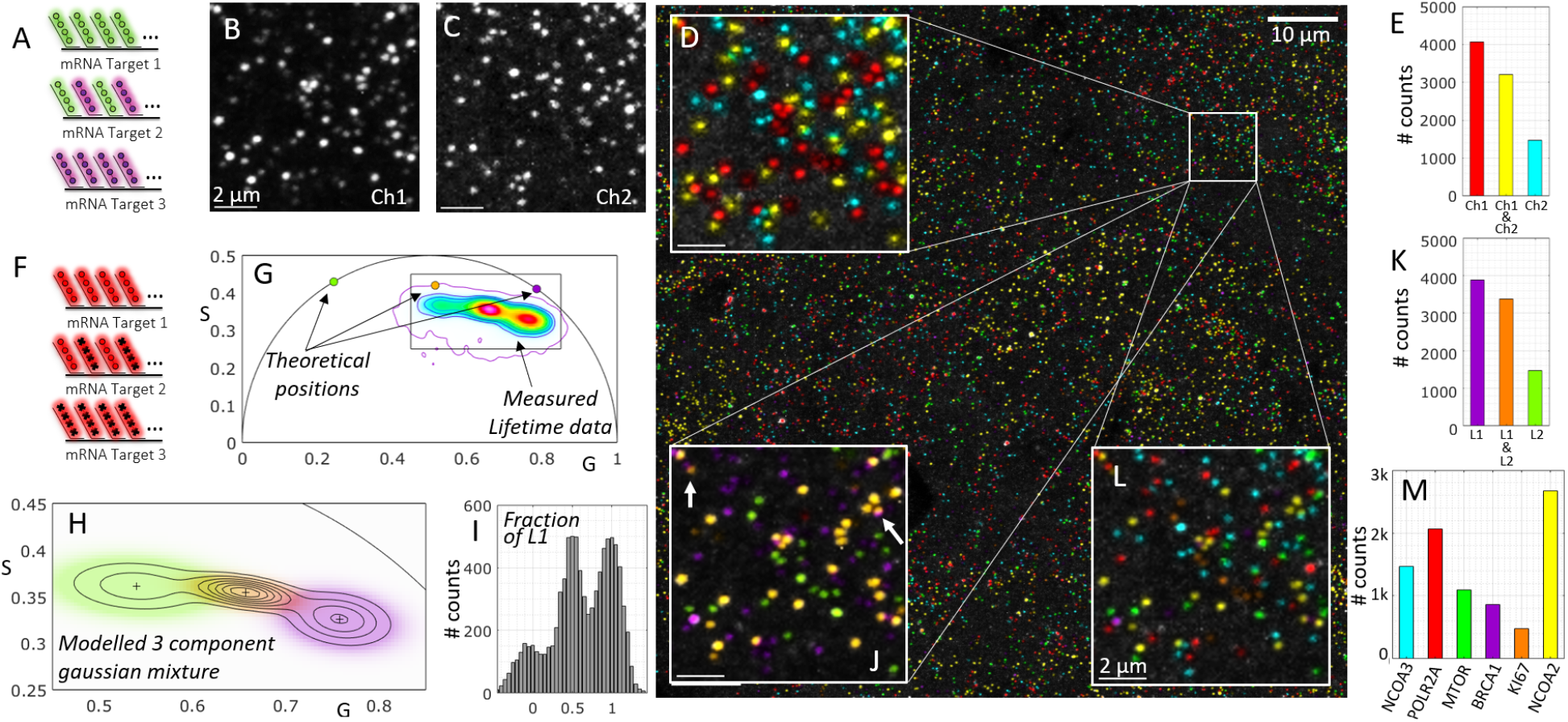
Working example of combinatorial labelling of three mRNA targets with two probes. **A)** Three different target genes are tagged using two probes with different spectra. Targets 1 and 3 are tagged each with one probe, Target 2 is tagged with both simultaneously. **B**,**C)** The fluorescence is collected in the two expected spectral channels for the known emission of the two probes. **D)** The maximum projection of the two channels is shown and pseudocolored depending on the presence in the respective channels (as an inset within the whole field of view. **E)** The actual counts of each target within the whole field of view. **F)** As a parallel example, three different target genes are tagged using two probes with different lifetime. Targets 1 and 3 are tagged each with one probe, Target 2 is tagged with both simultaneously. **G)** The phasor plot presents three populations, corresponding to the pixels with the three combinations; the two components by themselves plus the linear combination falling in the middle. **H)** Machine learning clustering technique is used to identify the groups (Gaussian mixture model). **I)** The multicomponent method is used to extract the fraction of one of the components in each detected puncta. **J)** The same inset is shown with the pseudocoloring now depending on the lifetime clustering. **K)** The counts for each lifetime cluster in the whole field of view. **L)** The combination of the information in both the spectral and the lifetime dimension yields a final 6-plex. **M)** The overall counts for the 6-plex detection including POLR2A (Alexa647 & ATTO565), MTOR (ATTO647 & ATTO565), KI67 (Alexa647 & ATTO647), BRCA1 (Alexa647), NCOA2 (ATTO647), NCOA3 (ATTO565) with the appropriate genes that correspond to each combination. Experiments were conducted with cultures of mNeon green cells.

Similarly, we show a case in which the targets are now labeled with two probes that have similar spectra but different lifetimes (Fig. 3F). In this case, we also introduce the use of the phasor approach to reveal the three expected populations, the pixels that contain both probes appear in the midpoint between the phasor positions of the pixels that contain only one of the probes. Figure 3G shows the phasor distribution obtained from the same field of view as in the spectral example, in which we also show the theoretical locations of the probes (corresponding to Alexa647 and ATTO647 with respective lifetimes of 1 ns and 3.5 ns). As is expected in real experimental conditions, there are additional fluorescent components in the sample. We broadly refer to the bulk of these additional components as autofluorescence, which pulls the data away from the expected positions and converges to the mean phasor position of the autofluorescent components. We have previously shown that the Gaussian Mixture Models is the most optimal machine learning clustering algorithm to model phasor data^31^, and we use this machine learning technique to infer the phasor locations of the probe combinations (Fig. 3H). We can now successfully classify each pixel of the original image into one of the clusters and obtain a probability of belonging to each, i.e., the posterior probability of the model. This allows us to color-code the transcripts depending on their assignment to one of the 3 clusters (Fig. 3J) and obtain the counts of the three lifetime components (Fig. 3K). Additionally, we apply our lifetime multicomponent analysis technique^32^ in which for each detected puncta, we estimate the presence of one of the lifetime components, in this case lifetime1 (Alexa647, purple in the figure), to obtain the expected result; that there are clearly three populations with respective fractions centered around [0, 1/2 and 1] (Fig. 3I).

In the general case, we combine the lifetime and spectral dimensions, and we perform the clustering of the data in a 4D spectral/lifetime phasor space. The clustering technique has the power to not only identify which puncta belong to each cluster but also to assign a probability of belonging to that cluster, which can be used to quantify the certainty of the labeling. For example, in the inset in Figure 3J, we show two cases of puncta that have relatively low confidence in the cluster assignment; they are depicted with blended colors because they fall in the regions of the phasor space where the two clusters are merging.

In this combinatorial example in Figure 3, the three clusters in the lifetime domain multiplexed with the channel-based in the spectral domain yield a 6-plex image using only 3 probes (Fig. 3L,M). The specific genes targeted for this experiment with the combined probes were POLR2A (Alexa647 & ATTO565), MTOR (ATTO647 & ATTO565), KI67 (Alexa647 & ATTO 647), BRCA1 (Alexa647), NCOA2 (ATTO647), NCOA3 (ATTO565). In the general combinatorial experiment using couples of N probes the total number of possible target genes grows quadratically:

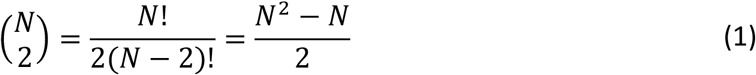

### Simultaneous 10-plex mRNA detection in fixed colorectal cancer SW480 cells using MOSAICA

We next applied MOSAICA to a 10-plex panel of mRNA targets in colorectal cancer SW480 cell culture samples. This cell line was chosen because its xenograft model exhibits spatial patterns of heterogeneity in WNT signaling^33^, which will allow us to study tumorigenesis in the spatial context and potentially identify cancer stem cell populations in colorectal cancer in future. Here, we selected this model as a validation platform to demonstrate the multiplexing scalability and error correction capabilities of our approach. We began by first identifying a set of 10 genes with known expression levels from our bulk sequencing data. Using the aforementioned probe design pipeline, we designed 80 probes (two pairs of 40 probes) for each gene: BRCA1, BRCA2, CENPF, CKAP5, KI67, MTOR, NCOA1, NCOA2, and NCOA3. These genes were chosen due to their housekeeping status or involvement in tumorigenesis in colorectal cancer. By encoding each gene with a distinct combination of two fluorophores, we generated a codebook of 10 labelling combinations from only five fluorophores following Equation 1: 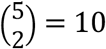(Fig. 4A) (see Table S2 for the probes used for each target). To assess the baseline nonspecific binding events of our assay, we included a negative probe control sample which was labelled with primary probes not targeting any specific sequence in the human genome or transcriptome but still containing readout regions for secondary fluorescent probes hybridization (Fig. 4A right). Matching numbers and concentrations of primary and secondary probes that were used in the 10-plex panel were used in this sample.

**Figure 4.**
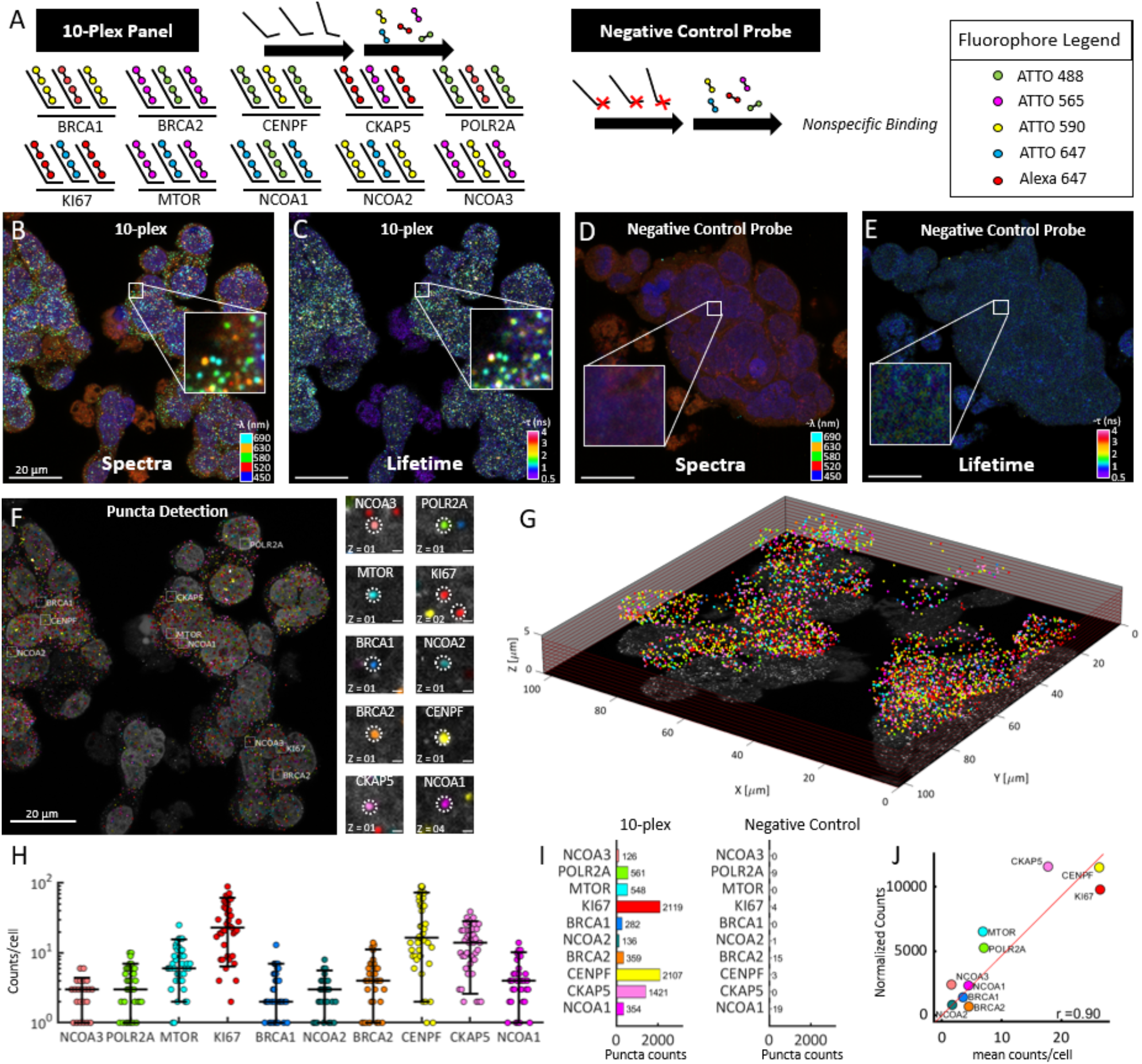
Simultaneous 10-plex detection of genes in colorectal cancer SW480 cells in a single round of labeling and imaging. **A)** 10 different types of gene transcripts are labeled with primary probes followed by respective and complementary fluorescent secondary probes. Each gene is labeled with a combination of 2 different fluorophores for 10 combinations. Negative control probes (mNeonGreen, DCT, TYRP1, and PAX3) targeting transcripts not present in the sample were used with their respective secondary fluorophore probes. **B)** Spectral image (max-projection in z) of a field of view of the labeled 10-plex sample (5-channel pseudo coloring). **C)** Lifetime image (max-projection in z) of a field of view of the labeled 10-plex sample (phasor projection on universal circle pseudo coloring). **D)** Spectral image of the labeled negative control probe sample. **E)** Lifetime image of the labeled negative control probe sample. **F)** Final puncta detection after being processed in our analysis software showing highlighted example puncta of each target (insets, right). Scale bar 20 µm in main image and 1 µm in insets. **G)** 3D representation of the field of view for the 10-plex sample. **H)** Number of puncta detected for each gene expression target in each cell for the labeled 10-plex sample (N=80 cells). **I)** Overall puncta count of each transcript in the 10-plex sample (left) and negative control probe sample (right). **J)** Correlation of detected puncta (mRNA puncta count) vs. RNA-bulk sequencing (normalized counts) is shown for each target with Pearson r of 0.9.

Figure 4B depicts a spectral image overlay (five fluorescent channels including DAPI) of the labeled 10-plex SW480 sample. Additionally, in the same measurement, the orthogonal lifetime information attained by interrogating each pixel for their lifetime components. These pixels were phasor-transformed and pseudo-colored based on their projected phasor coordinates on the universal circle, creating the image depicted in Figure 4C. In doing so, both dimensions of data can now be simultaneously accessed to determine which cluster of pixels meet the appropriate and stringent criteria for puncta classification. Similarly, Figure 4D/E depict the merged composite spectral and lifetime images of the corresponding negative control probe sample. Figure 4F depicts the now detected pseudo-colored clusters which were successfully classified as one of the RNA markers. A representative inset image for each marker and its targeted detection is provided on the right. With 40 Z-stack images, we generated a 3D spatial distribution of the field of view to visualize the spatial analysis in a 3D context (Fig. 4G).

MOSAICA employs an error-correction strategy by gating for specific and pre-encoded fluorophore combinations and rejecting any fluorescent signature which do not meet these criteria. Specifically, in addition to the population of pixels that represents the decodified puncta as mentioned above, there was a population of pixels of various shapes and sizes which did not fit the criteria. Simulations to look into overlapping and inconsistent signals were run (Fig S4A,B). Plotted in Supplementary Figure 4C are the total detected puncta in this experiment, separated into populations depending on our classification criteria. Of the total detected puncta (n = 13,521), a considerable fraction n = 4,488 (33%), were observed emitting fluorescence only in a single channel and was indicative of nonspecific binding events and/or autofluorescence moieties. As previously characterized by several groups, nonspecific binding events is an inherent issue with smFISH which arises from the stochastic binding of cDNA probes towards cellular components such as proteins, lipids, or nonspecific regions of RNA and follow a random distribution^14^. When combined with native autofluorescence moieties which can also exists as diffraction-limited structures that emits a strong fluorescent signal in any single channel, this group represents a confounding problem for standard intensity-based measurements and analysis because they share similar SNR and intensities to real labeled puncta and cannot be differentiated. This is the main benefit of implementing the combinatorial encoded criteria which rejects this large amount (around 33%) of stochastic and nonspecific binding labeling events as well as any event eliciting a lifetime signature that deviated from the utilized commercial fluorophores. Finally, we also observed a relatively small group of puncta emitting fluorescent signal across more than two spectral channels but still eliciting the same spectral and lifetime signatures as the utilized fluorophores, n = 880. To characterize this population, we performed a simulation running 20,000 iterations of various puncta densities and fitted the corresponding exponential model that characterizes the probability of puncta overlap (described in Methods section and Supplementary Figure 4A and B). We attained an interval for the fraction of lost puncta due to optical crowding ranging from 2.1% to 6.6% which accounts for the 880 puncta or 6% of the total detected puncta. We name this group the overlapped in Supplementary Figure 4C.

The number of puncta detected for each target gene in each cell for the labeled 10-plex sample was plotted (Fig. 4H). Figure 4I plots the total number of detected puncta for the labeled 10-plex sample split into the different genes classified using MOSAICA phasor analysis with combinatorial labeling. In comparison, we also show the MOSAICA pipeline results with the negative control sample obtaining counts of less than five per thousand mainly due to noise in the images. To validate these puncta count, we compared them to matching RNA-seq data from the same cell type. Shown in Figure 4J is a scatter plot of the average mRNA puncta count for each cell plotted against the normalized counts from DESeq2 of our bulk RNA-sequencing data for each gene. When fitting with a straight line we obtained a Pearson correlation of r = 0.904, indicating a significant positive association between the two methods. The number of counts per cell was obtained by simply dividing the total number of detected genes into the number of cells obtained by 3D segmentation of the individual nuclei stained with DAPI. The individual cell edges were estimated by growing the edges of the nuclei until convergence (see Methods section). A total of 80 cells were analyzed for this experiment.

To further evaluate the detection efficiency, we performed benchmarking tests with our method against LGC Stellaris™and RNAscope™which are commercial gold standard FISH methods (Fig. S5). Using the housekeeping gene, POLR2A, as an exemplary target, we found a significant association between the number of detected puncta by our method and LGC Stellaris™(t-test p-value = 0.4). When compared to RNASCOPE™, we observed that for this cell type and target, both our assays and LGC Stellaris™did not correlate significantly (p = 7.8⨯10^−4^ and p = 3.4⨯10^−4^), indicating a discrepancy in detection efficiency between the two methods. We attribute this difference to MOSAICA and LGC Stellaris™utilizing a direct labeling and amplification-free method while RNASCOPE™utilizes a tyramide signal amplification reaction which generates thousands of fluorophore substrate per transcript and can lead to overlapping puncta or undercounting of detected puncta. Together, these data show MOSAICA can robustly detect target mRNAs of broad dynamic range of expression levels from single digit to hundreds of copies per cell.

### Multiplexed mRNA analysis in clinical melanoma skin FFPE tissues

We next investigated whether MOSAICA can provide multiplexed mRNA detection and phasor-based error-correction to clinically relevant and challenging sample matrices. Assaying biomarkers *in situ* in tissue biopsies has great clinical values in disease diagnosis, prognosis and stratification including in oncology^34-36^. Specifically, we applied a mRNA panel consisting of KI67 (indicative of cell proliferation), POLR2A, BRCA1, MTOR, NCOA2, and NCOA3 to highly scattering and autofluorescent human melanoma skin biopsy FFPE tissues obtained from and characterized by the UCI Dermatopathology Center. Using the same probe design pipeline, primary probes were encoded with a combination of two fluorophores for each gene to exhibit a unique fluorescent signature (Fig. 5A left).

**Figure 5.**
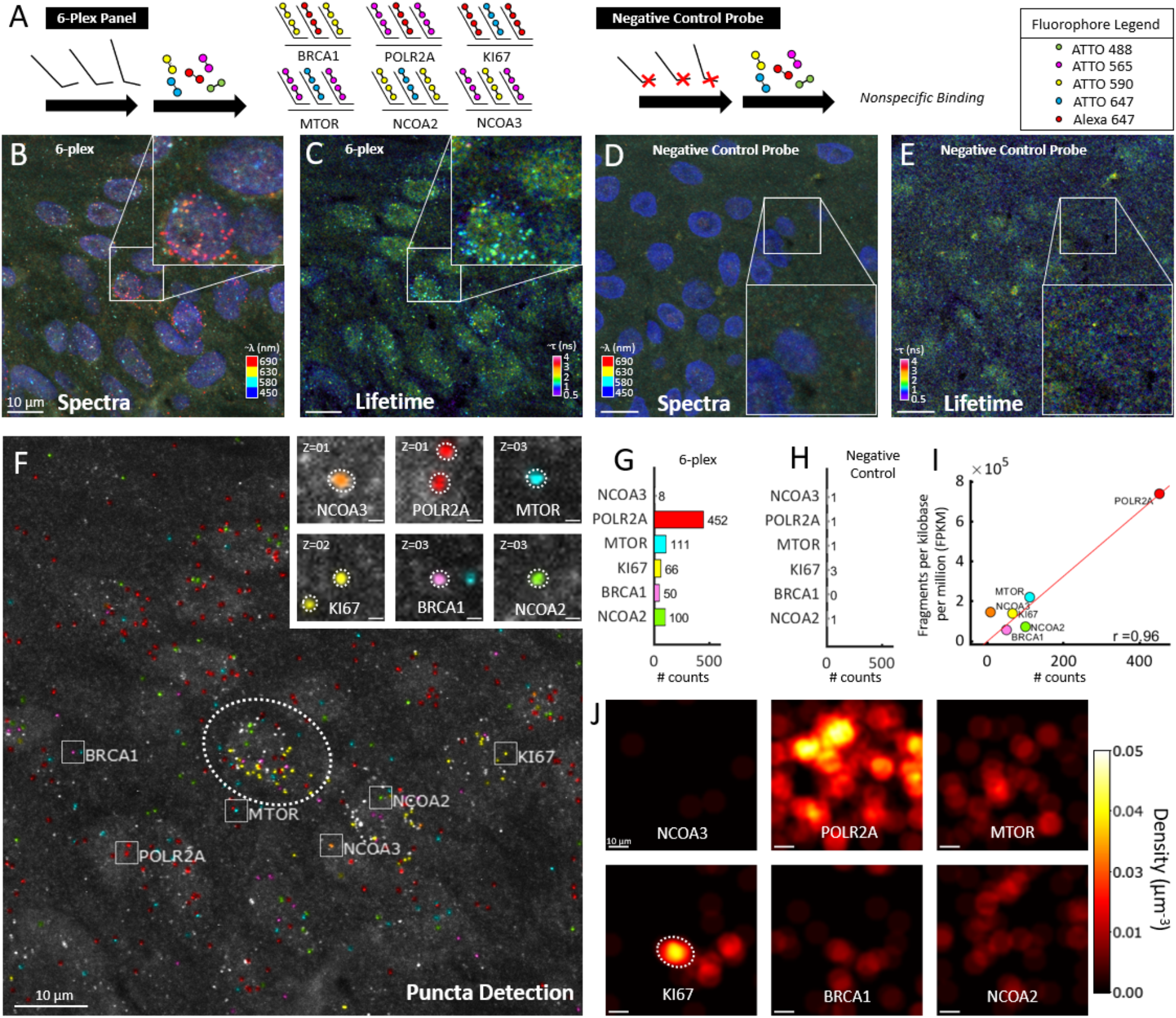
Multiplexed mRNA detection in epidermis region of human skin melanoma FFPE tissue. **A)** 6 different types of gene transcripts were labeled with primary probes followed by respective and complementary fluorescent secondary probes. Each gene was labeled with a combination of 2 different fluorophores for 10 combinations. Negative control probes targeting transcripts not present in the sample were used with their respective secondary fluorophore probes. **B)** Spectral image (max-projection in z) of a field of view of the labeled 10-plex sample (3 channel pseudo coloring). **C)** Lifetime image (max-projection in z) of a field of view of the labeled 10-plex sample (phasor projection on universal circle pseudo coloring). **D)** Spectral image of the labeled negative control probe sample is depicted. **E)** Lifetime image of the labeled negative control probe sample. **F)** Final puncta detection of the 6-plex field of view after being processed in our analysis software showing highlighted example puncta of each target (insets, right). Scale bar is 10 µm in the main image and 1 µm in insets. **G)** Overall puncta count of each transcript in the 6-plex sample. **H)** Puncta count for the negative control probe sample. **I)** Correlation of detected puncta (mRNA puncta count) vs. bulk sequencing (fragments per kilobase per million (FPKM)) is shown for each target. **J)** Transcript density in the field of view for each of the genes reveals clustering of specific genes, as an example KI67 appears highly expressed in three cells, one of them marked with a dotted ellipse that corresponds to location in F.

Figure 5B depicts a spectral image overlay (four fluorescent channels including DAPI) of the epidermis region of a labeled 6-plex skin tissue sample. Similarly, as in the previous section, the orthogonal lifetime image was attained after using phasor analysis to create the image depicted in Figure 5C. Figure 5D/E depict the merged composite spectral and lifetime images of the corresponding negative probe sample also in the epidermis region. Figure 5F depicts the pseudo-colored puncta which were successfully classified and identified as their assigned mRNA markers. A representative inset image for each marker and its targeted detection is provided on the right. Using this approach, we observed that a population of puncta consisting of nonspecific, autofluorescent, or unknown sample artifacts were rejected from analysis, (n = 438) or 29% of total detected clusters (Fig. S4D). In addition to this group, we observed MOSAICA rejecting a small group of puncta that emitted fluorescence in multiple spectral channels (n = 168). Since, this fraction (11%) exceeded the optical crowding range (2.1% to 6.6%) our simulations and model permits, we attribute this discrepancy to autofluorescence moieties which had fluorescent signatures that spans a broad emission spectrum and elicits a multispectral response. With conventional intensity-based measurements and analysis, both contaminating groups are inherent image artifacts that compromise the integrity of puncta detection unless complicated quenching steps or additional rounds of stripping, hybridization, and imaging are utilized^14,37^. With MOSAICA, these contaminating artifacts can be accounted for with the integration of spectral, lifetime, and shape-fitting algorithms.

Figure 5G/H plots the total number of detected puncta for the labeled 6-plex sample and the negative control probe sample to highlight the final counts obtained using MOSAICA. We observed MOSAICA platform results with the negative control sample obtaining counts of less than five per thousand. To validate these puncta counts and their relative expressions, we examined the relationship between the decodified puncta with matching bulk RNA-sequencing obtained from The Cancer Genome Atlas (TCGA) database (see Methods section). Shown in Figure 5I is a scatter plot of mRNA puncta count for each cell plotted against fragments per kilobase per million (FPKM). We observed a Pearson correlation of r = 0.959 for this 6-plex sample, indicating a significant positive association between the two methods and the capability for MOSAICA to capture a wide range of differentially expressing markers. Lastly, as shown in Figure 5J, the density map of the detected transcripts provides a visual method to identify spatial localization of clusters of genes, such as KI67 (indicative of proliferating tumor cells) being more prevalent in the dermis region while POLR2A is dispersed throughout the region. Overall, *in situ* profiling biomarkers such as KI67 and their spatial clustering can have diagnostic and prognostic values in malignant diseases and MOSAICA provides a robust platform to profile these markers^38^.

### Simultaneous co-detection of protein and mRNA

Spatial multi-omics analysis including especially simultaneous detection of protein and transcript within the same sample can reveal the genotypic and phenotypic heterogeneity and provide enriched information for biology and disease diagnosis. As a pilot experiment to demonstrate MOSAICA’s potential for multi-omics profiling, we utilized MOSAICA to detect 2 protein targets, Tubulin and Vimentin, and 2 mRNA targets, POLR2A and MTOR in colorectal cancer SW480 cell culture samples (Fig. 6). After staining the sample with the primary antibodies, secondary antibodies were added to fluorescently label the protein targets. After protein labeling, we utilized the same probe design pipeline and labeling strategy for mRNA detection, primary probes were generated and hybridized to the sample after antibody staining. Corresponding secondary probes were hybridized. Figure 6A-F depict the individual channels of the sample with Figure 6G showing the merged channels of the 4-plex panel. As both POLR2A and MTOR are assigned to the 647nm channel and cannot be separated spectrally (Fig. 6D), lifetime analysis is used to separate POLR2A (Fig. 6E) and MTOR (Fig. 6F). As shown in Figure 6H, 190 POLR2A puncta and 63 MTOR puncta were quantified using signal-to-noise ratio. In summary, we have demonstrated MOSAICA as a potential spatial multi-omics tool which harmonizes sample treatment between both labeling processes. MOSAICA utilizes staining protocols with efficient target retrieval, blocking, and pretreatment steps where the viability and labeling of both target RNA sequence and protein markers were not compromised after each assay.

**Figure 6.**
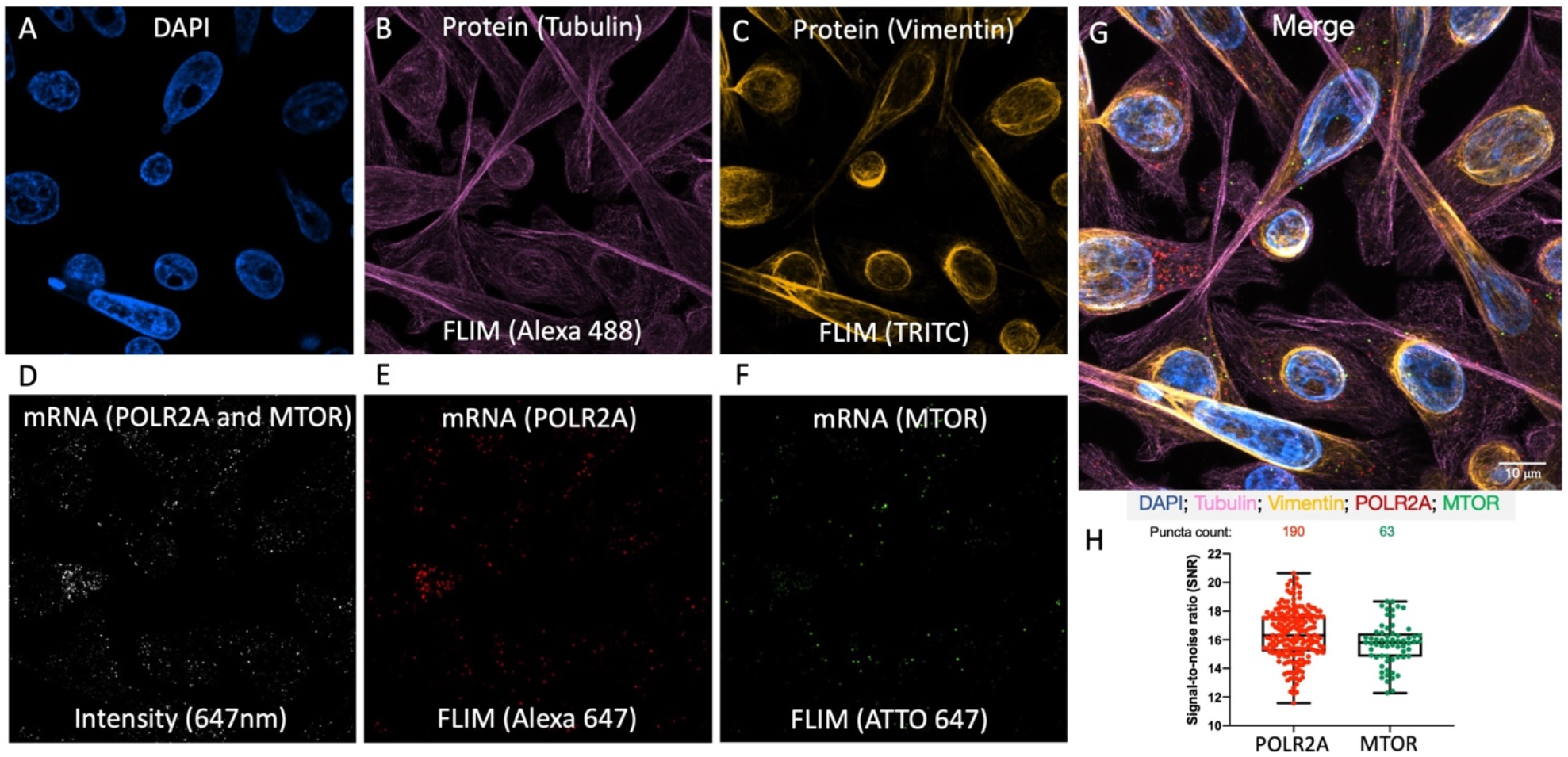
Simultaneous 4-plex co-detection of protein and mRNA in colorectal cancer SW480 cells. **A)** The individual DAPI channel. **B)** Lifetime image of the protein detection of Tubulin (TUBB4A mouse) with Alexa 488. **C)** Lifetime image of the protein detection of Vimentin (VIM rabbit) with TRITC. Secondary antibodies goat anti-mouse Alexa 488 and donkey anti-rabbit TRITC were used respectively. **D)** The 647nm channel intensity image showing mRNA targets POLR2A and MTOR. **E)** The lifetime image showing POLR2A puncta detected with Alexa 647. **F)** The lifetime image showing mTOR puncta detected with ATTO 647. **G)** Merged image of all channels. Scale bar is 10 µm. **H)** Signal-to-Noise and puncta count analysis for the mRNA targets. POLR2A n = 190. MTOR n = 63.

## Discussion

MOSAICA provides a new solution that fills a major gap in the spatialomics field (Fig. S6) by simultaneously achieving both simplicity and multiplexing by enabling direct *in situ* spatial profiling of many biomarkers in a single round of staining and imaging. This solution contrasts existing approaches such as conventional direct labeling approaches (e.g., LGC Stellaris™, RNAscope™, etc.) which can only analyze 3 - 4 targets due to limited spectral channels as well as recent spatial transcriptomic approaches (e.g., seqFISH) which can provide greater multiplexing capabilities but at the expense of many iterations of sample re-labeling, imaging, indexing, and error-prone image registration. MOASICA accomplishes this by uniquely integrating the lifetime dimension with the conventional spectral dimension, employing combinatorial fluorescence spectral and lifetime target encoding, and exploiting machine learning- and phasor-based deconvolution algorithms to enable high-plex analysis and error-correction. This enables MOSAICA to simultaneously access both spectral and lifetime dimensions to achieve higher plex detection without sacrificing assay throughput as well as correcting for autofluorescent moieties and stochastic nonspecific binding artifacts, which are inherent challenges associated with existing intensity-based measurements.

Compared to existing sequential hybridization and imaging approaches, MOSAICA significantly reduces the number of hybridization and imaging rounds required to profile larger multiplexed panels of RNA biomarkers. By imaging every fluorophore and labeled transcript simultaneously rather than sequentially, the need to relabel probes or reimaging samples is precluded which is often the bottleneck for these complex and lengthy assays. For a 10-plex panel, this drastically shortens assay time by at least of factor of 4 – 5x and scales up even more as the panel becomes more multiplexed, making these assays more practical, effective, and affordable for routine laboratory or clinical usage. This is particularly important in clinical settings, where biopsy samples are limited in quantity and the need to avoid repetitive stripping, rehybridization, and image registration to preclude damage to the tissue structure integrity and target molecules is highly critical. In terms of cost, compared to existing commercial platforms where assay ranges from several hundred dollars to thousand dollars per batch of samples, MOSAICA utilizes inexpensive DNA primary probes which can be purchased in batch or as microarrays for a minimal price. Fluorescently conjugated secondary probes can also be used and shared as a common set among many different genes, scaling down costs to several dollars per assay or 10 – 20x less than commercial vendors. Compared to indirect spatial transcriptomic methods that analyze barcoded regions of interest (ROIs) using downstream sequencing (e.g., GeoMx® Digital Spatial Profiler, 10x Genomics Visium), the direct, imaging-based MOSAICA would be advantageous for targeted mRNA profiling with higher spatial resolution (subcellular features or single molecules), simpler workflow, lower cost, and potentially higher throughput (number of tissue sections analyzed per day) with a camera-based system (below). Moreover, the MOSAICA platform utilizes standard fluorescent probes and fluorescence microscopy (several commercial instruments capable of acquiring both spectral and lifetime information are available, e.g., Leica SP8 FALCON, PicoQuant rapidFLIM, ISS FastFLIM, etc., and already exist in many shared core facilities in academia and industry). Fluorescence imaging remains the most familiar and widely used technique in biological research and its minimal requirements of MOSAICA will permit quick and broad adoption in the scientific community.

As a complementary tool to existing spatial-omics methods, MOSAICA broadly enables scientists and clinicians to better study biology, health, and disease and to develop precision diagnostics and treatments. Gene expression is heterogeneous and many different cell identities/states exist in a particular tissue. To fully characterize cells *in situ*, we need to be able to assess multiplexed panels of genes within the same cell, which can be readily addressed by MOSAICA. As such, MOSAICA can help accelerate spatiotemporal mapping experiments to construct 3D tissue cell atlas maps. Additionally, MOSAICA represents a powerful tool for targeted *in situ* validation of RNA sequencing data. scRNAseq returns cell identities in the form of “differentially expressed gene lists” that “define” cell types. However, the clustering process is subjective, variable and error-prone. Multiplex spatial transcriptomics using MOSAICA can validate whether a pattern of gene expression really defines a cell type or conflates multiple cell types. Furthermore, we are currently developing MOSAICA to serve as a clinical companion diagnostic tool for stratified care. Of particular interest is the spatial profiling of the organization and interactions between tumor cells, immune cells and stromal components in tumor tissues that can inform cancer diagnosis, prognosis, and patient stratification^39,40^. With MOSAICA’s ability to analyze numerous markers in a single round, we believe this feature will be particularly attractive for clinicians or researchers working with precious patient derived materials.

Further ongoing work for this platform includes improving multiplexing by scaling up the number of fluorophores and their combinations using our combinatorial-based labeling approach. For instance, Equation A and our simulation (Fig. S4) predict that with different combinations of 10 fluorophores using triple labeling strategy, MOSAICA can barcode and detect 120 mRNA targets. Our immediate next step is to scale our multiplexing capability to about 60-plex, which will bridge the gap between conventional FISH and sequential- and sequencing-based methods and can be rapidly employed to numerous applications in targeted spatial transcriptomic analysis in basic and translational research. In the future, we will further expand our codebook by implementing a Förster resonance energy transfer (FRET)-based barcoding method^41^ by using different FRET fluorophore pairs and tuning the distance between fluorophore donor and acceptor. We can utilize the FRET phenomena as an additional error correction mechanism at the nanometer level to resolve multiple transcripts in the same voxel. Moreover, we will improve imaging throughput with our recently developed camera-based light-sheet imaging ^42^ and hyperspectral imaging^30^. Indeed, MOSAICA is versatile and can be further integrated with other imaging modalities such as super-resolution, expansion microscope and multi-photon imaging techniques ^43-45^ to increase subcellular resolution and enable imaging large highly scattering tissues.

Additionally, we will further develop our image analysis software with a user interface which enables classification and visualization of single-cell phenotypes, spatial organization and neighborhood relationship. Our current software platform’s puncta detection and puncta classifier algorithm can also be improved using modern image processing techniques such as convolutional neural networks with clinical training sets to optimize true and false positive detection of biomarkers. We anticipate on benchmarking these sets with experimental data with stripping methods to determine fidelity of labeled targets. Lastly, we will further improve protein multiplexing in our multi-omics analysis using antibody-DNA conjugates where we can conduct combinatorial labeling and barcoding to scale up multiplexing as we do for the mRNA detection.

## Methods

### Primary probe design

A set of primary probes were designed for each gene. A python code was used to rapidly design the primary probes while controlling various aspects of the probes such as GC content, length, spacing, melting temp, and prohibited sequences. To begin, probes are designed using exons within the CDS region. However, if that region does not provide over 40 probes, the exons from the CDS and UTR regions are used. The candidate probes are then aligned to the genome using Bowtie2, an NGS aligner, to determine if these probes are specific. Probes that are determined specific are then aligned to the RNA sequencing data on the UCSC Genome Browser, further eliminating probes that do not align to regions with an adequate number of reads. While mapping the probes to the genome on the UCSC Genome Browser, the probes are aligned with BLAT (BLAST-like alignment tool). A local BLAST query was run on the probes for the genes in the panel to eliminate off-target hits. For this experiment, each gene had the maximum number of probes that could be designed with our pipeline and requirements. The final primary probe design included 2 assigned readout sequences of secondary probe with a “TTT” connector in between, another connector, then one of the probes specific for the gene. The primary probes were ordered from Sigma Aldrich and pooled together for each gene. The sequences of all probes used in this study are listed in Table S3.

### Secondary probe design

Secondary probe structures were based on the design from the Zhuang group^46^. In short, the 20-nt, three-letter readout sequences were designed by generating a random set of sequences with the per-base probability of 25% for A, 25% for T, and 50% for G. Sequences generated in this fashion can vary in their nucleotide content. To eliminate outlier sequences, only sequences with a GC content between 40% and 50% were kept. In addition, sequences with internal stretches of G longer than 3 nt were removed to eliminate the presence of G-quadruplets, which can form secondary structures that inhibit synthesis and binding. To remove the possibility of significant cross-binding between these readout sequences, algorithms from previous reports were used to identify a subset of these sequences with no cross-homology regions longer than 11 contiguous bases^46^. Probes were then checked with BLAST to identify and eliminate sequences with contiguous homology regions longer than 11 nt to the human transcriptome. From the readout sequences satisfying the above requirements, 16 were selected.

### Cell culture

Human embryonic kidney (HEK293T) cells (632180; Takara) were cultured in DMEM (10-013-CV; Corning) supplemented with 10% FBS (1500-500; Seradigm) and 1% penicillin (25-512; GenClone). Human colorectal adenocarcinoma (SW480) cells were cultured in DMEM with high glucose (SH30081.02; HyClone) supplemented with 10% FBS (1500-500; Seradigm), 1x L-Glutamine (25-509; GenClone), and 1% penicillin (25-512; GenClone). The cells were plated into 8-well chambers and then fixed. The 8-well plates (155409; Thermo Scientific) for HEK293-T and SW480 cells were coated with fibronectin bovine plasma (F1141-2MG; Sigma Aldrich) before seeding cells onto the 8-well plates. All cultures were grown at 37°Cwith 5% CO_2_.

### mNeonGreen cell engineering

A mNeonGreen construct was transfected into HEK293T-X cells with FuGENE HD Transfection Reagent (E2311; Promega). The cells were then selected with puromycin (NC9138068; Invivogen) and Zeocin (AAJ67140XF; Alfa Aesar) 3 days after transfection.

### Preparation of fixed cells in cell chambers

When the cells reached 70% confluency, cells were fixed for 30 minutes using 4% paraformaldehyde (15710; Electron Microscopy Science), then washed with PBS 3 times. The cells were then incubated with sodium borohydride (102894; MP Biomedicals) for 5 minutes and washed with PBS 3 times. 0.5% Triton X-100 (T8787-100ML; Sigma-Aldrich) in PBS was incubated in each well for 5 minutes and cells were washed with 2x SSCT (2x SSC with 0.1% TWEEN® 20 (P9416-100ML; Sigma-Aldrich). For storage, cells were left in 70% ethanol at 4°C.

### Preparation of FFPE tissues

The University of California Irvine IRB approved this study for IRB exemption under protocol number HS# 2019-5054. All human melanoma cases were de-identified samples to the research team at all points and therefore considered exempt for participation consent by the IRB. Fully characterized human patient skin melanoma FFPE tissues with an immune cell score of brisk were obtained from the UCI dermatopathology center then sectioned to 5 µm slices using a rotary microtome, collected in a water bath at 35°C, and mounted to positively charged Fisher super frost coated slides. The tissue sections were then baked at 60°C for 1 hour. For antigen unmasking, slides were deparaffinized, rehydrated then followed by target retrieval (with citrate buffer).

### Primary probe hybridization

Blocking buffer containing 100 mg/ml Dextran sulfate sodium salt (D8906-100G; Sigma-Aldrich), 1 mg/ml Deoxyribonucleic acid from herring sperm (D3159-100G; Sigma-Aldrich), 0.01% Sodium Azide (S2002-100G; Sigma-Aldrich), 0.01% tween, and 15% ethylene carbonate (AC118410010; Fisher Scientific) in 2x sodium saline citrate (SSC) was added to the fixed cells or tissues and incubated at 60°Cfor 8 minutes and then at 37°Cfor 7 minutes. Following this pre-block step, primary probes with 5nM of each probe in blocking buffer were added to the samples and incubated at 60°Cfor 30 minutes and then overnight at 37°C.

### Secondary probe hybridization

Once the primary probe solution is removed, the sample is washed with 2x Saline-Sodium Citrate Tween (SSCT) twice. Wash buffer (2xSSCT with 10% ethylene carbonate (EC)) is used for 3 washes and incubated in 60°Cfor 5 minutes each time. Blocking buffer is added and incubated at room temperature for 5 minutes. The sample is then incubated in a solution with 5 nM of the secondary probes in blocking buffer at 37°Cfor an hour. The sample is washed with 2x SSCT twice before using wash buffer to wash 3 times and incubated in 42°Cfor 5 minutes each time. For the first wash, 10 mg/mL Hoechst (H3570; Invitrogen) is diluted 1:1000 in PBS and added to cells. Later, the wash buffer is then removed and replaced with glycerol mounting media and ready for imaging.

### Codetection of protein and mRNA

Prior to mRNA labeling, fixed SW480 cells were blocked with 1% Bovine Serum Albumin (RLBSA50; VWR), 0.1% TWEEN® 20, 1:1,000 Sodium Azide, 0.2 U/ml Protector RNase inhibitor (3335399001; Sigma-Aldrich), and 1 mM DTT in RNAse-free PBS (AM9625; Life Technologies) for 30 min at room temperature. These cells were then washed 3 times with 0.1% TWEEN® 20 in RNAse-free PBS for 5 min each wash at room temperature. Antibody solutions containing 1:1,000 Mouse anti-Tubulin (3873BF; Cell Signaling) and 1:200 Rabbit anti-Vimentin (5741BF; Cell Signaling) in the same blocking buffer were subsequently added to the samples and incubated overnight at 4°C. Following 3 additional washes with 0.1% TWEEN® 20 in RNAse-free PBS for 5 min each at room temperature, antibody solutions containing fluorescently labeled 1:200 Donkey anti-Mouse Alexa-488 (R37114; Fisher Scientific) and 1:200 Donkey anti-Rabbit TRITC (711-025-152; Jackson Laboratories) in the same blocking buffer were added at room temperature for 1 hour. After 3 washes with RNAse-free PBS with 0.1% TWEEN® 20 for 10 min each wash at room temperature, 4% PFA in PBS was added for 15 min at room temperature. These cells were then washed 3 times with 0.1% TWEEN® 20 in PBS at room temperature for 5 min. For mRNA labeling, the previously described methods regarding primary and secondary probe hybridization were utilized.

#### LGC Stellaris ™

LGC Stellaris™RNA FISH probes (Biosearch Technologies, CA, USA) were used, with 48 × 20 mer fluorophore conjugated oligos tiling the length of the target transcript. The POLR2A probe set were supplied as predesigned controls conjugated to Quasar 570 fluorophores. Labeling/staining was carried out as described in the LGC Stellaris™protocol for adherent mammalian cells. The POLR2A probe sets were used at 50 nM.

#### RNAscope™

The FFPE tissue sections were deparaffinized before endogenous peroxidase activity was quenched with hydrogen peroxide. Target retrieval was then performed, followed by protease plus treatment. The fixed cells pretreatment included treatment with hydrogen peroxide and protease 3. The RNAscope™assay was then performed using the RNAscope™Multiplex Fluorescent V2 kit and Akoya Cy5 TSA fluorophore. The positive control (POLR2A) and negative control (dapB) were in C1.

### Microscopy Imaging

Our samples can be imaged with any instrument provided that it has spectral and lifetime acquisition capabilities. Our measurements were taken on two separate instruments; a generic FLIM instrument is depicted in Supplementary Figure 7. For validation of fluorophores and their spectral and lifetime signatures, measurements were taken on a 2-channel ISS Alba5 STED platform. This system is equipped with a pulsed white laser (NKT SuperK EXTREME) system where the excitation wavelength(s) can be selected with an acousto-optic tunable filter (NKT SuperK SELECT). Single photons were detected with avalanche photodiode detectors (Excelitas Technologies) and their arrival times with respect to the stimulating frequency (78MHz) were measured with a FPGA-based electronic board (ISS FastFLIM). Imaging was achieved by fast beam scanning with galvo mirrors and 3D stacks of images were acquired with a z-piezo mount on the objective.

For measurements of multiplexed/combinatorial labeling and detection experiments (Figures 4 and 5), we utilized a Leica SP8 with the Falcon module. This platform employs a white light laser and an acoustic optic beam splitter dichroic, and the Leica hybrid detectors with excitation band selectable by means of a prism. 3D measurements of cells and tissue samples were taken with a 100x plan apochromat oil objective with a numerical aperture of 1.40, yielding images with an x-y resolution of 100 nm and z-spacing of 500 nm.

For epifluorescence measurements (Supplementary Figures 2 and 3), images of labeled mRNA transcripts were taken on an inverted Ti-E using a 100× plan apochromat oil objective with a numerical aperture of 1.40. Samples were illuminated with a Spectra-X (Lumencor) LED light source at the 395 nm, 470 nm, 555 nm and/or 640 nm excitation wavelengths. Images were acquired with an Andor Zyla 4.2 sCMOS camera at 4K resolution with 6.5 µm pixels.

### Image Processing

A custom set of scripts were written in MATLAB to process the acquired image stacks, identify individual transcripts and assign each of them to each gene. After reconstructing the images out of the digital list of photons, the analysis runs in parallel a 3D blob detection pipeline on the intensity image stacks to identify each transcript and on the other a clustering pipeline on the phasor-transformed lifetime/spectral phasor data to detect distinct spectral/lifetime populations. A classifier then assigns pixels as belonging to a particular gene. The whole pipeline is depicted in Supplementary Figure 8.

Briefly, the intensity 3D stacks are run through a blob detection algorithm that was developed in order to identify each transcript. The images can be seen as a 3D space where the transcripts appear as spherically symmetric locations with a radial increase in intensity, namely puncta. The algorithm first computes the low frequency background noise by means of a median filter with a kernel 10 times the size of the expected puncta (the diffraction limit of the instrument, in our case around 250 nm). This low frequency background is subtracted from the high-pass filtered data obtained by convolving by a gaussian filter of the expected size of the puncta. This on one hand enhances the puncta in the image by giving a prominence value at each pixel with respect to the surrounding regions and on the other suppresses noise in the images. A search for local maxima is performed by finding the locations where the gradient goes to zero and the divergence of the gradient is negative. Once the centers in the 3D coordinate space are obtained the size, absolute brightness and prominence of each puncta is measured.

In parallel, the raw photon counts are used to construct the photon arrival time histogram and photon spectral histogram at each pixel. Phasor transforms are applied to each pixel in each image of the 3D stack in order to construct the stacks’ phasor plot. This phasor data is in general a 4-dimensional, each pixel in the intensity image has four additional coordinates; two for the spectral phasor transform plus two for the lifetime phasor transform. The phasor coordinates are clustered using Gaussian Mixture Models^47^. We used an initial experiment tagging house-keeping genes in order to guarantee that all expected populations were present and we trained the Gaussian Mixture Model using this initial experiment. This pretrained model is then applied to the new sets of data in order to classify each pixel into one of the clusters allowing for the presence of empty clusters. The number of clusters N intuitively should be the number of distinct fluorescent probes or different combinations of probes used to tag the sample, but one must allow for additional populations in the sample, e.g., autofluorescent species. For this reason, in the training of the Gaussian Mixture Models we allowed for one additional cluster to account for autofluorescence and noise.

Finally, by computing the mean phasor coordinates of the pixels within each detected puncta, we can compute the phasor position of each puncta and assign a gene label to it by a priori knowing the expected positions of each combination of probes depending on the spectra and lifetime of the probes.

DAPI image stacks are segmented by means of simple thresholding, estimating the threshold value by hard splitting of the histogram of photon counts in the channel. The 3D segmented nuclei are then iteratively grown by convolution by a minimal 3×3×3 kernel. This convolution is applied at each pixel of the edge of the segmented volume until no available space is left in between the segmented volumes. This yields a division of the imaged volume into a number of polyhedra where each face is exactly in the plane bisecting the two closest nuclei edges. This process is analogous to a Voronoi tessellation using the surface of the nuclei instead of points.

### Simulations

In order to test the detection and classification pipeline, we wrote a set of scripts to simulate spectral/lifetime data which provided a ground truth towards detection and accurate classification debugging. This data generation script allows randomly distributing N diffraction-limited transcripts in an arbitrarily big 3-dimensional space, each with a gaussian intensity profile. We simulated our transcript gaussian profile with a X-Y standard deviation of 200 nm and a Z standard deviation of 500 nm, a peak intensity of 1 ± 0.3 (the intensity becomes relevant when simulating background noise). In the simulation run that we used to test the crowding limitations of the system we simulated tagging genes with couples out of a total of 12 fluorescent probes; 4 distinct spectral probes and 3 distinct lifetimes in each, yielding a total of 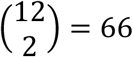possible genes.

We generated the simulated images in a cubic space of 10×10×10 µm, discretized as an image stack of 33 images of 1000 × 1000 pixels (yielding a voxel resolution of 100×100×300 nm). This volume was generated containing increasing densities of transcripts, ranging from a single transcript of each gene (66 transcripts) up to 2,000 transcripts of each gene (132k transcripts) and for each possible value of density a total of 10 iterations each time. These 20k simulated image stack sets were individually processed by our image processing pipeline and the transcript position and labelling obtained by the pipeline was compared to the known ground truth of the generated data. This simulation provided a benchmark of the density limitations of the method but at the same time giving an idea of the underestimation of the number of transcripts as a function of local density. The simulations allowed us to model the estimated number of overlapping transcripts as a function of density.

A similar set of simulations was run by emulating the conditions in the 10-plex experiment (Figure 4) where the genes are tagged with combinations of 2 out of 5 probes. The 20k iterations for different densities allowed to plot the density of the classification obtained after detection compared to the real number of transcripts in the simulations. This simulation was fit to the probabilistic model obtained from calculating the number of transcripts that are not overlapped in space (see next section), from which the true number of puncta was extracted (see Supplementary Figure 4).

### Overlapping Probability

The fraction of puncta that do not overlap with any other puncta depends on the total number of puncta present in the volume of study and the relative proportion between said total volume and the volume of each individual puncta. The following expression is obtained as the product of N-1 times the fraction of available space having removed the volume occupied by one transcript:

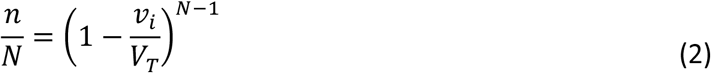

where *n* is the number of isolated puncta, *N* is the total number of puncta, *v*_*i*_ is the volume of each puncta and *V*_*T*_ is the total volume (simulated or scanned). The real number of transcripts N cannot be analytically isolated from the previous equation, but one can graphically obtain it. Due to the fact that the transcripts are sub-diffraction limit, the value of *v*_*i*_ depends on the point spread function of the instrument. Using the detected number of counts *n=*13.5k and the estimated cellular volume 68 kµm^3^, both obtained from the two image stacks shown in Figure 4, we proceeded to estimate the real number of transcripts present in the sample using the previous expression. Assuming an interval of possible volumes for the transcripts between 0.1 and 0.3 µm^3^ we obtained an estimated percentage of overlapping puncta in the interval [2, 6.6]%, which accounts for the number of puncta detected in more than 2 channels in the 10-plex experiment (6%). See Supplementary Figure 4.

### Sequencing Data

Colorectal cancer SW480 cell bulk RNA sequencing (unpublished data) was analyzed with DESeq2. Average expression is then obtained for comparison to the MOSAICA puncta count for each gene. For the human skin melanoma FFPE tissue, the patient sample did not have corresponding sequencing data. RNA sequencing data was obtained from The Cancer Genome Atlas (TCGA), available on the National Cancer Institute (NCI) Genomic Data Commons (GDC) data portal, from 5 human skin melanoma FFPE biopsy thigh punch samples [Entity ID: <u>TCGA-EE-A2GO-06A-11R-A18S-07, TCGA-EE-A20C-06A-11R-A18S-07, TCGA-YG-AA3N-01A-11R-A38C-07, TCGA-DA-A95Z-06A-11R-A37K-07, TCGA-GN-A26C-01A-11R-A18T-07</u>]. The sequencing data were analyzed with HTseq and normalized for sequencing depth and gene length using Fragments Per Kilobase Million (FPKM). The average of the 5 patient samples for each transcript were used for correlation graphs with Spectral-FILM puncta count.

### Statistical Analysis

When comparing distributions of puncta counts, signal-to-noise ratios, and intensity values, Student t-tests were performed for against the probability that the measured distributions belong to distributions with equal means. The reported probability values in the figures are symbolized with (* for p<0.05, ** for p<0.01, *** for p<10^−3^ and **** for p<10^−4^). Pearson correlation coefficient was computed to determine the correlation between the average expression level to the puncta count of each transcript.

### Data Availability

Scripts and algorithms used for the image manipulation, puncta detection and gene classification are available at request. Raw images and data used for figures 3, 4 and 5 are also available at request. Probe sequences used for labeling are included in the supplementary material section.

## Acknowledgement

We thank Dr. Arthur Lander and the UCI Cancer Systems Biology U54 center for their scientific inputs, Dr. Delia Tifrea, Dr. Jeffrey Kim and the UC Irvine Experimental Tissue Resource (ETR) for their help on tissue preparation and pathological characterization, Fairlie Reese for advice about sequencing data processing, Amber Habowski for help processing the bulk-RNA sequencing data, and UCLA’s California NanoSystems Institute and UCI’s Beckman Laser Institute for their microscope support. Some figures were partly created using BioRender.com. This work was funded by a U54CA217378 grant to the UCI Cancer Systems Biology Center (CaSB@UCI), a P30AR075047 NIH/NIAMS P30 Skin Biology Resource Center grant, a P30CA062203 cancer center support grant to the UCI Chao Family Comprehensive Cancer Center, and NIH/NIGMS 1R21GM135493 to P.H. E.G. acknowledge NIH grant NIH P41-GM103540. This work was supported by UCI’s Genomics and High Throughput Facility and Experimental Tissue Resource (ETR) through the Cancer Center Support Grant (P30CA62203). T.V. was supported by NSF GRFP (DGE-1839285). J.G. was supported by a UCI Immunology NIH T32 training grant (AI 060573).

## Author Contribution

T.V., A.V. and J.G. designed, conducted and analyzed the experiments. T.V., A.V., J.G. and W.Z. wrote the manuscript. K.L., Q.X., J.F., J.Z. and C.D. conducted experiments. J.S., L.H., J.W., M.W., A.G., P.H. and E.G. provided technical support and consulted on the study. E.G. and W.Z. designed and directed the project.

## Competing Interests

T.V and J.G are shareholders of Arvetas Biosciences Inc. A.V. and K.L recently joined Arvetas Biosciences Inc as employees. W.Z. is a cofounder of Velox Biosystems Inc., Amberstone Biosciences Inc, and Arvetas Biosciences Inc.

Correspondence and requests for materials should be addressed to E.G. or W.Z.

