## Supplementary Materials for "Spatial transcriptomics using combinatorial fluorescence spectral and lifetime encoding, imaging and analysis"

**Supplementary Material:**

Table S1. List of Fluorophores Used

| **Fluorophore** | Detection Channel | Excitation Max (nm) | Emission Max (nm) | Extinction Coefficient (M-1cm-1) | Quantum Yield | Brightness | Lifetime (ns) |
|---|---|---|---|---|---|---|---|
| ALEXA 647 | 1 | 650 | 668 | 270,000 | 0.33 | 89,100 | 1.04 |
| ATTO 647N | 1 | 646 | 664 | 150,000 | 0.65 | 97,500 | 3.5 |
| ATTO 590 | 2 | 593 | 622 | 120,000 | 0.8 | 96,000 | 3.7 |
| ATTO 565 | 3 | 564 | 590 | 120000 | 0.9 | 108,000 | 4 |
| ATTO 488 | 4 | 500 | 520 | 90,000 | 0.8 | 72,000 | 4.1 |

Table S2. List of Genes Used and their Assigned Fluorophore Combination

| **Gene** | Fluorophore 1 | Fluorophore 2 | No. of Probes per Gene | Present in Channel(s) | Used in Figure(s) |
|---|---|---|---|---|---|
| NCOA3 | ATTO 565 | ATTO 590 | 80 | 2, 3 | 4 |
| POLR2A | ATTO 488 | ALEXA 647 | 80 | 1, 4 | 4 |
| MTOR | ATTO 565 | ATTO 647 | 80 | 1, 3 | 4 |
| MKI67 | ATTO 647 | ALEXA 647 | 80 | 1 | 4 |
| BRCA1 | ATTO 590 | ALEXA 647 | 80 | 1, 2 | 4 |
| NCOA2 | ATTO 590 | ATTO 647 | 80 | 1, 2 | 4 |
| BRCA2 | ATTO 488 | ATTO 565 | 80 | 3, 4 | 4, 5 |
| CENPF | ATTO 488 | ATTO 590 | 80 | 2, 4 | 4, 5 |
| CKAP5 | ATTO 565 | ALEXA 647 | 80 | 1, 3 | 4, 5 |
| NCOA1 | ATTO 488 | ATTO 647 | 80 | 1, 4 | 4, 5 |

### Table S3. List of Sequences Used

| Gene | Probe Number | Codebook | Target Region | Readout 1 | Readout 2 | Full Sequence |
|---|---|---|---|---|---|---|
| MNEONGREEN | 1 | 1 | CTCGACCTTTCTCTTCTTCTTGGGGCT | GGGATGTATTGAAGGAGGAT | GGGATGTATTGAAGGAGGAT | GGGATGTATTGAAGGAGGATTTTGGGATGTATTGAAGGAGGATTTTCTCGACCTTTCTCTTCTTCTTGGGGCT |
| MNEONGREEN | 2 | 1 | GAACTCGCTGCCATCGGCTGTTCTCTT | GGGATGTATTGAAGGAGGAT | GGGATGTATTGAAGGAGGAT | GGGATGTATTGAAGGAGGATTTTGGGATGTATTGAAGGAGGATTTTGAACTCGCTGCCATCGGCTGTTCTCTT |
| MNEONGREEN | 3 | 1 | CTCGCCAGAGCCGGCAGCAGATCCAGC | GGGATGTATTGAAGGAGGAT | GGGATGTATTGAAGGAGGAT | GGGATGTATTGAAGGAGGATTTTGGGATGTATTGAAGGAGGATTTTCTCGCCAGAGCCGGCAGCAGATCCAGC |
| MNEONGREEN | 4 | 1 | GCCCTTGTACAGCTCGTCCATGCCCAT | GGGATGTATTGAAGGAGGAT | GGGATGTATTGAAGGAGGAT | GGGATGTATTGAAGGAGGATTTTGGGATGTATTGAAGGAGGATTTTGCCCTTGTACAGCTCGTCCATGCCCAT |
| MNEONGREEN | 5 | 1 | GTCGGTGAAGGCTTTCTGCCACTCTTT | GGGATGTATTGAAGGAGGAT | GGGATGTATTGAAGGAGGAT | GGGATGTATTGAAGGAGGATTTTGGGATGTATTGAAGGAGGATTTTGTCGGTGAAGGCTTTCTGCCACTCTTT |
| MNEONGREEN | 6 | 1 | GTTCAGTTCTGTCTTGGAGTGCTTCAG | GGGATGTATTGAAGGAGGAT | GGGATGTATTGAAGGAGGAT | GGGATGTATTGAAGGAGGATTTTGGGATGTATTGAAGGAGGATTTTGTTCAGTTCTGTCTTGGAGTGCTTCAG |
| MNEONGREEN | 7 | 1 | GGTCTTGCGGAACACGTACATAGGCTG | GGGATGTATTGAAGGAGGAT | GGGATGTATTGAAGGAGGAT | GGGATGTATTGAAGGAGGATTTTGGGATGTATTGAAGGAGGATTTTGGTCTTGCGGAACACGTACATAGGCTG |
| MNEONGREEN | 8 | 1 | CTTCAGGTAGTTGGCGGCCATGGGTTT | GGGATGTATTGAAGGAGGAT | GGGATGTATTGAAGGAGGAT | GGGATGTATTGAAGGAGGATTTTGGGATGTATTGAAGGAGGATTTTCTTCAGGTAGTTGGCGGCCATGGGTTT |
| MNEONGREEN | 9 | 1 | GAAGGTGTAGGTGGTCCGGGCGGTGCT | GGGATGTATTGAAGGAGGAT | GGGATGTATTGAAGGAGGAT | GGGATGTATTGAAGGAGGATTTTGGGATGTATTGAAGGAGGATTTTGAAGGTGTAGGTGGTCCGGGCGGTGCT |
| MNEONGREEN | 10 | 1 | GTACCGCTTGCCATTGCCGGTGGTGTA | GGGATGTATTGAAGGAGGAT | GGGATGTATTGAAGGAGGAT | GGGATGTATTGAAGGAGGATTTTGGGATGTATTGAAGGAGGATTTTGTACCGCTTGCCATTGCCGGTGGTGTA |
| MNEONGREEN | 11 | 1 | CTCGACCTTTCTCTTCTTCTTGGGGCT | GGGATGTATTGAAGGAGGAT | GGGATGTATTGAAGGAGGAT | GGGATGTATTGAAGGAGGATTTTGGGATGTATTGAAGGAGGATTTTCTCGACCTTTCTCTTCTTCTTGGGGCT |
| MNEONGREEN | 12 | 1 | GAACTCGCTGCCATCGGCTGTTCTCTT | GGGATGTATTGAAGGAGGAT | GGGATGTATTGAAGGAGGAT | GGGATGTATTGAAGGAGGATTTTGGGATGTATTGAAGGAGGATTTTGAACTCGCTGCCATCGGCTGTTCTCTT |
| MNEONGREEN | 13 | 1 | CTCGCCAGAGCCGGCAGCAGATCCAGC | GGGATGTATTGAAGGAGGAT | GGGATGTATTGAAGGAGGAT | GGGATGTATTGAAGGAGGATTTTGGGATGTATTGAAGGAGGATTTTCTCGCCAGAGCCGGCAGCAGATCCAGC |
| MNEONGREEN | 14 | 1 | GCCCTTGTACAGCTCGTCCATGCCCAT | GGGATGTATTGAAGGAGGAT | GGGATGTATTGAAGGAGGAT | GGGATGTATTGAAGGAGGATTTTGGGATGTATTGAAGGAGGATTTTGCCCTTGTACAGCTCGTCCATGCCCAT |
| NCOA3 | 1 | 2 | TTCAATGTCAAACCAGATAAATGTGCG | AGAGTGAGTAGTAGTGGAGT | AGAGTGAGTAGTAGTGGAGT | AGAGTGAGTAGTAGTGGAGTTTTAGAGTGAGTAGTAGTGGAGTTTTTTCAATGTCAAACCAGATAAATGTGCG |
| NCOA3 | 2 | 2 | GTGGTGAATCGAGACGGAAACATTG | TGTGATGGAAGTTAGAGGGT | TGTGATGGAAGTTAGAGGGT | TGTGATGGAAGTTAGAGGGTTTTTGTGATGGAAGTTAGAGGGTTTTGTGGTGAATCGAGACGGAAACATTG |
| NCOA3 | 3 | 2 | CCATACATTTAATTGCCGTATGTTGATGA | AGAGTGAGTAGTAGTGGAGT | AGAGTGAGTAGTAGTGGAGT | AGAGTGAGTAGTAGTGGAGTTTTAGAGTGAGTAGTAGTGGAGTTTTCCATACATTTAATTGCCGTATGTTGATGA |
| NCOA3 | 4 | 2 | ACCACATGATATTCTGGAAGACATAAACG | TGTGATGGAAGTTAGAGGGT | TGTGATGGAAGTTAGAGGGT | TGTGATGGAAGTTAGAGGGTTTTTGTGATGGAAGTTAGAGGGTTTTACCACATGATATTCTGGAAGACATAAACG |
| NCOA3 | 5 | 2 | TGCAGTGCTTTGCCCTGTCTCAGCC | AGAGTGAGTAGTAGTGGAGT | AGAGTGAGTAGTAGTGGAGT | AGAGTGAGTAGTAGTGGAGTTTTAGAGTGAGTAGTAGTGGAGTTTTTGCAGTGCTTTGCCCTGTCTCAGCC |
| NCOA3 | 6 | 2 | CTGGCTTTGAAGATATAATCCGAAGGT | TGTGATGGAAGTTAGAGGGT | TGTGATGGAAGTTAGAGGGT | TGTGATGGAAGTTAGAGGGTTTTTGTGATGGAAGTTAGAGGGTTTTCTGGCTTTGAAGATATAATCCGAAGGT |
| NCOA3 | 7 | 2 | AAAAGCAAACTCTTCCGAAATCCTG | AGAGTGAGTAGTAGTGGAGT | AGAGTGAGTAGTAGTGGAGT | AGAGTGAGTAGTAGTGGAGTTTTAGAGTGAGTAGTAGTGGAGTTTTAAAAGCAAACTCTTCCGAAATCCTG |
| NCOA3 | 8 | 2 | AAATGATCGACATGGCTTTGTCTCA | TGTGATGGAAGTTAGAGGGT | TGTGATGGAAGTTAGAGGGT | TGTGATGGAAGTTAGAGGGTTTTTGTGATGGAAGTTAGAGGGTTTTAAATGATCGACATGGCTTTGTCTCA |
| NCOA3 | 9 | 2 | GATTAGACCACCTATGGCTGGATGC | AGAGTGAGTAGTAGTGGAGT | AGAGTGAGTAGTAGTGGAGT | AGAGTGAGTAGTAGTGGAGTTTTAGAGTGAGTAGTAGTGGAGTTTTGATTAGACCACCTATGGCTGGATGC |
| NCOA3 | 10 | 2 | GTTCGGTAGGCGGCATGAGTATGTC | TGTGATGGAAGTTAGAGGGT | TGTGATGGAAGTTAGAGGGT | TGTGATGGAAGTTAGAGGGTTTTTGTGATGGAAGTTAGAGGGTTTTGTTCGGTAGGCGGCATGAGTATGTC |
| NCOA3 | 11 | 2 | GGCCTATGGCTTGGCAGACCCTAGC | AGAGTGAGTAGTAGTGGAGT | AGAGTGAGTAGTAGTGGAGT | AGAGTGAGTAGTAGTGGAGTTTTAGAGTGAGTAGTAGTGGAGTTTTGGCCTATGGCTTGGCAGACCCTAGC |
| NCOA3 | 12 | 2 | CAGGGCAGATGAGTGGAGCTAGGTA | TGTGATGGAAGTTAGAGGGT | TGTGATGGAAGTTAGAGGGT | TGTGATGGAAGTTAGAGGGTTTTTGTGATGGAAGTTAGAGGGTTTTCAGGGCAGATGAGTGGAGCTAGGTA |
| NCOA3 | 13 | 2 | GGGCCAGGCATGCAATCACCATCTT | AGAGTGAGTAGTAGTGGAGT | AGAGTGAGTAGTAGTGGAGT | AGAGTGAGTAGTAGTGGAGTTTTAGAGTGAGTAGTAGTGGAGTTTTGGGCCAGGCATGCAATCACCATCTT |
| NCOA3 | 14 | 2 | CCAGAACAACAACTATGGGCTCAAC | TGTGATGGAAGTTAGAGGGT | TGTGATGGAAGTTAGAGGGT | TGTGATGGAAGTTAGAGGGTTTTTGTGATGGAAGTTAGAGGGTTTTCCAGAACAACAACTATGGGCTCAAC |
| NCOA3 | 15 | 2 | ACCAGCAGAATATCATGATTTCTCCT | AGAGTGAGTAGTAGTGGAGT | AGAGTGAGTAGTAGTGGAGT | AGAGTGAGTAGTAGTGGAGTTTTAGAGTGAGTAGTAGTGGAGTTTTACCAGCAGAATATCATGATTTCTCCT |
| NCOA3 | 16 | 2 | ATCGTGGGAGTCCAAAGATAGCCTC | TGTGATGGAAGTTAGAGGGT | TGTGATGGAAGTTAGAGGGT | TGTGATGGAAGTTAGAGGGTTTTTGTGATGGAAGTTAGAGGGTTTTATCGTGGGAGTCCAAAGATAGCCTC |
| NCOA3 | 17 | 2 | GCCCTGCAAGCCATCAGTGAAGGTG | AGAGTGAGTAGTAGTGGAGT | AGAGTGAGTAGTAGTGGAGT | AGAGTGAGTAGTAGTGGAGTTTTAGAGTGAGTAGTAGTGGAGTTTTGCCCTGCAAGCCATCAGTGAAGGTG |
| NCOA3 | 18 | 2 | CCCAAATTGGATAACTCTCCCAATATG | TGTGATGGAAGTTAGAGGGT | TGTGATGGAAGTTAGAGGGT | TGTGATGGAAGTTAGAGGGTTTTTGTGATGGAAGTTAGAGGGTTTTCCCAAATTGGATAACTCTCCCAATATG |
| NCOA3 | 19 | 2 | TTCCAAGAGTCCTCTGGGCTTTTAT | AGAGTGAGTAGTAGTGGAGT | AGAGTGAGTAGTAGTGGAGT | AGAGTGAGTAGTAGTGGAGTTTTAGAGTGAGTAGTAGTGGAGTTTTTTCCAAGAGTCCTCTGGGCTTTTAT |
| NCOA3 | 20 | 2 | ACCAAAATCCAGTGGAGAGTTCAAT | TGTGATGGAAGTTAGAGGGT | TGTGATGGAAGTTAGAGGGT | TGTGATGGAAGTTAGAGGGTTTTTGTGATGGAAGTTAGAGGGTTTTACCAAAATCCAGTGGAGAGTTCAAT |
| NCOA3 | 21 | 2 | CAAAGAAAGTAAGGAGAGCAGTGTTG | AGAGTGAGTAGTAGTGGAGT | AGAGTGAGTAGTAGTGGAGT | AGAGTGAGTAGTAGTGGAGTTTTAGAGTGAGTAGTAGTGGAGTTTTCAAAGAAAGTAAGGAGAGCAGTGTTG |
| NCOA3 | 22 | 2 | AATTACTGCAGTTACTTACCTGTTCTTCT | TGTGATGGAAGTTAGAGGGT | TGTGATGGAAGTTAGAGGGT | TGTGATGGAAGTTAGAGGGTTTTTGTGATGGAAGTTAGAGGGTTTTAATTACTGCAGTTACTTACCTGTTCTTCT |
| NCOA3 | 23 | 2 | CTGTTACAAGAGAAGCACCGGATTT | AGAGTGAGTAGTAGTGGAGT | AGAGTGAGTAGTAGTGGAGT | AGAGTGAGTAGTAGTGGAGTTTTAGAGTGAGTAGTAGTGGAGTTTTCTGTTACAAGAGAAGCACCGGATTT |
| NCOA3 | 24 | 2 | CAAGTTGCTGCAGAATGGGAATTCA | TGTGATGGAAGTTAGAGGGT | TGTGATGGAAGTTAGAGGGT | TGTGATGGAAGTTAGAGGGTTTTTGTGATGGAAGTTAGAGGGTTTTCAAGTTGCTGCAGAATGGGAATTCA |
| NCOA3 | 25 | 2 | ACTGGGAAAGACACCAGCAGTATAA | AGAGTGAGTAGTAGTGGAGT | AGAGTGAGTAGTAGTGGAGT | AGAGTGAGTAGTAGTGGAGTTTTAGAGTGAGTAGTAGTGGAGTTTTACTGGGAAAGACACCAGCAGTATAA |
| NCOA3 | 26 | 2 | GGGACGGAAATGTTGTCAAGCAGGA | TGTGATGGAAGTTAGAGGGT | TGTGATGGAAGTTAGAGGGT | TGTGATGGAAGTTAGAGGGTTTTTGTGATGGAAGTTAGAGGGTTTTGGGACGGAAATGTTGTCAAGCAGGA |
| NCOA3 | 27 | 2 | CAAGAGAAAGACCCTAAAATTAAGACAGAG | AGAGTGAGTAGTAGTGGAGT | AGAGTGAGTAGTAGTGGAGT | AGAGTGAGTAGTAGTGGAGTTTTAGAGTGAGTAGTAGTGGAGTTTTCAAGAGAAAGACCCTAAAATTAAGACAGAG |
| NCOA3 | 28 | 2 | ATTCCATATCCTCAAATGGTAGTCATCT | TGTGATGGAAGTTAGAGGGT | TGTGATGGAAGTTAGAGGGT | TGTGATGGAAGTTAGAGGGTTTTTGTGATGGAAGTTAGAGGGTTTTATTCCATATCCTCAAATGGTAGTCATCT |
| NCOA3 | 29 | 2 | CCCATGTTACCAAAGCAACCCATGT | AGAGTGAGTAGTAGTGGAGT | AGAGTGAGTAGTAGTGGAGT | AGAGTGAGTAGTAGTGGAGTTTTAGAGTGAGTAGTAGTGGAGTTTTCCCATGTTACCAAAGCAACCCATGT |
| NCOA3 | 30 | 2 | TGGAACCTATGAATTCAAACTCCATGG | TGTGATGGAAGTTAGAGGGT | TGTGATGGAAGTTAGAGGGT | TGTGATGGAAGTTAGAGGGTTTTTGTGATGGAAGTTAGAGGGTTTTTGGAACCTATGAATTCAAACTCCATGG |
| NCOA3 | 31 | 2 | CCTGCACTGGGTGGCTCTATTCCCA | AGAGTGAGTAGTAGTGGAGT | AGAGTGAGTAGTAGTGGAGT | AGAGTGAGTAGTAGTGGAGTTTTAGAGTGAGTAGTAGTGGAGTTTTCCTGCACTGGGTGGCTCTATTCCCA |
| NCOA3 | 32 | 2 | GCCTCTTCGGTCTAATAGCATACCA | TGTGATGGAAGTTAGAGGGT | TGTGATGGAAGTTAGAGGGT | TGTGATGGAAGTTAGAGGGTTTTTGTGATGGAAGTTAGAGGGTTTTGCCTCTTCGGTCTAATAGCATACCA |
| NCOA3 | 33 | 2 | GCAACACAGATGCCACAGGCCTGGA | AGAGTGAGTAGTAGTGGAGT | AGAGTGAGTAGTAGTGGAGT | AGAGTGAGTAGTAGTGGAGTTTTAGAGTGAGTAGTAGTGGAGTTTTGCAACACAGATGCCACAGGCCTGGA |
| NCOA3 | 34 | 2 | AACAGGATGCTTTCCAAGGCCAAGA | TGTGATGGAAGTTAGAGGGT | TGTGATGGAAGTTAGAGGGT | TGTGATGGAAGTTAGAGGGTTTTTGTGATGGAAGTTAGAGGGTTTTAACAGGATGCTTTCCAAGGCCAAGA |
| NCOA3 | 35 | 2 | ATATGGACAGACATACCCAGCACAG | AGAGTGAGTAGTAGTGGAGT | AGAGTGAGTAGTAGTGGAGT | AGAGTGAGTAGTAGTGGAGTTTTAGAGTGAGTAGTAGTGGAGTTTTATATGGACAGACATACCCAGCACAG |
| NCOA3 | 36 | 2 | CTCCAATGCAAGGAGGCTTTCATCT | TGTGATGGAAGTTAGAGGGT | TGTGATGGAAGTTAGAGGGT | TGTGATGGAAGTTAGAGGGTTTTTGTGATGGAAGTTAGAGGGTTTTCTCCAATGCAAGGAGGCTTTCATCT |
| NCOA3 | 37 | 2 | AGATGAACCAGCAAGGCAATTTTCC | AGAGTGAGTAGTAGTGGAGT | AGAGTGAGTAGTAGTGGAGT | AGAGTGAGTAGTAGTGGAGTTTTAGAGTGAGTAGTAGTGGAGTTTTAGATGAACCAGCAAGGCAATTTTCC |
| NCOA3 | 38 | 2 | CCCAAGCAACTTAGAATGCAGCTTC | TGTGATGGAAGTTAGAGGGT | TGTGATGGAAGTTAGAGGGT | TGTGATGGAAGTTAGAGGGTTTTTGTGATGGAAGTTAGAGGGTTTTCCCAAGCAACTTAGAATGCAGCTTC |
| NCOA3 | 39 | 2 | CACAATGCCACAAGCTCCTCCGCAA | AGAGTGAGTAGTAGTGGAGT | AGAGTGAGTAGTAGTGGAGT | AGAGTGAGTAGTAGTGGAGTTTTAGAGTGAGTAGTAGTGGAGTTTTCACAATGCCACAAGCTCCTCCGCAA |
| NCOA3 | 40 | 2 | CAACCAGATCCAGCCTTTGGTCGAG | TGTGATGGAAGTTAGAGGGT | TGTGATGGAAGTTAGAGGGT | TGTGATGGAAGTTAGAGGGTTTTTGTGATGGAAGTTAGAGGGTTTTCAACCAGATCCAGCCTTTGGTCGAG |
| NCOA3 | 41 | 2 | CAACACCCGCAGGCTGCATCCATCT | AGAGTGAGTAGTAGTGGAGT | AGAGTGAGTAGTAGTGGAGT | AGAGTGAGTAGTAGTGGAGTTTTAGAGTGAGTAGTAGTGGAGTTTTCAACACCCGCAGGCTGCATCCATCT |
| NCOA3 | 42 | 2 | GTCCTCAGAAATGAAGGGCTGGCCA | TGTGATGGAAGTTAGAGGGT | TGTGATGGAAGTTAGAGGGT | TGTGATGGAAGTTAGAGGGTTTTTGTGATGGAAGTTAGAGGGTTTTGTCCTCAGAAATGAAGGGCTGGCCA |
| NCOA3 | 43 | 2 | CATGAATGGCAGCAGTGGTCACATG | AGAGTGAGTAGTAGTGGAGT | AGAGTGAGTAGTAGTGGAGT | AGAGTGAGTAGTAGTGGAGTTTTAGAGTGAGTAGTAGTGGAGTTTTCATGAATGGCAGCAGTGGTCACATG |
| NCOA3 | 44 | 2 | CACCAGGACCTCTTAAGGAAACCAC | TGTGATGGAAGTTAGAGGGT | TGTGATGGAAGTTAGAGGGT | TGTGATGGAAGTTAGAGGGTTTTTGTGATGGAAGTTAGAGGGTTTTCACCAGGACCTCTTAAGGAAACCAC |
| NCOA3 | 45 | 2 | GAATCATTGTTCCAGGCATCCATCT | AGAGTGAGTAGTAGTGGAGT | AGAGTGAGTAGTAGTGGAGT | AGAGTGAGTAGTAGTGGAGTTTTAGAGTGAGTAGTAGTGGAGTTTTGAATCATTGTTCCAGGCATCCATCT |
| NCOA3 | 46 | 2 | AGAAAGGACCAGCTTTGAGCTCCAT | TGTGATGGAAGTTAGAGGGT | TGTGATGGAAGTTAGAGGGT | TGTGATGGAAGTTAGAGGGTTTTTGTGATGGAAGTTAGAGGGTTTTAGAAAGGACCAGCTTTGAGCTCCAT |
| NCOA3 | 47 | 2 | GATTTTAAGCCGAAGGGCAATATCTAC | AGAGTGAGTAGTAGTGGAGT | AGAGTGAGTAGTAGTGGAGT | AGAGTGAGTAGTAGTGGAGTTTTAGAGTGAGTAGTAGTGGAGTTTTGATTTTAAGCCGAAGGGCAATATCTAC |
| NCOA3 | 48 | 2 | CCCTCCTTCTGCTGTGTATCATGGT | TGTGATGGAAGTTAGAGGGT | TGTGATGGAAGTTAGAGGGT | TGTGATGGAAGTTAGAGGGTTTTTGTGATGGAAGTTAGAGGGTTTTCCCTCCTTCTGCTGTGTATCATGGT |
| NCOA3 | 49 | 2 | TTCTGGAGACATGGAGTGTTACTGA | AGAGTGAGTAGTAGTGGAGT | AGAGTGAGTAGTAGTGGAGT | AGAGTGAGTAGTAGTGGAGTTTTAGAGTGAGTAGTAGTGGAGTTTTTTCTGGAGACATGGAGTGTTACTGA |
| NCOA3 | 50 | 2 | TTTTCTGCCTTGCTAGCCAAAATCT | TGTGATGGAAGTTAGAGGGT | TGTGATGGAAGTTAGAGGGT | TGTGATGGAAGTTAGAGGGTTTTTGTGATGGAAGTTAGAGGGTTTTTTTTCTGCCTTGCTAGCCAAAATCT |
| NCOA3 | 51 | 2 | AATATCTGGTTTCTCTAGTTGCAGTATTGG | AGAGTGAGTAGTAGTGGAGT | AGAGTGAGTAGTAGTGGAGT | AGAGTGAGTAGTAGTGGAGTTTTAGAGTGAGTAGTAGTGGAGTTTTAATATCTGGTTTCTCTAGTTGCAGTATTGG |
| NCOA3 | 52 | 2 | AGAGCATAGTCCCAGCCTTCAGGTG | TGTGATGGAAGTTAGAGGGT | TGTGATGGAAGTTAGAGGGT | TGTGATGGAAGTTAGAGGGTTTTTGTGATGGAAGTTAGAGGGTTTTAGAGCATAGTCCCAGCCTTCAGGTG |
| NCOA3 | 53 | 2 | TGATTCTGAATTGTCCTTTACTAATGGTG | AGAGTGAGTAGTAGTGGAGT | AGAGTGAGTAGTAGTGGAGT | AGAGTGAGTAGTAGTGGAGTTTTAGAGTGAGTAGTAGTGGAGTTTTTGATTCTGAATTGTCCTTTACTAATGGTG |
| NCOA3 | 54 | 2 | GTTGCTCTGTCCCTATTATTTGCCC | TGTGATGGAAGTTAGAGGGT | TGTGATGGAAGTTAGAGGGT | TGTGATGGAAGTTAGAGGGTTTTTGTGATGGAAGTTAGAGGGTTTTGTTGCTCTGTCCCTATTATTTGCCC |
| NCOA3 | 55 | 2 | GTCATGGATGTCCAAATGGCTTTGC | AGAGTGAGTAGTAGTGGAGT | AGAGTGAGTAGTAGTGGAGT | AGAGTGAGTAGTAGTGGAGTTTTAGAGTGAGTAGTAGTGGAGTTTTGTCATGGATGTCCAAATGGCTTTGC |
| NCOA3 | 56 | 2 | GTAGCAAACTTTTCACATGCTAATGTG | TGTGATGGAAGTTAGAGGGT | TGTGATGGAAGTTAGAGGGT | TGTGATGGAAGTTAGAGGGTTTTTGTGATGGAAGTTAGAGGGTTTTGTAGCAAACTTTTCACATGCTAATGTG |
| NCOA3 | 57 | 2 | TTTTAAAGTACTGGTGTCACCCTTTGC | AGAGTGAGTAGTAGTGGAGT | AGAGTGAGTAGTAGTGGAGT | AGAGTGAGTAGTAGTGGAGTTTTAGAGTGAGTAGTAGTGGAGTTTTTTTTAAAGTACTGGTGTCACCCTTTGC |
| NCOA3 | 58 | 2 | AAAATAAACTTCTGAAAACCCAAGGCC | TGTGATGGAAGTTAGAGGGT | TGTGATGGAAGTTAGAGGGT | TGTGATGGAAGTTAGAGGGTTTTTGTGATGGAAGTTAGAGGGTTTTAAAATAAACTTCTGAAAACCCAAGGCC |
| NCOA3 | 59 | 2 | GGCTACCAGTGGAAGAGACATCCCT | AGAGTGAGTAGTAGTGGAGT | AGAGTGAGTAGTAGTGGAGT | AGAGTGAGTAGTAGTGGAGTTTTAGAGTGAGTAGTAGTGGAGTTTTGGCTACCAGTGGAAGAGACATCCCT |
| NCOA3 | 60 | 2 | GGGTAGTGCTCCACAGCTTTTCCTT | TGTGATGGAAGTTAGAGGGT | TGTGATGGAAGTTAGAGGGT | TGTGATGGAAGTTAGAGGGTTTTTGTGATGGAAGTTAGAGGGTTTTGGGTAGTGCTCCACAGCTTTTCCTT |
| NCOA3 | 61 | 2 | AAGAGAATCTTTTGGATTTTAAGCAGTCC | AGAGTGAGTAGTAGTGGAGT | AGAGTGAGTAGTAGTGGAGT | AGAGTGAGTAGTAGTGGAGTTTTAGAGTGAGTAGTAGTGGAGTTTTAAGAGAATCTTTTGGATTTTAAGCAGTCC |
| NCOA3 | 62 | 2 | TAATAGCAATTCATGGGCTGTGTGT | TGTGATGGAAGTTAGAGGGT | TGTGATGGAAGTTAGAGGGT | TGTGATGGAAGTTAGAGGGTTTTTGTGATGGAAGTTAGAGGGTTTTTAATAGCAATTCATGGGCTGTGTGT |
| NCOA3 | 63 | 2 | GCTAAGACCTGAATCCCATATTGAGA | AGAGTGAGTAGTAGTGGAGT | AGAGTGAGTAGTAGTGGAGT | AGAGTGAGTAGTAGTGGAGTTTTAGAGTGAGTAGTAGTGGAGTTTTGCTAAGACCTGAATCCCATATTGAGA |
| NCOA3 | 64 | 2 | GAGTAGAATCCTTGGTGTGGTTTCT | TGTGATGGAAGTTAGAGGGT | TGTGATGGAAGTTAGAGGGT | TGTGATGGAAGTTAGAGGGTTTTTGTGATGGAAGTTAGAGGGTTTTGAGTAGAATCCTTGGTGTGGTTTCT |
| NCOA3 | 65 | 2 | TTGACACCTGTTTTAGTCAGTTGGG | AGAGTGAGTAGTAGTGGAGT | AGAGTGAGTAGTAGTGGAGT | AGAGTGAGTAGTAGTGGAGTTTTAGAGTGAGTAGTAGTGGAGTTTTTTGACACCTGTTTTAGTCAGTTGGG |
| NCOA3 | 66 | 2 | TCCCTTTATCTGGGTTAATTCATTTGGT | TGTGATGGAAGTTAGAGGGT | TGTGATGGAAGTTAGAGGGT | TGTGATGGAAGTTAGAGGGTTTTTGTGATGGAAGTTAGAGGGTTTTTCCCTTTATCTGGGTTAATTCATTTGGT |
| NCOA3 | 67 | 2 | CTGTGAATTTCAGAGGTCTCTGCTA | AGAGTGAGTAGTAGTGGAGT | AGAGTGAGTAGTAGTGGAGT | AGAGTGAGTAGTAGTGGAGTTTTAGAGTGAGTAGTAGTGGAGTTTTCTGTGAATTTCAGAGGTCTCTGCTA |
| NCOA3 | 68 | 2 | TGGTATCATTTTCTAGCAATAACTGAGAGC | TGTGATGGAAGTTAGAGGGT | TGTGATGGAAGTTAGAGGGT | TGTGATGGAAGTTAGAGGGTTTTTGTGATGGAAGTTAGAGGGTTTTTGGTATCATTTTCTAGCAATAACTGAGAGC |
| NCOA3 | 69 | 2 | TCTTTGATATGCTTACGAGTAAGTGAATCC | AGAGTGAGTAGTAGTGGAGT | AGAGTGAGTAGTAGTGGAGT | AGAGTGAGTAGTAGTGGAGTTTTAGAGTGAGTAGTAGTGGAGTTTTTCTTTGATATGCTTACGAGTAAGTGAATCC |
| NCOA3 | 70 | 2 | TAAATCCGAAAACTTCCATTGTAGTGG | TGTGATGGAAGTTAGAGGGT | TGTGATGGAAGTTAGAGGGT | TGTGATGGAAGTTAGAGGGTTTTTGTGATGGAAGTTAGAGGGTTTTTAAATCCGAAAACTTCCATTGTAGTGG |
| NCOA3 | 71 | 2 | ACAGAGGAAGAACCAGGTCAGTCTG | AGAGTGAGTAGTAGTGGAGT | AGAGTGAGTAGTAGTGGAGT | AGAGTGAGTAGTAGTGGAGTTTTAGAGTGAGTAGTAGTGGAGTTTTACAGAGGAAGAACCAGGTCAGTCTG |
| NCOA3 | 72 | 2 | TTTTCAGCTCAATTGTATCTGACCCT | TGTGATGGAAGTTAGAGGGT | TGTGATGGAAGTTAGAGGGT | TGTGATGGAAGTTAGAGGGTTTTTGTGATGGAAGTTAGAGGGTTTTTTTTCAGCTCAATTGTATCTGACCCT |
| NCOA3 | 73 | 2 | TTTTAGGTAATTCTTTGTACTCCTGCTGT | AGAGTGAGTAGTAGTGGAGT | AGAGTGAGTAGTAGTGGAGT | AGAGTGAGTAGTAGTGGAGTTTTAGAGTGAGTAGTAGTGGAGTTTTTTTTAGGTAATTCTTTGTACTCCTGCTGT |
| NCOA3 | 74 | 2 | TTTTCAAACCTTGGTATCTGTTGGGT | TGTGATGGAAGTTAGAGGGT | TGTGATGGAAGTTAGAGGGT | TGTGATGGAAGTTAGAGGGTTTTTGTGATGGAAGTTAGAGGGTTTTTTTTCAAACCTTGGTATCTGTTGGGT |
| NCOA3 | 75 | 2 | GGGTAGGGTATTACAGAAGATAATTGGC | AGAGTGAGTAGTAGTGGAGT | AGAGTGAGTAGTAGTGGAGT | AGAGTGAGTAGTAGTGGAGTTTTAGAGTGAGTAGTAGTGGAGTTTTGGGTAGGGTATTACAGAAGATAATTGGC |
| NCOA3 | 76 | 2 | TGTCCTAGAAGTTCTTTGATCCAGAG | TGTGATGGAAGTTAGAGGGT | TGTGATGGAAGTTAGAGGGT | TGTGATGGAAGTTAGAGGGTTTTTGTGATGGAAGTTAGAGGGTTTTTGTCCTAGAAGTTCTTTGATCCAGAG |
| NCOA3 | 77 | 2 | CCAGTTACATGTATGCCTGCCCAGT | AGAGTGAGTAGTAGTGGAGT | AGAGTGAGTAGTAGTGGAGT | AGAGTGAGTAGTAGTGGAGTTTTAGAGTGAGTAGTAGTGGAGTTTTCCAGTTACATGTATGCCTGCCCAGT |
| NCOA3 | 78 | 2 | TTTGCAGAAGCTGTGAGTTTTGTTC | TGTGATGGAAGTTAGAGGGT | TGTGATGGAAGTTAGAGGGT | TGTGATGGAAGTTAGAGGGTTTTTGTGATGGAAGTTAGAGGGTTTTTTTGCAGAAGCTGTGAGTTTTGTTC |
| NCOA3 | 79 | 2 | TTGATTTATTGTTTTCAACTCCAAGGCAC | AGAGTGAGTAGTAGTGGAGT | AGAGTGAGTAGTAGTGGAGT | AGAGTGAGTAGTAGTGGAGTTTTAGAGTGAGTAGTAGTGGAGTTTTTTGATTTATTGTTTTCAACTCCAAGGCAC |
| NCOA3 | 80 | 2 | TTAATAAACGAGCAGGGTGTTTTCTC | TGTGATGGAAGTTAGAGGGT | TGTGATGGAAGTTAGAGGGT | TGTGATGGAAGTTAGAGGGTTTTTGTGATGGAAGTTAGAGGGTTTTTTAATAAACGAGCAGGGTGTTTTCTC |
| POLR2A | 1 | 3 | ACTGGACTCTCTTGATGGTGCGCAGC | AGGGTGTGTTTGTAAAGGGT | AGGGTGTGTTTGTAAAGGGT | AGGGTGTGTTTGTAAAGGGTTTTAGGGTGTGTTTGTAAAGGGTTTTACTGGACTCTCTTGATGGTGCGCAGC |
| POLR2A | 2 | 3 | TTGATGCCACCCTCCGTCACAGACAT | GGGATGTATTGAAGGAGGAT | GGGATGTATTGAAGGAGGAT | GGGATGTATTGAAGGAGGATTTTGGGATGTATTGAAGGAGGATTTTTTGATGCCACCCTCCGTCACAGACAT |
| POLR2A | 3 | 3 | AGGAAGCCCACGTGAAACACAGGCTT | AGGGTGTGTTTGTAAAGGGT | AGGGTGTGTTTGTAAAGGGT | AGGGTGTGTTTGTAAAGGGTTTTAGGGTGTGTTTGTAAAGGGTTTTAGGAAGCCCACGTGAAACACAGGCTT |
| POLR2A | 4 | 3 | CACAAGCAGTTTGGAGCAGAAGAAGC | GGGATGTATTGAAGGAGGAT | GGGATGTATTGAAGGAGGAT | GGGATGTATTGAAGGAGGATTTTGGGATGTATTGAAGGAGGATTTTCACAAGCAGTTTGGAGCAGAAGAAGC |
| POLR2A | 5 | 3 | GGATATCCTTGATCTTTGGGTTGTTAGA | AGGGTGTGTTTGTAAAGGGT | AGGGTGTGTTTGTAAAGGGT | AGGGTGTGTTTGTAAAGGGTTTTAGGGTGTGTTTGTAAAGGGTTTTGGATATCCTTGATCTTTGGGTTGTTAGA |
| POLR2A | 6 | 3 | TTGCAAAGGTCGTAGACATGTGTGAG | GGGATGTATTGAAGGAGGAT | GGGATGTATTGAAGGAGGAT | GGGATGTATTGAAGGAGGATTTTGGGATGTATTGAAGGAGGATTTTTTGCAAAGGTCGTAGACATGTGTGAG |
| POLR2A | 7 | 3 | ACCGAACTTGTTGTCCATCTCCTCCC | AGGGTGTGTTTGTAAAGGGT | AGGGTGTGTTTGTAAAGGGT | AGGGTGTGTTTGTAAAGGGTTTTAGGGTGTGTTTGTAAAGGGTTTTACCGAACTTGTTGTCCATCTCCTCCC |
| POLR2A | 8 | 3 | CATACAGCTCTAGGCCAGAACGCCGG | GGGATGTATTGAAGGAGGAT | GGGATGTATTGAAGGAGGAT | GGGATGTATTGAAGGAGGATTTTGGGATGTATTGAAGGAGGATTTTCATACAGCTCTAGGCCAGAACGCCGG |
| POLR2A | 9 | 3 | AGAGTCCTCATTAACGTGCTTCCATT | AGGGTGTGTTTGTAAAGGGT | AGGGTGTGTTTGTAAAGGGT | AGGGTGTGTTTGTAAAGGGTTTTAGGGTGTGTTTGTAAAGGGTTTTAGAGTCCTCATTAACGTGCTTCCATT |
| POLR2A | 10 | 3 | AGATGCGTTTGAAGATCTCATGCACT | GGGATGTATTGAAGGAGGAT | GGGATGTATTGAAGGAGGAT | GGGATGTATTGAAGGAGGATTTTGGGATGTATTGAAGGAGGATTTTAGATGCGTTTGAAGATCTCATGCACT |
| POLR2A | 11 | 3 | CATGCCCAGCACAAAACACTCCTCAT | AGGGTGTGTTTGTAAAGGGT | AGGGTGTGTTTGTAAAGGGT | AGGGTGTGTTTGTAAAGGGTTTTAGGGTGTGTTTGTAAAGGGTTTTCATGCCCAGCACAAAACACTCCTCAT |
| POLR2A | 12 | 3 | TTCACGATGTCAGCCAGTTTGTGAGT | GGGATGTATTGAAGGAGGAT | GGGATGTATTGAAGGAGGAT | GGGATGTATTGAAGGAGGATTTTGGGATGTATTGAAGGAGGATTTTTTCACGATGTCAGCCAGTTTGTGAGT |
| POLR2A | 13 | 3 | GCTCATTGCGCCGCAGCTGATTGTTG | AGGGTGTGTTTGTAAAGGGT | AGGGTGTGTTTGTAAAGGGT | AGGGTGTGTTTGTAAAGGGTTTTAGGGTGTGTTTGTAAAGGGTTTTGCTCATTGCGCCGCAGCTGATTGTTG |
| POLR2A | 14 | 3 | CAACCGCTGCTTCAGGGACTTGAGGG | GGGATGTATTGAAGGAGGAT | GGGATGTATTGAAGGAGGAT | GGGATGTATTGAAGGAGGATTTTGGGATGTATTGAAGGAGGATTTTCAACCGCTGCTTCAGGGACTTGAGGG |
| POLR2A | 15 | 3 | TCTTTTGCCCATCAGGTTCCCTCGCA | AGGGTGTGTTTGTAAAGGGT | AGGGTGTGTTTGTAAAGGGT | AGGGTGTGTTTGTAAAGGGTTTTAGGGTGTGTTTGTAAAGGGTTTTTCTTTTGCCCATCAGGTTCCCTCGCA |
| POLR2A | 16 | 3 | CAAAGGTCATGTTGGCAGCAATGGAG | GGGATGTATTGAAGGAGGAT | GGGATGTATTGAAGGAGGAT | GGGATGTATTGAAGGAGGATTTTGGGATGTATTGAAGGAGGATTTTCAAAGGTCATGTTGGCAGCAATGGAG |

| | | | | | | |
|---|---|---|---|---|---|---|
| POLR2A | 17 | 3 | TGATGTACTTGGCGCCTGGGTACTGG | AGGGTGTGTTTGTAAAGGGT | AGGGTGTGTTTGTAAAGGGT | AGGGTGTGTTTGTAAAGGGTTTTAGGGTGTGTTTGTAAAGGGTTTTTGATGTACTTGGCGCCTGGGTACTGG |
| POLR2A | 18 | 3 | TAGCCGGTCTGCAGGTGAAGGTCACTG | GGGATGTATTGAAGGAGGAT | GGGATGTATTGAAGGAGGAT | GGGATGTATTGAAGGAGGATTTTGGGATGTATTGAAGGAGGATTTTTAGCCGGTCTGCAGGTGAAGGTCACTG |
| POLR2A | 19 | 3 | CTGCCGGTTGAAGATAACAATGTCCC | AGGGTGTGTTTGTAAAGGGT | AGGGTGTGTTTGTAAAGGGT | AGGGTGTGTTTGTAAAGGGTTTTAGGGTGTGTTTGTAAAGGGTTTTCTGCCGGTTGAAGATAACAATGTCCC |
| POLR2A | 20 | 3 | TAGACCATGGGAGAATGCGGACCCGA | GGGATGTATTGAAGGAGGAT | GGGATGTATTGAAGGAGGAT | GGGATGTATTGAAGGAGGATTTTGGGATGTATTGAAGGAGGATTTTTAGACCATGGGAGAATGCGGACCCGA |
| POLR2A | 21 | 3 | TGGCAGGTGCAAGTTCATCTCATCCC | AGGGTGTGTTTGTAAAGGGT | AGGGTGTGTTTGTAAAGGGT | AGGGTGTGTTTGTAAAGGGTTTTAGGGTGTGTTTGTAAAGGGTTTTTGGCAGGTGCAAGTTCATCTCATCCC |
| POLR2A | 22 | 3 | CAATACCCATGACAGGCCGATTGCTC | GGGATGTATTGAAGGAGGAT | GGGATGTATTGAAGGAGGAT | GGGATGTATTGAAGGAGGATTTTGGGATGTATTGAAGGAGGATTTTCAATACCCATGACAGGCCGATTGCTC |
| POLR2A | 23 | 3 | TTTGCGCACTGCTGTGAGTGTGTCCT | AGGGTGTGTTTGTAAAGGGT | AGGGTGTGTTTGTAAAGGGT | AGGGTGTGTTTGTAAAGGGTTTTAGGGTGTGTTTGTAAAGGGTTTTTTTGCGCACTGCTGTGAGTGTGTCCT |
| POLR2A | 24 | 3 | GGAACATCAGGAGGTTCATCACTTCA | GGGATGTATTGAAGGAGGAT | GGGATGTATTGAAGGAGGAT | GGGATGTATTGAAGGAGGATTTTGGGATGTATTGAAGGAGGATTTTGGAACATCAGGAGGTTCATCACTTCA |
| POLR2A | 25 | 3 | ACAATTGATGTGACCAGGTATGATGAG | AGGGTGTGTTTGTAAAGGGT | AGGGTGTGTTTGTAAAGGGT | AGGGTGTGTTTGTAAAGGGTTTTAGGGTGTGTTTGTAAAGGGTTTTACAATTGATGTGACCAGGTATGATGAG |
| POLR2A | 26 | 3 | GGAGAGATGTGCTTGTAAGGGCCACT | GGGATGTATTGAAGGAGGAT | GGGATGTATTGAAGGAGGAT | GGGATGTATTGAAGGAGGATTTTGGGATGTATTGAAGGAGGATTTTGGAGAGATGTGCTTGTAAGGGCCACT |
| POLR2A | 27 | 3 | CTTACACAGGATGCCCATGATCAGCT | AGGGTGTGTTTGTAAAGGGT | AGGGTGTGTTTGTAAAGGGT | AGGGTGTGTTTGTAAAGGGTTTTAGGGTGTGTTTGTAAAGGGTTTTCTTACACAGGATGCCCATGATCAGCT |
| POLR2A | 28 | 3 | GACAGTCTGAATGTTGGAGTAGAAGA | GGGATGTATTGAAGGAGGAT | GGGATGTATTGAAGGAGGAT | GGGATGTATTGAAGGAGGATTTTGGGATGTATTGAAGGAGGATTTTGACAGTCTGAATGTTGGAGTAGAAGA |
| POLR2A | 29 | 3 | TGACCCTCGATGAGGAGCCAGTTGTT | AGGGTGTGTTTGTAAAGGGT | AGGGTGTGTTTGTAAAGGGT | AGGGTGTGTTTGTAAAGGGTTTTAGGGTGTGTTTGTAAAGGGTTTTTGACCCTCGATGAGGAGCCAGTTGTT |
| POLR2A | 30 | 3 | GCCTTCTTAATAGTGTTCTGAATGTCCTG | GGGATGTATTGAAGGAGGAT | GGGATGTATTGAAGGAGGAT | GGGATGTATTGAAGGAGGATTTTGGGATGTATTGAAGGAGGATTTTGCCTTCTTAATAGTGTTCTGAATGTCCTG |
| POLR2A | 31 | 3 | TCAAACGTCTGCCGCAGAGTGTTCCC | AGGGTGTGTTTGTAAAGGGT | AGGGTGTGTTTGTAAAGGGT | AGGGTGTGTTTGTAAAGGGTTTTAGGGTGTGTTTGTAAAGGGTTTTTCAAACGTCTGCCGCAGAGTGTTCCC |
| POLR2A | 32 | 3 | CTGAGCAGAGGAGCCAGTCTTGTCTC | GGGATGTATTGAAGGAGGAT | GGGATGTATTGAAGGAGGAT | GGGATGTATTGAAGGAGGATTTTGGGATGTATTGAAGGAGGATTTTCTGAGCAGAGGAGCCAGTCTTGTCTC |
| POLR2A | 33 | 3 | ACTTGAAGTTATTGTATTCAGACAGGGA | AGGGTGTGTTTGTAAAGGGT | AGGGTGTGTTTGTAAAGGGT | AGGGTGTGTTTGTAAAGGGTTTTAGGGTGTGTTTGTAAAGGGTTTTACTTGAAGTTATTGTATTCAGACAGGGA |
| POLR2A | 34 | 3 | CAAATGGAATCCGCTTGCCCTCGACG | GGGATGTATTGAAGGAGGAT | GGGATGTATTGAAGGAGGAT | GGGATGTATTGAAGGAGGATTTTGGGATGTATTGAAGGAGGATTTTCAAATGGAATCCGCTTGCCCTCGACG |
| POLR2A | 35 | 3 | GAAGTGAGGCAGAGTCCGGTGCTTGA | AGGGTGTGTTTGTAAAGGGT | AGGGTGTGTTTGTAAAGGGT | AGGGTGTGTTTGTAAAGGGTTTTAGGGTGTGTTTGTAAAGGGTTTTGAAGTGAGGCAGAGTCCGGTGCTTGA |
| POLR2A | 36 | 3 | CGGCTAGGTAGGAGTTCTCCACAAAG | GGGATGTATTGAAGGAGGAT | GGGATGTATTGAAGGAGGAT | GGGATGTATTGAAGGAGGATTTTGGGATGTATTGAAGGAGGATTTTCGGCTAGGTAGGAGTTCTCCACAAAG |
| POLR2A | 37 | 3 | ACTTCACCATCACTGACTCCATGGAC | AGGGTGTGTTTGTAAAGGGT | AGGGTGTGTTTGTAAAGGGT | AGGGTGTGTTTGTAAAGGGTTTTAGGGTGTGTTTGTAAAGGGTTTTACTTCACCATCACTGACTCCATGGAC |
| POLR2A | 38 | 3 | AGGTTCTGGAACTCAACGCTCTCGCC | GGGATGTATTGAAGGAGGAT | GGGATGTATTGAAGGAGGAT | GGGATGTATTGAAGGAGGATTTTGGGATGTATTGAAGGAGGATTTTAGGTTCTGGAACTCAACGCTCTCGCC |
| POLR2A | 39 | 3 | CCTTCACCAGGTCCTCCTGCAGAGTG | AGGGTGTGTTTGTAAAGGGT | AGGGTGTGTTTGTAAAGGGT | AGGGTGTGTTTGTAAAGGGTTTTAGGGTGTGTTTGTAAAGGGTTTTCCTTCACCAGGTCCTCCTGCAGAGTG |
| POLR2A | 40 | 3 | CGCATCCGCTCAAATTCCCGCTCCAA | GGGATGTATTGAAGGAGGAT | GGGATGTATTGAAGGAGGAT | GGGATGTATTGAAGGAGGATTTTGGGATGTATTGAAGGAGGATTTTCGCATCCGCTCAAATTCCCGCTCCAA |
| POLR2A | 41 | 3 | AGATGACCCTGAGCACCTCCCGATCC | AGGGTGTGTTTGTAAAGGGT | AGGGTGTGTTTGTAAAGGGT | AGGGTGTGTTTGTAAAGGGTTTTAGGGTGTGTTTGTAAAGGGTTTTAGATGACCCTGAGCACCTCCCGATCC |
| POLR2A | 42 | 3 | GTGGGTTGATGTGGAAGATTTTCTGA | GGGATGTATTGAAGGAGGAT | GGGATGTATTGAAGGAGGAT | GGGATGTATTGAAGGAGGATTTTGGGATGTATTGAAGGAGGATTTTGTGGGTTGATGTGGAAGATTTTCTGA |
| POLR2A | 43 | 3 | TCACAATCACCAGCTTCTTGCTCAAT | AGGGTGTGTTTGTAAAGGGT | AGGGTGTGTTTGTAAAGGGT | AGGGTGTGTTTGTAAAGGGTTTTAGGGTGTGTTTGTAAAGGGTTTTTCACAATCACCAGCTTCTTGCTCAAT |
| POLR2A | 44 | 3 | ATGTTGAAGAGCAGCGTGGCATTTTC | GGGATGTATTGAAGGAGGAT | GGGATGTATTGAAGGAGGAT | GGGATGTATTGAAGGAGGATTTTGGGATGTATTGAAGGAGGATTTTATGTTGAAGAGCAGCGTGGCATTTTC |
| POLR2A | 45 | 3 | CCACTGAGCCGAAACTCCTCTGCCAT | AGGGTGTGTTTGTAAAGGGT | AGGGTGTGTTTGTAAAGGGT | AGGGTGTGTTTGTAAAGGGTTTTAGGGTGTGTTTGTAAAGGGTTTTCCACTGAGCCGAAACTCCTCTGCCAT |
| POLR2A | 46 | 3 | TCTGGGTGGCAGGTTCTCCAAGGGAC | GGGATGTATTGAAGGAGGAT | GGGATGTATTGAAGGAGGAT | GGGATGTATTGAAGGAGGATTTTGGGATGTATTGAAGGAGGATTTTTCTGGGTGGCAGGTTCTCCAAGGGAC |
| POLR2A | 47 | 3 | ACCAGCATAGTGGAAGGTATTCAAGG | AGGGTGTGTTTGTAAAGGGT | AGGGTGTGTTTGTAAAGGGT | AGGGTGTGTTTGTAAAGGGTTTTAGGGTGTGTTTGTAAAGGGTTTTACCAGCATAGTGGAAGGTATTCAAGG |
| POLR2A | 48 | 3 | AAATGTTGATGAGCTCCTTAAGTCGG | GGGATGTATTGAAGGAGGAT | GGGATGTATTGAAGGAGGAT | GGGATGTATTGAAGGAGGATTTTGGGATGTATTGAAGGAGGATTTTAAATGTTGATGAGCTCCTTAAGTCGG |
| POLR2A | 49 | 3 | AGTAAGCGAAGGAGTCTTTGGCTTCT | AGGGTGTGTTTGTAAAGGGT | AGGGTGTGTTTGTAAAGGGT | AGGGTGTGTTTGTAAAGGGTTTTAGGGTGTGTTTGTAAAGGGTTTTAGTAAGCGAAGGAGTCTTTGGCTTCT |
| POLR2A | 50 | 3 | TATGCTCCAGACGGCACAGAATATCC | GGGATGTATTGAAGGAGGAT | GGGATGTATTGAAGGAGGAT | GGGATGTATTGAAGGAGGATTTTGGGATGTATTGAAGGAGGATTTTTATGCTCCAGACGGCACAGAATATCC |
| POLR2A | 51 | 3 | TGTGTTGGCAGTCACCTTCCTCAACG | AGGGTGTGTTTGTAAAGGGT | AGGGTGTGTTTGTAAAGGGT | AGGGTGTGTTTGTAAAGGGTTTTAGGGTGTGTTTGTAAAGGGTTTTTGTGTTGGCAGTCACCTTCCTCAACG |
| POLR2A | 52 | 3 | CAAAGTCAGGCATTTCATAGTAGACATTC | GGGATGTATTGAAGGAGGAT | GGGATGTATTGAAGGAGGAT | GGGATGTATTGAAGGAGGATTTTGGGATGTATTGAAGGAGGATTTTCAAAGTCAGGCATTTCATAGTAGACATTC |
| POLR2A | 53 | 3 | AGTCATGTGCTTCCGATCCAGCTCCA | AGGGTGTGTTTGTAAAGGGT | AGGGTGTGTTTGTAAAGGGT | AGGGTGTGTTTGTAAAGGGTTTTAGGGTGTGTTTGTAAAGGGTTTTAGTCATGTGCTTCCGATCCAGCTCCA |
| POLR2A | 54 | 3 | GCATTGTCATCATTAAAGATGCAGTTCA | GGGATGTATTGAAGGAGGAT | GGGATGTATTGAAGGAGGAT | GGGATGTATTGAAGGAGGATTTTGGGATGTATTGAAGGAGGATTTTGCATTGTCATCATTAAAGATGCAGTTCA |
| POLR2A | 55 | 3 | TGATGCGAATACGGAGCACCAGCTTC | AGGGTGTGTTTGTAAAGGGT | AGGGTGTGTTTGTAAAGGGT | AGGGTGTGTTTGTAAAGGGTTTTAGGGTGTGTTTGTAAAGGGTTTTTGATGCGAATACGGAGCACCAGCTTC |
| POLR2A | 56 | 3 | TCATCCATCTTGTCCACCACCTCTTC | GGGATGTATTGAAGGAGGAT | GGGATGTATTGAAGGAGGAT | GGGATGTATTGAAGGAGGATTTTGGGATGTATTGAAGGAGGATTTTTCATCCATCTTGTCCACCACCTCTTC |
| POLR2A | 57 | 3 | ATGTTGGACTCGATGCAGCGCAGGAA | AGGGTGTGTTTGTAAAGGGT | AGGGTGTGTTTGTAAAGGGT | AGGGTGTGTTTGTAAAGGGTTTTAGGGTGTGTTTGTAAAGGGTTTTATGTTGGACTCGATGCAGCGCAGGAA |
| POLR2A | 58 | 3 | CCACTCCTGCAGGGCCTTGAATTCCC | GGGATGTATTGAAGGAGGAT | GGGATGTATTGAAGGAGGAT | GGGATGTATTGAAGGAGGATTTTGGGATGTATTGAAGGAGGATTTTCCACTCCTGCAGGGCCTTGAATTCCC |
| POLR2A | 59 | 3 | CTCACTCAGCACCCGCATCAAGCTCA | AGGGTGTGTTTGTAAAGGGT | AGGGTGTGTTTGTAAAGGGT | AGGGTGTGTTTGTAAAGGGTTTTAGGGTGTGTTTGTAAAGGGTTTTCTCACTCAGCACCCGCATCAAGCTCA |
| POLR2A | 60 | 3 | AGATGACGTGGTACAGCTCCCGCTCC | GGGATGTATTGAAGGAGGAT | GGGATGTATTGAAGGAGGAT | GGGATGTATTGAAGGAGGATTTTGGGATGTATTGAAGGAGGATTTTAGATGACGTGGTACAGCTCCCGCTCC |
| POLR2A | 61 | 3 | TCGGTAATTGACATAGGAGCCATCAA | AGGGTGTGTTTGTAAAGGGT | AGGGTGTGTTTGTAAAGGGT | AGGGTGTGTTTGTAAAGGGTTTTAGGGTGTGTTTGTAAAGGGTTTTTCGGTAATTGACATAGGAGCCATCAA |
| POLR2A | 62 | 3 | TTCATGAGTGGTCCTGTGTCCTGGCG | GGGATGTATTGAAGGAGGAT | GGGATGTATTGAAGGAGGAT | GGGATGTATTGAAGGAGGATTTTGGGATGTATTGAAGGAGGATTTTTTCATGAGTGGTCCTGTGTCCTGGCG |
| POLR2A | 63 | 3 | GCCCAGCATGATATTCTCAGAGACCC | AGGGTGTGTTTGTAAAGGGT | AGGGTGTGTTTGTAAAGGGT | AGGGTGTGTTTGTAAAGGGTTTTAGGGTGTGTTTGTAAAGGGTTTTGCCCAGCATGATATTCTCAGAGACCC |
| POLR2A | 64 | 3 | CCCTGGTTCCAAGGTGTCATGGCAGG | GGGATGTATTGAAGGAGGAT | GGGATGTATTGAAGGAGGAT | GGGATGTATTGAAGGAGGATTTTGGGATGTATTGAAGGAGGATTTTCCCTGGTTCCAAGGTGTCATGGCAGG |
| POLR2A | 65 | 3 | GTGACGTTGGCGAGTAGCTGGGAGAC | AGGGTGTGTTTGTAAAGGGT | AGGGTGTGTTTGTAAAGGGT | AGGGTGTGTTTGTAAAGGGTTTTAGGGTGTGTTTGTAAAGGGTTTTGTGACGTTGGCGAGTAGCTGGGAGAC |
| POLR2A | 66 | 3 | GGGTGATGTGGGACTATAGTTGGGAC | GGGATGTATTGAAGGAGGAT | GGGATGTATTGAAGGAGGAT | GGGATGTATTGAAGGAGGATTTTGGGATGTATTGAAGGAGGATTTTGGGTGATGTGGGACTATAGTTGGGAC |
| POLR2A | 67 | 3 | GAGTAGCTGGGTGACGTTGGCGAATA | AGGGTGTGTTTGTAAAGGGT | AGGGTGTGTTTGTAAAGGGT | AGGGTGTGTTTGTAAAGGGTTTTAGGGTGTGTTTGTAAAGGGTTTTGAGTAGCTGGGTGACGTTGGCGAATA |
| POLR2A | 68 | 3 | CGAGTAGCTGGGAGATGTCGGCGAGT | GGGATGTATTGAAGGAGGAT | GGGATGTATTGAAGGAGGAT | GGGATGTATTGAAGGAGGATTTTGGGATGTATTGAAGGAGGATTTTCGAGTAGCTGGGAGATGTCGGCGAGT |
| POLR2A | 69 | 3 | GAAGTGGGAGAATAGCTGGGTGAAGT | AGGGTGTGTTTGTAAAGGGT | AGGGTGTGTTTGTAAAGGGT | AGGGTGTGTTTGTAAAGGGTTTTAGGGTGTGTTTGTAAAGGGTTTTGAAGTGGGAGAATAGCTGGGTGAAGT |
| POLR2A | 70 | 3 | TAGTTCGGGCTGGTGGGTGAGTAACT | GGGATGTATTGAAGGAGGAT | GGGATGTATTGAAGGAGGAT | GGGATGTATTGAAGGAGGATTTTGGGATGTATTGAAGGAGGATTTTTAGTTCGGGCTGGTGGGTGAGTAACT |
| POLR2A | 71 | 3 | GATGTCGGGCTGTAGCTGGGTGATGT | AGGGTGTGTTTGTAAAGGGT | AGGGTGTGTTTGTAAAGGGT | AGGGTGTGTTTGTAAAGGGTTTTAGGGTGTGTTTGTAAAGGGTTTTGATGTCGGGCTGTAGCTGGGTGATGT |
| POLR2A | 72 | 3 | TGGGCTGTAGTTAGGGCTGGTAGGTG | GGGATGTATTGAAGGAGGAT | GGGATGTATTGAAGGAGGAT | GGGATGTATTGAAGGAGGATTTTGGGATGTATTGAAGGAGGATTTTTGGGCTGTAGTTAGGGCTGGTAGGTG |
| POLR2A | 73 | 3 | GGTGATGTTGGAGAGTAGCTTGGAGA | AGGGTGTGTTTGTAAAGGGT | AGGGTGTGTTTGTAAAGGGT | AGGGTGTGTTTGTAAAGGGTTTTAGGGTGTGTTTGTAAAGGGTTTTGGTGATGTTGGAGAGTAGCTTGGAGA |
| POLR2A | 74 | 3 | TGGGACTGTAAGAAGGACTGGTTGGG | GGGATGTATTGAAGGAGGAT | GGGATGTATTGAAGGAGGAT | GGGATGTATTGAAGGAGGATTTTGGGATGTATTGAAGGAGGATTTTTGGGACTGTAAGAAGGACTGGTTGGG |
| POLR2A | 75 | 3 | GAATAGGTGGGACTGGTAGGCGAGTA | AGGGTGTGTTTGTAAAGGGT | AGGGTGTGTTTGTAAAGGGT | AGGGTGTGTTTGTAAAGGGTTTTAGGGTGTGTTTGTAAAGGGTTTTGAATAGGTGGGACTGGTAGGCGAGTA |
| POLR2A | 76 | 3 | GATAGCCGGGCTTGTGAGACTGTAGG | GGGATGTATTGAAGGAGGAT | GGGATGTATTGAAGGAGGAT | GGGATGTATTGAAGGAGGATTTTGGGATGTATTGAAGGAGGATTTTGATAGCCGGGCTTGTGAGACTGTAGG |
| POLR2A | 77 | 3 | CAGTTCTCCTCGTCACTGTCATCCGG | AGGGTGTGTTTGTAAAGGGT | AGGGTGTGTTTGTAAAGGGT | AGGGTGTGTTTGTAAAGGGTTTTAGGGTGTGTTTGTAAAGGGTTTTCAGTTCTCCTCGTCACTGTCATCCGG |
| MTOR | 1 | 4 | ACTGGCAAACTGCTGCAGGACGCTC | AGAGTGAGTAGTAGTGGAGT | AGAGTGAGTAGTAGTGGAGT | AGAGTGAGTAGTAGTGGAGTTTTAGAGTGAGTAGTAGTGGAGTTTTACTGGCAAACTGCTGCAGGACGCTC |
| MTOR | 2 | 4 | TGGTTTCCTCATTCCGGCTCTTTAG | ATAGGAAATGGTGGTAGTGT | ATAGGAAATGGTGGTAGTGT | ATAGGAAATGGTGGTAGTGTTTTATAGGAAATGGTGGTAGTGTTTTTGGTTTCCTCATTCCGGCTCTTTAG |
| MTOR | 3 | 4 | CCATGACAACTGGGTCATTGGAGGG | AGAGTGAGTAGTAGTGGAGT | AGAGTGAGTAGTAGTGGAGT | AGAGTGAGTAGTAGTGGAGTTTTAGAGTGAGTAGTAGTGGAGTTTTCCATGACAACTGGGTCATTGGAGGG |
| MTOR | 4 | 4 | GGCAAGACGGCCAATGGCCTTGGAT | ATAGGAAATGGTGGTAGTGT | ATAGGAAATGGTGGTAGTGT | ATAGGAAATGGTGGTAGTGTTTTATAGGAAATGGTGGTAGTGTTTTGGCAAGACGGCCAATGGCCTTGGAT |
| MTOR | 5 | 4 | TTCCAGGGCTCGCTTCACCTCAAAT | AGAGTGAGTAGTAGTGGAGT | AGAGTGAGTAGTAGTGGAGT | AGAGTGAGTAGTAGTGGAGTTTTAGAGTGAGTAGTAGTGGAGTTTTTTCCAGGGCTCGCTTCACCTCAAAT |
| MTOR | 6 | 4 | TTGCTGGAAGAAGAAGGTAGGGACG | ATAGGAAATGGTGGTAGTGT | ATAGGAAATGGTGGTAGTGT | ATAGGAAATGGTGGTAGTGTTTTATAGGAAATGGTGGTAGTGTTTTTTGCTGGAAGAAGAAGGTAGGGACG |
| MTOR | 7 | 4 | ATCAGACAGGCACGAAGGGCGGCTA | AGAGTGAGTAGTAGTGGAGT | AGAGTGAGTAGTAGTGGAGT | AGAGTGAGTAGTAGTGGAGTTTTAGAGTGAGTAGTAGTGGAGTTTTATCAGACAGGCACGAAGGGCGGCTA |
| MTOR | 8 | 4 | CTCCTTCGGCTCACGCTGGGTTGTG | ATAGGAAATGGTGGTAGTGT | ATAGGAAATGGTGGTAGTGT | ATAGGAAATGGTGGTAGTGTTTTATAGGAAATGGTGGTAGTGTTTTCTCCTTCGGCTCACGCTGGGTTGTG |
| MTOR | 9 | 4 | AGGCTCCATGGATCCGATCATCCCG | AGAGTGAGTAGTAGTGGAGT | AGAGTGAGTAGTAGTGGAGT | AGAGTGAGTAGTAGTGGAGTTTTAGAGTGAGTAGTAGTGGAGTTTTAGGCTCCATGGATCCGATCATCCCG |
| MTOR | 10 | 4 | ATTCGGACCAGCTCGTTAAGGATCA | ATAGGAAATGGTGGTAGTGT | ATAGGAAATGGTGGTAGTGT | ATAGGAAATGGTGGTAGTGTTTTATAGGAAATGGTGGTAGTGTTTTATTCGGACCAGCTCGTTAAGGATCA |
| MTOR | 11 | 4 | TCTTTGCAGTACTTGTCGTGTACCA | AGAGTGAGTAGTAGTGGAGT | AGAGTGAGTAGTAGTGGAGT | AGAGTGAGTAGTAGTGGAGTTTTAGAGTGAGTAGTAGTGGAGTTTTTCTTTGCAGTACTTGTCGTGTACCA |
| MTOR | 12 | 4 | ACGAGGTTTTGTTCCGAAGCCCATG | ATAGGAAATGGTGGTAGTGT | ATAGGAAATGGTGGTAGTGT | ATAGGAAATGGTGGTAGTGTTTTATAGGAAATGGTGGTAGTGTTTTACGAGGTTTTGTTCCGAAGCCCATG |
| MTOR | 13 | 4 | TGAGGCCTTGGTGAGAGCTGTACCC | AGAGTGAGTAGTAGTGGAGT | AGAGTGAGTAGTAGTGGAGT | AGAGTGAGTAGTAGTGGAGTTTTAGAGTGAGTAGTAGTGGAGTTTTTGAGGCCTTGGTGAGAGCTGTACCC |
| MTOR | 14 | 4 | CCAGGGTGGACTTAGCTGGACTGGG | ATAGGAAATGGTGGTAGTGT | ATAGGAAATGGTGGTAGTGT | ATAGGAAATGGTGGTAGTGTTTTATAGGAAATGGTGGTAGTGTTTTCCAGGGTGGACTTAGCTGGACTGGG |
| MTOR | 15 | 4 | TCTTGACACAGCTTAGGACATGGTT | AGAGTGAGTAGTAGTGGAGT | AGAGTGAGTAGTAGTGGAGT | AGAGTGAGTAGTAGTGGAGTTTTAGAGTGAGTAGTAGTGGAGTTTTTCTTGACACAGCTTAGGACATGGTT |
| MTOR | 16 | 4 | ACCTCACAGCCACAGAAAGTAGCCC | ATAGGAAATGGTGGTAGTGT | ATAGGAAATGGTGGTAGTGT | ATAGGAAATGGTGGTAGTGTTTTATAGGAAATGGTGGTAGTGTTTTACCTCACAGCCACAGAAAGTAGCCC |
| MTOR | 17 | 4 | CTTTAGCTGTGGAATCTGACGGCTC | AGAGTGAGTAGTAGTGGAGT | AGAGTGAGTAGTAGTGGAGT | AGAGTGAGTAGTAGTGGAGTTTTAGAGTGAGTAGTAGTGGAGTTTTCTTTAGCTGTGGAATCTGACGGCTC |
| MTOR | 18 | 4 | CATTTTCAGTAGCCCATCTTGAATGTC | ATAGGAAATGGTGGTAGTGT | ATAGGAAATGGTGGTAGTGT | ATAGGAAATGGTGGTAGTGTTTTATAGGAAATGGTGGTAGTGTTTTCATTTTCAGTAGCCCATCTTGAATGTC |
| MTOR | 19 | 4 | ATTCAAAGCTGCCAAGCGTTCGGAG | AGAGTGAGTAGTAGTGGAGT | AGAGTGAGTAGTAGTGGAGT | AGAGTGAGTAGTAGTGGAGTTTTAGAGTGAGTAGTAGTGGAGTTTTATTCAAAGCTGCCAAGCGTTCGGAG |
| MTOR | 20 | 4 | GTTCAGGAAATGATCCGCACAGTGG | ATAGGAAATGGTGGTAGTGT | ATAGGAAATGGTGGTAGTGT | ATAGGAAATGGTGGTAGTGTTTTATAGGAAATGGTGGTAGTGTTTTGTTCAGGAAATGATCCGCACAGTGG |
| MTOR | 21 | 4 | GAGGTGGATGGAGGGTGTGAGCAGG | AGAGTGAGTAGTAGTGGAGT | AGAGTGAGTAGTAGTGGAGT | AGAGTGAGTAGTAGTGGAGTTTTAGAGTGAGTAGTAGTGGAGTTTTGAGGTGGATGGAGGGTGTGAGCAGG |
| MTOR | 22 | 4 | GGCTAACCACATGAGCATGGCCACT | ATAGGAAATGGTGGTAGTGT | ATAGGAAATGGTGGTAGTGT | ATAGGAAATGGTGGTAGTGTTTTATAGGAAATGGTGGTAGTGTTTTGGCTAACCACATGAGCATGGCCACT |
| MTOR | 23 | 4 | GTGCATCAAAGCGCTCGTCCAGGGA | AGAGTGAGTAGTAGTGGAGT | AGAGTGAGTAGTAGTGGAGT | AGAGTGAGTAGTAGTGGAGTTTTAGAGTGAGTAGTAGTGGAGTTTTGTGCATCAAAGCGCTCGTCCAGGGA |
| MTOR | 24 | 4 | CAAGGCCTGCAAGTTCTCCGCCTGG | ATAGGAAATGGTGGTAGTGT | ATAGGAAATGGTGGTAGTGT | ATAGGAAATGGTGGTAGTGTTTTATAGGAAATGGTGGTAGTGTTTTCAAGGCCTGCAAGTTCTCCGCCTGG |
| MTOR | 25 | 4 | CATGCGGGCACTCTGCTCTTTGATT | AGAGTGAGTAGTAGTGGAGT | AGAGTGAGTAGTAGTGGAGT | AGAGTGAGTAGTAGTGGAGTTTTAGAGTGAGTAGTAGTGGAGTTTTCATGCGGGCACTCTGCTCTTTGATT |
| MTOR | 26 | 4 | GGGCTCTACTACATAGCCAGTGCTG | ATAGGAAATGGTGGTAGTGT | ATAGGAAATGGTGGTAGTGT | ATAGGAAATGGTGGTAGTGTTTTATAGGAAATGGTGGTAGTGTTTTGGGCTCTACTACATAGCCAGTGCTG |
| MTOR | 27 | 4 | TCTGCTCAGTCTTCAGAAAATTCAGTA | AGAGTGAGTAGTAGTGGAGT | AGAGTGAGTAGTAGTGGAGT | AGAGTGAGTAGTAGTGGAGTTTTAGAGTGAGTAGTAGTGGAGTTTTTCTGCTCAGTCTTCAGAAAATTCAGTA |
| MTOR | 28 | 4 | TTGTGCTTGTAAGGATCCAAAGCCC | ATAGGAAATGGTGGTAGTGT | ATAGGAAATGGTGGTAGTGT | ATAGGAAATGGTGGTAGTGTTTTATAGGAAATGGTGGTAGTGTTTTTTGTGCTTGTAAGGATCCAAAGCCC |
| MTOR | 29 | 4 | CTGACAGGCTGACAGCAGAGGCATC | AGAGTGAGTAGTAGTGGAGT | AGAGTGAGTAGTAGTGGAGT | AGAGTGAGTAGTAGTGGAGTTTTAGAGTGAGTAGTAGTGGAGTTTTCTGACAGGCTGACAGCAGAGGCATC |
| MTOR | 30 | 4 | GTTGACCAGCATTTCACTAGTGCTA | ATAGGAAATGGTGGTAGTGT | ATAGGAAATGGTGGTAGTGT | ATAGGAAATGGTGGTAGTGTTTTATAGGAAATGGTGGTAGTGTTTTGTTGACCAGCATTTCACTAGTGCTA |
| MTOR | 31 | 4 | ATCAGGGCCACCATGGACACAGCTG | AGAGTGAGTAGTAGTGGAGT | AGAGTGAGTAGTAGTGGAGT | AGAGTGAGTAGTAGTGGAGTTTTAGAGTGAGTAGTAGTGGAGTTTTATCAGGGCCACCATGGACACAGCTG |
| MTOR | 32 | 4 | AGAGAGTGACTGGTCTCGGAAGATC | ATAGGAAATGGTGGTAGTGT | ATAGGAAATGGTGGTAGTGT | ATAGGAAATGGTGGTAGTGTTTTATAGGAAATGGTGGTAGTGTTTTAGAGAGTGACTGGTCTCGGAAGATC |
| MTOR | 33 | 4 | AGTCCCAGGGACTTGAAGATGAAGG | AGAGTGAGTAGTAGTGGAGT | AGAGTGAGTAGTAGTGGAGT | AGAGTGAGTAGTAGTGGAGTTTTAGAGTGAGTAGTAGTGGAGTTTTAGTCCCAGGGACTTGAAGATGAAGG |
| MTOR | 34 | 4 | TTAAGGAACGTGGGCATGACCTGGG | ATAGGAAATGGTGGTAGTGT | ATAGGAAATGGTGGTAGTGT | ATAGGAAATGGTGGTAGTGTTTTATAGGAAATGGTGGTAGTGTTTTTTAAGGAACGTGGGCATGACCTGGG |
| MTOR | 35 | 4 | GCTCAATGAGAAGAATGATCGTGCT | AGAGTGAGTAGTAGTGGAGT | AGAGTGAGTAGTAGTGGAGT | AGAGTGAGTAGTAGTGGAGTTTTAGAGTGAGTAGTAGTGGAGTTTTGCTCAATGAGAAGAATGATCGTGCT |
| MTOR | 36 | 4 | GGGCAGGTAGAGCTTAAATTCACCC | ATAGGAAATGGTGGTAGTGT | ATAGGAAATGGTGGTAGTGT | ATAGGAAATGGTGGTAGTGTTTTATAGGAAATGGTGGTAGTGTTTTGGGCAGGTAGAGCTTAAATTCACCC |
| MTOR | 37 | 4 | GTCAGGCGGTCCACAGTCTCTAGCG | AGAGTGAGTAGTAGTGGAGT | AGAGTGAGTAGTAGTGGAGT | AGAGTGAGTAGTAGTGGAGTTTTAGAGTGAGTAGTAGTGGAGTTTTGTCAGGCGGTCCACAGTCTCTAGCG |
| MTOR | 38 | 4 | GGCATAGTCAGTGAAATCCAGGGAC | ATAGGAAATGGTGGTAGTGT | ATAGGAAATGGTGGTAGTGT | ATAGGAAATGGTGGTAGTGTTTTATAGGAAATGGTGGTAGTGTTTTGGCATAGTCAGTGAAATCCAGGGAC |
| MTOR | 39 | 4 | TCCATGGCTGTGGAGCGCAGTTCTG | AGAGTGAGTAGTAGTGGAGT | AGAGTGAGTAGTAGTGGAGT | AGAGTGAGTAGTAGTGGAGTTTTAGAGTGAGTAGTAGTGGAGTTTTTCCATGGCTGTGGAGCGCAGTTCTG |
| MTOR | 40 | 4 | TCAAAGGATCCTCCTCTTCATCAGC | ATAGGAAATGGTGGTAGTGT | ATAGGAAATGGTGGTAGTGT | ATAGGAAATGGTGGTAGTGTTTTATAGGAAATGGTGGTAGTGTTTTTCAAAGGATCCTCCTCTTCATCAGC |
| MTOR | 41 | 4 | ACTGGTCCACTAGCCAATGCATCCC | AGAGTGAGTAGTAGTGGAGT | AGAGTGAGTAGTAGTGGAGT | AGAGTGAGTAGTAGTGGAGTTTTAGAGTGAGTAGTAGTGGAGTTTTACTGGTCCACTAGCCAATGCATCCC |
| MTOR | 42 | 4 | GTGCAGTTTCTTCATGGGTCCTGTT | ATAGGAAATGGTGGTAGTGT | ATAGGAAATGGTGGTAGTGT | ATAGGAAATGGTGGTAGTGTTTTATAGGAAATGGTGGTAGTGTTTTGTGCAGTTTCTTCATGGGTCCTGTT |
| MTOR | 43 | 4 | GGCTCAGCCGTCTCAGCCATTCCAG | AGAGTGAGTAGTAGTGGAGT | AGAGTGAGTAGTAGTGGAGT | AGAGTGAGTAGTAGTGGAGTTTTAGAGTGAGTAGTAGTGGAGTTTTGGCTCAGCCGTCTCAGCCATTCCAG |
| MTOR | 44 | 4 | GGCGATGATGAGTCCTTCAGCAGCT | ATAGGAAATGGTGGTAGTGT | ATAGGAAATGGTGGTAGTGT | ATAGGAAATGGTGGTAGTGTTTTATAGGAAATGGTGGTAGTGTTTTGGCGATGATGAGTCCTTCAGCAGCT |
| MTOR | 45 | 4 | GAGGGCCAACTCGATGCTTCTGATG | AGAGTGAGTAGTAGTGGAGT | AGAGTGAGTAGTAGTGGAGT | AGAGTGAGTAGTAGTGGAGTTTTAGAGTGAGTAGTAGTGGAGTTTTGAGGGCCAACTCGATGCTTCTGATG |
| MTOR | 46 | 4 | GTGTGACTTCAGCGATGTCTTGTGA | ATAGGAAATGGTGGTAGTGT | ATAGGAAATGGTGGTAGTGT | ATAGGAAATGGTGGTAGTGTTTTATAGGAAATGGTGGTAGTGTTTTGTGTGACTTCAGCGATGTCTTGTGA |
| MTOR | 47 | 4 | GGCAGCTCTCTCACCCAGCAGAACA | AGAGTGAGTAGTAGTGGAGT | AGAGTGAGTAGTAGTGGAGT | AGAGTGAGTAGTAGTGGAGTTTTAGAGTGAGTAGTAGTGGAGTTTTGGCAGCTCTCTCACCCAGCAGAACA |
| MTOR | 48 | 4 | GTAGTGCTTTGGCATATGCTCGGCA | ATAGGAAATGGTGGTAGTGT | ATAGGAAATGGTGGTAGTGT | ATAGGAAATGGTGGTAGTGTTTTATAGGAAATGGTGGTAGTGTTTTGTAGTGCTTTGGCATATGCTCGGCA |
| MTOR | 49 | 4 | GTCATAGGCCACAAGGGCATCCTCC | AGAGTGAGTAGTAGTGGAGT | AGAGTGAGTAGTAGTGGAGT | AGAGTGAGTAGTAGTGGAGTTTTAGAGTGAGTAGTAGTGGAGTTTTGTCATAGGCCACAAGGGCATCCTCC |
| MTOR | 50 | 4 | GGTCGTCCTTGTTGGTGTCCATTTT | ATAGGAAATGGTGGTAGTGT | ATAGGAAATGGTGGTAGTGT | ATAGGAAATGGTGGTAGTGTTTTATAGGAAATGGTGGTAGTGTTTTGGTCGTCCTTGTTGGTGTCCATTTT |
| MTOR | 51 | 4 | TGCCAAGGAGAAGAGGTCCTGATGC | AGAGTGAGTAGTAGTGGAGT | AGAGTGAGTAGTAGTGGAGT | AGAGTGAGTAGTAGTGGAGTTTTAGAGTGAGTAGTAGTGGAGTTTTTGCCAAGGAGAAGAGGTCCTGATGC |

| | | | | | | |
|---|---|---|---|---|---|---|
| MTOR | 52 | 4 | AGTTTGTACTGGATAACCTCCTCCA | ATAGGAAATGGTGGTAGTGT | ATAGGAAATGGTGGTAGTGT | ATAGGAAATGGTGGTAGTGTTTTATAGGAAATGGTGGTAGTGTTTTAGTTTGTACTGGATAACCTCCTCCA |
| MTOR | 53 | 4 | ATGTCTTCATGAGGGCTGACCACAA | AGAGTGAGTAGTAGTGGAGT | AGAGTGAGTAGTAGTGGAGT | AGAGTGAGTAGTAGTGGAGTTTTAGAGTGAGTAGTAGTGGAGTTTTATGTCTTCATGAGGGCTGACCACAA |
| MTOR | 54 | 4 | CAGGCTTGCATACTTGAGCCAGGTT | ATAGGAAATGGTGGTAGTGT | ATAGGAAATGGTGGTAGTGT | ATAGGAAATGGTGGTAGTGTTTTATAGGAAATGGTGGTAGTGTTTTCAGGCTTGCATACTTGAGCCAGGTT |
| MTOR | 55 | 4 | TTGCGGGCACTCTTCCACATGTTTT | AGAGTGAGTAGTAGTGGAGT | AGAGTGAGTAGTAGTGGAGT | AGAGTGAGTAGTAGTGGAGTTTTAGAGTGAGTAGTAGTGGAGTTTTTTGCGGGCACTCTTCCACATGTTTT |
| MTOR | 56 | 4 | TCATTGATGCCCTGTAGATTCAGCT | ATAGGAAATGGTGGTAGTGT | ATAGGAAATGGTGGTAGTGT | ATAGGAAATGGTGGTAGTGTTTTATAGGAAATGGTGGTAGTGTTTTTCATTGATGCCCTGTAGATTCAGCT |
| MTOR | 57 | 4 | TTTCTTCTTCTCATCGCGGGCTTGG | AGAGTGAGTAGTAGTGGAGT | AGAGTGAGTAGTAGTGGAGT | AGAGTGAGTAGTAGTGGAGTTTTAGAGTGAGTAGTAGTGGAGTTTTTTTCTTCTTCTCATCGCGGGCTTGG |
| MTOR | 58 | 4 | GGCAGTGGTGGCGTTGGTGATGTTG | ATAGGAAATGGTGGTAGTGT | ATAGGAAATGGTGGTAGTGT | ATAGGAAATGGTGGTAGTGTTTTATAGGAAATGGTGGTAGTGTTTTGGCAGTGGTGGCGTTGGTGATGTTG |
| MTOR | 59 | 4 | CCAAGGGTCTGGGCGTATCAATTCT | AGAGTGAGTAGTAGTGGAGT | AGAGTGAGTAGTAGTGGAGT | AGAGTGAGTAGTAGTGGAGTTTTAGAGTGAGTAGTAGTGGAGTTTTCCAAGGGTCTGGGCGTATCAATTCT |
| MTOR | 60 | 4 | GTGAGAAGCTGGTGAATGAGACGTC | ATAGGAAATGGTGGTAGTGT | ATAGGAAATGGTGGTAGTGT | ATAGGAAATGGTGGTAGTGTTTTATAGGAAATGGTGGTAGTGTTTTGTGAGAAGCTGGTGAATGAGACGTC |
| MTOR | 61 | 4 | TCAGAATCTTGTTGGCTGCATTGTG | AGAGTGAGTAGTAGTGGAGT | AGAGTGAGTAGTAGTGGAGT | AGAGTGAGTAGTAGTGGAGTTTTAGAGTGAGTAGTAGTGGAGTTTTTCAGAATCTTGTTGGCTGCATTGTG |
| MTOR | 62 | 4 | AGGGTGTTGCTGTGCTCACACATGT | ATAGGAAATGGTGGTAGTGT | ATAGGAAATGGTGGTAGTGT | ATAGGAAATGGTGGTAGTGTTTTATAGGAAATGGTGGTAGTGTTTTAGGGTGTTGCTGTGCTCACACATGT |
| MTOR | 63 | 4 | TCTTCCAGGCCTTCATGCCACATCT | AGAGTGAGTAGTAGTGGAGT | AGAGTGAGTAGTAGTGGAGT | AGAGTGAGTAGTAGTGGAGTTTTAGAGTGAGTAGTAGTGGAGTTTTTCTTCCAGGCCTTCATGCCACATCT |
| MTOR | 64 | 4 | AACATGCCTTTCACGTTCCTTTCCC | ATAGGAAATGGTGGTAGTGT | ATAGGAAATGGTGGTAGTGT | ATAGGAAATGGTGGTAGTGTTTTATAGGAAATGGTGGTAGTGTTTTAACATGCCTTTCACGTTCCTTTCCC |
| MTOR | 65 | 4 | CATTAAATCTCGACCATAGGCCTGA | AGAGTGAGTAGTAGTGGAGT | AGAGTGAGTAGTAGTGGAGT | AGAGTGAGTAGTAGTGGAGTTTTAGAGTGAGTAGTAGTGGAGTTTTCATTAAATCTCGACCATAGGCCTGA |
| MTOR | 66 | 4 | TGTACTTCCTGCACCACTCTTGGGC | ATAGGAAATGGTGGTAGTGT | ATAGGAAATGGTGGTAGTGT | ATAGGAAATGGTGGTAGTGTTTTATAGGAAATGGTGGTAGTGTTTTTGTACTTCCTGCACCACTCTTGGGC |
| MTOR | 67 | 4 | GAGGCAGCTGCTTTGAGATTCGTCG | AGAGTGAGTAGTAGTGGAGT | AGAGTGAGTAGTAGTGGAGT | AGAGTGAGTAGTAGTGGAGTTTTAGAGTGAGTAGTAGTGGAGTTTTGAGGCAGCTGCTTTGAGATTCGTCG |
| MTOR | 68 | 4 | ACATATTGCAGCTCTAAGGATGTGA | ATAGGAAATGGTGGTAGTGT | ATAGGAAATGGTGGTAGTGT | ATAGGAAATGGTGGTAGTGTTTTATAGGAAATGGTGGTAGTGTTTTACATATTGCAGCTCTAAGGATGTGA |
| MTOR | 69 | 4 | CATATGTTCCTGGCACAGCCAATTC | AGAGTGAGTAGTAGTGGAGT | AGAGTGAGTAGTAGTGGAGT | AGAGTGAGTAGTAGTGGAGTTTTAGAGTGAGTAGTAGTGGAGTTTTCATATGTTCCTGGCACAGCCAATTC |
| MTOR | 70 | 4 | TGAATGCGAATGATTGGCTGGTTGG | ATAGGAAATGGTGGTAGTGT | ATAGGAAATGGTGGTAGTGT | ATAGGAAATGGTGGTAGTGTTTTATAGGAAATGGTGGTAGTGTTTTTGAATGCGAATGATTGGCTGGTTGG |
| MTOR | 71 | 4 | TTTAGAAGGAAAACAAACTCATGTCCG | AGAGTGAGTAGTAGTGGAGT | AGAGTGAGTAGTAGTGGAGT | AGAGTGAGTAGTAGTGGAGTTTTAGAGTGAGTAGTAGTGGAGTTTTTTTAGAAGGAAAACAAACTCATGTCCG |
| MTOR | 72 | 4 | CTCATCCTGGCGCAGATCTTCATGG | ATAGGAAATGGTGGTAGTGT | ATAGGAAATGGTGGTAGTGT | ATAGGAAATGGTGGTAGTGTTTTATAGGAAATGGTGGTAGTGTTTTCTCATCCTGGCGCAGATCTTCATGG |
| MTOR | 73 | 4 | GATGTTGGGTCATTGGCCAGAAGGG | AGAGTGAGTAGTAGTGGAGT | AGAGTGAGTAGTAGTGGAGT | AGAGTGAGTAGTAGTGGAGTTTTAGAGTGAGTAGTAGTGGAGTTTTGATGTTGGGTCATTGGCCAGAAGGG |
| MTOR | 74 | 4 | GCCCGAGTTGGTCGATAAAGGGATG | ATAGGAAATGGTGGTAGTGT | ATAGGAAATGGTGGTAGTGT | ATAGGAAATGGTGGTAGTGTTTTATAGGAAATGGTGGTAGTGTTTTGCCCGAGTTGGTCGATAAAGGGATG |
| MTOR | 75 | 4 | CAACATGATGCGATGCTCGATGTTG | AGAGTGAGTAGTAGTGGAGT | AGAGTGAGTAGTAGTGGAGT | AGAGTGAGTAGTAGTGGAGTTTTAGAGTGAGTAGTAGTGGAGTTTTCAACATGATGCGATGCTCGATGTTG |
| MTOR | 76 | 4 | TTTTCAGCCACAGCAGCTTGGCCAG | ATAGGAAATGGTGGTAGTGT | ATAGGAAATGGTGGTAGTGT | ATAGGAAATGGTGGTAGTGTTTTATAGGAAATGGTGGTAGTGTTTTTTTTCAGCCACAGCAGCTTGGCCAG |
| MTOR | 77 | 4 | ACCTCCATCACTGTGTGGCATGTGA | AGAGTGAGTAGTAGTGGAGT | AGAGTGAGTAGTAGTGGAGT | AGAGTGAGTAGTAGTGGAGTTTTAGAGTGAGTAGTAGTGGAGTTTTACCTCCATCACTGTGTGGCATGTGA |
| MTOR | 78 | 4 | GGTCATAGACAAAGGCTTCCAGCAC | ATAGGAAATGGTGGTAGTGT | ATAGGAAATGGTGGTAGTGT | ATAGGAAATGGTGGTAGTGTTTTATAGGAAATGGTGGTAGTGTTTTGGTCATAGACAAAGGCTTCCAGCAC |
| MTOR | 79 | 4 | AGTGTCATCATGAGAGAAGTCCCGA | AGAGTGAGTAGTAGTGGAGT | AGAGTGAGTAGTAGTGGAGT | AGAGTGAGTAGTAGTGGAGTTTTAGAGTGAGTAGTAGTGGAGTTTTAGTGTCATCATGAGAGAAGTCCCGA |
| MTOR | 80 | 4 | TTTTCATGGGATGTCGCTTGTTTGA | ATAGGAAATGGTGGTAGTGT | ATAGGAAATGGTGGTAGTGT | ATAGGAAATGGTGGTAGTGTTTTATAGGAAATGGTGGTAGTGTTTTTTTTCATGGGATGTCGCTTGTTTGA |
| MKI67 | 1 | 5 | TTTGTTGCTGTGAATATTGTACAGGG | ATAGGAAATGGTGGTAGTGT | ATAGGAAATGGTGGTAGTGT | ATAGGAAATGGTGGTAGTGTTTTATAGGAAATGGTGGTAGTGTTTTTTTGTTGCTGTGAATATTGTACAGGG |
| MKI67 | 2 | 5 | AGGAGGCAAGTTTTCATCAAATAGTTC | GGGATGTATTGAAGGAGGAT | GGGATGTATTGAAGGAGGAT | GGGATGTATTGAAGGAGGATTTTGGGATGTATTGAAGGAGGATTTTAGGAGGCAAGTTTTCATCAAATAGTTC |
| MKI67 | 3 | 5 | AGGACCTACGGCGTTGATCACTGGCA | ATAGGAAATGGTGGTAGTGT | ATAGGAAATGGTGGTAGTGT | ATAGGAAATGGTGGTAGTGTTTTATAGGAAATGGTGGTAGTGTTTTAGGACCTACGGCGTTGATCACTGGCA |
| MKI67 | 4 | 5 | CTTTCTCCCTCCTCTCTTAGGAACCT | GGGATGTATTGAAGGAGGAT | GGGATGTATTGAAGGAGGAT | GGGATGTATTGAAGGAGGATTTTGGGATGTATTGAAGGAGGATTTTCTTTCTCCCTCCTCTCTTAGGAACCT |
| MKI67 | 5 | 5 | AGGGTCGAGCAGGCACATGTACTTTT | ATAGGAAATGGTGGTAGTGT | ATAGGAAATGGTGGTAGTGT | ATAGGAAATGGTGGTAGTGTTTTATAGGAAATGGTGGTAGTGTTTTAGGGTCGAGCAGGCACATGTACTTTT |
| MKI67 | 6 | 5 | AGGGCTTGCAGAGCATTTATCAGATG | GGGATGTATTGAAGGAGGAT | GGGATGTATTGAAGGAGGAT | GGGATGTATTGAAGGAGGATTTTGGGATGTATTGAAGGAGGATTTTAGGGCTTGCAGAGCATTTATCAGATG |
| MKI67 | 7 | 5 | GACCTGTTTGCAGTGGATACTGTTTT | ATAGGAAATGGTGGTAGTGT | ATAGGAAATGGTGGTAGTGT | ATAGGAAATGGTGGTAGTGTTTTATAGGAAATGGTGGTAGTGTTTTGACCTGTTTGCAGTGGATACTGTTTT |
| MKI67 | 8 | 5 | CTGTATATTCCTGAACTCTGTAGACCTT | GGGATGTATTGAAGGAGGAT | GGGATGTATTGAAGGAGGAT | GGGATGTATTGAAGGAGGATTTTGGGATGTATTGAAGGAGGATTTTCTGTATATTCCTGAACTCTGTAGACCTT |
| MKI67 | 9 | 5 | AGTAGTGTTGCCTTCTGACCTCTTTT | ATAGGAAATGGTGGTAGTGT | ATAGGAAATGGTGGTAGTGT | ATAGGAAATGGTGGTAGTGTTTTATAGGAAATGGTGGTAGTGTTTTAGTAGTGTTGCCTTCTGACCTCTTTT |
| MKI67 | 10 | 5 | TATCAGGCAAGCTCTTGAGGTCTGTC | GGGATGTATTGAAGGAGGAT | GGGATGTATTGAAGGAGGAT | GGGATGTATTGAAGGAGGATTTTGGGATGTATTGAAGGAGGATTTTTATCAGGCAAGCTCTTGAGGTCTGTC |
| MKI67 | 11 | 5 | TTTTGCCTTTCTCACTCTTTGGTGCC | ATAGGAAATGGTGGTAGTGT | ATAGGAAATGGTGGTAGTGT | ATAGGAAATGGTGGTAGTGTTTTATAGGAAATGGTGGTAGTGTTTTTTTTGCCTTTCTCACTCTTTGGTGCC |
| MKI67 | 12 | 5 | TTGTAATGACTGGCAGGGCATTTTAG | GGGATGTATTGAAGGAGGAT | GGGATGTATTGAAGGAGGAT | GGGATGTATTGAAGGAGGATTTTGGGATGTATTGAAGGAGGATTTTTTGTAATGACTGGCAGGGCATTTTAG |
| MKI67 | 13 | 5 | CCTGACGTCCGTGTGAACTTGCCGAC | ATAGGAAATGGTGGTAGTGT | ATAGGAAATGGTGGTAGTGT | ATAGGAAATGGTGGTAGTGTTTTATAGGAAATGGTGGTAGTGTTTTCCTGACGTCCGTGTGAACTTGCCGAC |
| MKI67 | 14 | 5 | ATCTCCTGCTGGCTCTCTGTGCGTGT | GGGATGTATTGAAGGAGGAT | GGGATGTATTGAAGGAGGAT | GGGATGTATTGAAGGAGGATTTTGGGATGTATTGAAGGAGGATTTTATCTCCTGCTGGCTCTCTGTGCGTGT |
| MKI67 | 15 | 5 | TTCATTCCAGTTACACGGGCTGCTGG | ATAGGAAATGGTGGTAGTGT | ATAGGAAATGGTGGTAGTGT | ATAGGAAATGGTGGTAGTGTTTTATAGGAAATGGTGGTAGTGTTTTTTCATTCCAGTTACACGGGCTGCTGG |
| MKI67 | 16 | 5 | CCTCTTCCTTAGGCGTTCTTGGCCAC | GGGATGTATTGAAGGAGGAT | GGGATGTATTGAAGGAGGAT | GGGATGTATTGAAGGAGGATTTTGGGATGTATTGAAGGAGGATTTTCCTCTTCCTTAGGCGTTCTTGGCCAC |
| MKI67 | 17 | 5 | AGTCTGTAGCTGTCTTTTGCTGCCAG | ATAGGAAATGGTGGTAGTGT | ATAGGAAATGGTGGTAGTGT | ATAGGAAATGGTGGTAGTGTTTTATAGGAAATGGTGGTAGTGTTTTAGTCTGTAGCTGTCTTTTGCTGCCAG |
| MKI67 | 18 | 5 | AGCCACTAATTCCTCGGTGTGACCAG | GGGATGTATTGAAGGAGGAT | GGGATGTATTGAAGGAGGAT | GGGATGTATTGAAGGAGGATTTTGGGATGTATTGAAGGAGGATTTTAGCCACTAATTCCTCGGTGTGACCAG |
| MKI67 | 19 | 5 | ATGGCATTAGATTCCTGCACGCTAAG | ATAGGAAATGGTGGTAGTGT | ATAGGAAATGGTGGTAGTGT | ATAGGAAATGGTGGTAGTGTTTTATAGGAAATGGTGGTAGTGTTTTATGGCATTAGATTCCTGCACGCTAAG |
| MKI67 | 20 | 5 | CCACTGCTTCTTCAGTATGACCAGGG | GGGATGTATTGAAGGAGGAT | GGGATGTATTGAAGGAGGAT | GGGATGTATTGAAGGAGGATTTTGGGATGTATTGAAGGAGGATTTTCCACTGCTTCTTCAGTATGACCAGGG |
| MKI67 | 21 | 5 | TCTTGGGCTGCCTTCTTGTGCTTGTT | ATAGGAAATGGTGGTAGTGT | ATAGGAAATGGTGGTAGTGT | ATAGGAAATGGTGGTAGTGTTTTATAGGAAATGGTGGTAGTGTTTTTCTTGGGCTGCCTTCTTGTGCTTGTT |
| MKI67 | 22 | 5 | CTTCTGTACGTCCCTTTTCTCCAAAG | GGGATGTATTGAAGGAGGAT | GGGATGTATTGAAGGAGGAT | GGGATGTATTGAAGGAGGATTTTGGGATGTATTGAAGGAGGATTTTCTTCTGTACGTCCCTTTTCTCCAAAG |
| MKI67 | 23 | 5 | TCAGTGCATACTGGTGTCTGGAAGAG | ATAGGAAATGGTGGTAGTGT | ATAGGAAATGGTGGTAGTGT | ATAGGAAATGGTGGTAGTGTTTTATAGGAAATGGTGGTAGTGTTTTTCAGTGCATACTGGTGTCTGGAAGAG |
| MKI67 | 24 | 5 | TGGTAGTTTTCTCGTGAGTCGTGGGC | GGGATGTATTGAAGGAGGAT | GGGATGTATTGAAGGAGGAT | GGGATGTATTGAAGGAGGATTTTGGGATGTATTGAAGGAGGATTTTTGGTAGTTTTCTCGTGAGTCGTGGGC |
| MKI67 | 25 | 5 | TTTCCTGAGTGCGAAGAATTCTTCTT | ATAGGAAATGGTGGTAGTGT | ATAGGAAATGGTGGTAGTGT | ATAGGAAATGGTGGTAGTGTTTTATAGGAAATGGTGGTAGTGTTTTTTTCCTGAGTGCGAAGAATTCTTCTT |
| MKI67 | 26 | 5 | TCTCACCACTTACTGCTGGTTTGGGT | GGGATGTATTGAAGGAGGAT | GGGATGTATTGAAGGAGGAT | GGGATGTATTGAAGGAGGATTTTGGGATGTATTGAAGGAGGATTTTTCTCACCACTTACTGCTGGTTTGGGT |
| MKI67 | 27 | 5 | TTTGCAGGCTACTTTGGCAGTTTTAT | ATAGGAAATGGTGGTAGTGT | ATAGGAAATGGTGGTAGTGT | ATAGGAAATGGTGGTAGTGTTTTATAGGAAATGGTGGTAGTGTTTTTTTGCAGGCTACTTTGGCAGTTTTAT |
| MKI67 | 28 | 5 | CCGTCGCTTGGAGCTTGCTGGGTTTT | GGGATGTATTGAAGGAGGAT | GGGATGTATTGAAGGAGGAT | GGGATGTATTGAAGGAGGATTTTGGGATGTATTGAAGGAGGATTTTCCGTCGCTTGGAGCTTGCTGGGTTTT |
| MKI67 | 29 | 5 | GCTTTCATGCTCTTACCATCTCCTGT | ATAGGAAATGGTGGTAGTGT | ATAGGAAATGGTGGTAGTGT | ATAGGAAATGGTGGTAGTGTTTTATAGGAAATGGTGGTAGTGTTTTGCTTTCATGCTCTTACCATCTCCTGT |
| MKI67 | 30 | 5 | AGACTTTCCCTTAGGAGTTCTCAGCT | GGGATGTATTGAAGGAGGAT | GGGATGTATTGAAGGAGGAT | GGGATGTATTGAAGGAGGATTTTGGGATGTATTGAAGGAGGATTTTAGACTTTCCCTTAGGAGTTCTCAGCT |
| MKI67 | 31 | 5 | AGCTCTGTAGGATACTTTGGTAGTTTT | ATAGGAAATGGTGGTAGTGT | ATAGGAAATGGTGGTAGTGT | ATAGGAAATGGTGGTAGTGTTTTATAGGAAATGGTGGTAGTGTTTTAGCTCTGTAGGATACTTTGGTAGTTTT |
| MKI67 | 32 | 5 | TGGCGTTTGTTTCCTAAATGCTAAAA | GGGATGTATTGAAGGAGGAT | GGGATGTATTGAAGGAGGAT | GGGATGTATTGAAGGAGGATTTTGGGATGTATTGAAGGAGGATTTTTGGCGTTTGTTTCCTAAATGCTAAAA |
| MKI67 | 33 | 5 | CTGAAGCCAGTCAGTTCTTCTAGAGC | ATAGGAAATGGTGGTAGTGT | ATAGGAAATGGTGGTAGTGT | ATAGGAAATGGTGGTAGTGTTTTATAGGAAATGGTGGTAGTGTTTTCTGAAGCCAGTCAGTTCTTCTAGAGC |
| MKI67 | 34 | 5 | TATCAGTGCATGGTGTCTGGAAAAGC | GGGATGTATTGAAGGAGGAT | GGGATGTATTGAAGGAGGAT | GGGATGTATTGAAGGAGGATTTTGGGATGTATTGAAGGAGGATTTTTATCAGTGCATGGTGTCTGGAAAAGC |
| MKI67 | 35 | 5 | TGGGCCGTTGCTTTGTGTTTGTTGGG | ATAGGAAATGGTGGTAGTGT | ATAGGAAATGGTGGTAGTGT | ATAGGAAATGGTGGTAGTGTTTTATAGGAAATGGTGGTAGTGTTTTTGGGCCGTTGCTTTGTGTTTGTTGGG |
| MKI67 | 36 | 5 | CTCTACGTCTGCTTTCTTGAGGCTTC | GGGATGTATTGAAGGAGGAT | GGGATGTATTGAAGGAGGAT | GGGATGTATTGAAGGAGGATTTTGGGATGTATTGAAGGAGGATTTTCTCTACGTCTGCTTTCTTGAGGCTTC |
| MKI67 | 37 | 5 | TGAGTCGTTGCTTGGAGCTTGTTGGG | ATAGGAAATGGTGGTAGTGT | ATAGGAAATGGTGGTAGTGT | ATAGGAAATGGTGGTAGTGTTTTATAGGAAATGGTGGTAGTGTTTTTGAGTCGTTGCTTGGAGCTTGTTGGG |
| MKI67 | 38 | 5 | AAACGCTTTGATGCTCTTTCCATCTC | GGGATGTATTGAAGGAGGAT | GGGATGTATTGAAGGAGGAT | GGGATGTATTGAAGGAGGATTTTGGGATGTATTGAAGGAGGATTTTAAACGCTTTGATGCTCTTTCCATCTC |
| MKI67 | 39 | 5 | TTGGGCCTCTTCCTTAGGTGTTCTTG | ATAGGAAATGGTGGTAGTGT | ATAGGAAATGGTGGTAGTGT | ATAGGAAATGGTGGTAGTGTTTTATAGGAAATGGTGGTAGTGTTTTTTGGGCCTCTTCCTTAGGTGTTCTTG |
| MKI67 | 40 | 5 | CCTCAGTGTGGTCTGGTGTCTGGAAG | GGGATGTATTGAAGGAGGAT | GGGATGTATTGAAGGAGGAT | GGGATGTATTGAAGGAGGATTTTGGGATGTATTGAAGGAGGATTTTCCTCAGTGTGGTCTGGTGTCTGGAAG |
| MKI67 | 41 | 5 | TGTGTGGTCTGTGTGAGCTGCTTCAG | ATAGGAAATGGTGGTAGTGT | ATAGGAAATGGTGGTAGTGT | ATAGGAAATGGTGGTAGTGTTTTATAGGAAATGGTGGTAGTGTTTTTGTGTGGTCTGTGTGAGCTGCTTCAG |
| MKI67 | 42 | 5 | TATCCTCATCTCCTGGTACTTTGTCT | GGGATGTATTGAAGGAGGAT | GGGATGTATTGAAGGAGGAT | GGGATGTATTGAAGGAGGATTTTGGGATGTATTGAAGGAGGATTTTTATCCTCATCTCCTGGTACTTTGTCT |
| MKI67 | 43 | 5 | GAGTTCTTGGCTGCCTCTTGCTACCA | ATAGGAAATGGTGGTAGTGT | ATAGGAAATGGTGGTAGTGT | ATAGGAAATGGTGGTAGTGTTTTATAGGAAATGGTGGTAGTGTTTTGAGTTCTTGGCTGCCTCTTGCTACCA |
| MKI67 | 44 | 5 | TTTCAAGCCAGCCAAGTCTTCTAGGG | GGGATGTATTGAAGGAGGAT | GGGATGTATTGAAGGAGGAT | GGGATGTATTGAAGGAGGATTTTGGGATGTATTGAAGGAGGATTTTTTTCAAGCCAGCCAAGTCTTCTAGGG |
| MKI67 | 45 | 5 | TGGTAGTTTTCTCATGAGTCGTGGGC | ATAGGAAATGGTGGTAGTGT | ATAGGAAATGGTGGTAGTGT | ATAGGAAATGGTGGTAGTGTTTTATAGGAAATGGTGGTAGTGTTTTTGGTAGTTTTCTCATGAGTCGTGGGC |
| MKI67 | 46 | 5 | TGGACTGTGGCTTGAAGATTGTTGGG | GGGATGTATTGAAGGAGGAT | GGGATGTATTGAAGGAGGAT | GGGATGTATTGAAGGAGGATTTTGGGATGTATTGAAGGAGGATTTTTGGACTGTGGCTTGAAGATTGTTGGG |
| MKI67 | 47 | 5 | CTTAGGAGTTTGTGGCCATCTTTTGC | ATAGGAAATGGTGGTAGTGT | ATAGGAAATGGTGGTAGTGT | ATAGGAAATGGTGGTAGTGTTTTATAGGAAATGGTGGTAGTGTTTTCTTAGGAGTTTGTGGCCATCTTTTGC |
| MKI67 | 48 | 5 | GGAGTCTGTGGCTGTCTCTTGCTGCC | GGGATGTATTGAAGGAGGAT | GGGATGTATTGAAGGAGGAT | GGGATGTATTGAAGGAGGATTTTGGGATGTATTGAAGGAGGATTTTGGAGTCTGTGGCTGTCTCTTGCTGCC |
| MKI67 | 49 | 5 | TGTCTGGAAGAGTTCTTTGAAGCCAA | ATAGGAAATGGTGGTAGTGT | ATAGGAAATGGTGGTAGTGT | ATAGGAAATGGTGGTAGTGTTTTATAGGAAATGGTGGTAGTGTTTTTGTCTGGAAGAGTTCTTTGAAGCCAA |
| MKI67 | 50 | 5 | TTCTTTCATGTCCACTTTCACCAGGG | GGGATGTATTGAAGGAGGAT | GGGATGTATTGAAGGAGGAT | GGGATGTATTGAAGGAGGATTTTGGGATGTATTGAAGGAGGATTTTTTCTTTCATGTCCACTTTCACCAGGG |
| MKI67 | 51 | 5 | CTCTGTGTGTGTTTGCGTAGTCTCCC | ATAGGAAATGGTGGTAGTGT | ATAGGAAATGGTGGTAGTGT | ATAGGAAATGGTGGTAGTGTTTTATAGGAAATGGTGGTAGTGTTTTCTCTGTGTGTGTTTGCGTAGTCTCCC |
| MKI67 | 52 | 5 | GCTTTGATGCTCTTACTATCTCCTGT | GGGATGTATTGAAGGAGGAT | GGGATGTATTGAAGGAGGAT | GGGATGTATTGAAGGAGGATTTTGGGATGTATTGAAGGAGGATTTTGCTTTGATGCTCTTACTATCTCCTGT |
| MKI67 | 53 | 5 | TGAAGTCAACCAGGTCTTCTAGAGCA | ATAGGAAATGGTGGTAGTGT | ATAGGAAATGGTGGTAGTGT | ATAGGAAATGGTGGTAGTGTTTTATAGGAAATGGTGGTAGTGTTTTTGAAGTCAACCAGGTCTTCTAGAGCA |
| MKI67 | 54 | 5 | TGGCAGTGTCTGTTAGTTCTGGTGGG | GGGATGTATTGAAGGAGGAT | GGGATGTATTGAAGGAGGAT | GGGATGTATTGAAGGAGGATTTTGGGATGTATTGAAGGAGGATTTTTGGCAGTGTCTGTTAGTTCTGGTGGG |
| MKI67 | 55 | 5 | GTGTGTGCTTTGCCCTGATGTTTGCG | ATAGGAAATGGTGGTAGTGT | ATAGGAAATGGTGGTAGTGT | ATAGGAAATGGTGGTAGTGTTTTATAGGAAATGGTGGTAGTGTTTTGTGTGTGCTTTGCCCTGATGTTTGCG |
| MKI67 | 56 | 5 | TCATCACCGCTTGCTGGTTCTTTGTG | GGGATGTATTGAAGGAGGAT | GGGATGTATTGAAGGAGGAT | GGGATGTATTGAAGGAGGATTTTGGGATGTATTGAAGGAGGATTTTTCATCACCGCTTGCTGGTTCTTTGTG |
| MKI67 | 57 | 5 | TTCCTCTTCTACTGGGTTTGGTTTCT | ATAGGAAATGGTGGTAGTGT | ATAGGAAATGGTGGTAGTGT | ATAGGAAATGGTGGTAGTGTTTTATAGGAAATGGTGGTAGTGTTTTTTCCTCTTCTACTGGGTTTGGTTTCT |
| MKI67 | 58 | 5 | TTAGGTGCTCTTGGCCTTCTCCTGCT | GGGATGTATTGAAGGAGGAT | GGGATGTATTGAAGGAGGAT | GGGATGTATTGAAGGAGGATTTTGGGATGTATTGAAGGAGGATTTTTTAGGTGCTCTTGGCCTTCTCCTGCT |
| MKI67 | 59 | 5 | CGCAGGGTATTTTAGTGGCTTTGCCA | ATAGGAAATGGTGGTAGTGT | ATAGGAAATGGTGGTAGTGT | ATAGGAAATGGTGGTAGTGTTTTATAGGAAATGGTGGTAGTGTTTTCGCAGGGTATTTTAGTGGCTTTGCCA |
| MKI67 | 60 | 5 | ACGTGTCCTGAGATGCCTCTTTGTGC | GGGATGTATTGAAGGAGGAT | GGGATGTATTGAAGGAGGAT | GGGATGTATTGAAGGAGGATTTTGGGATGTATTGAAGGAGGATTTTACGTGTCCTGAGATGCCTCTTTGTGC |
| MKI67 | 61 | 5 | CCTGATGTTTGTGTGAACTTGACTGC | ATAGGAAATGGTGGTAGTGT | ATAGGAAATGGTGGTAGTGT | ATAGGAAATGGTGGTAGTGTTTTATAGGAAATGGTGGTAGTGTTTTCCTGATGTTTGTGTGAACTTGACTGC |
| MKI67 | 62 | 5 | CTGGTTCTTTGTCTGCATCCGTGGTT | GGGATGTATTGAAGGAGGAT | GGGATGTATTGAAGGAGGAT | GGGATGTATTGAAGGAGGATTTTGGGATGTATTGAAGGAGGATTTTCTGGTTCTTTGTCTGCATCCGTGGTT |
| MKI67 | 63 | 5 | TGGAGCCGGTGTCTGTTTTGCAGATT | ATAGGAAATGGTGGTAGTGT | ATAGGAAATGGTGGTAGTGT | ATAGGAAATGGTGGTAGTGTTTTATAGGAAATGGTGGTAGTGTTTTTGGAGCCGGTGTCTGTTTTGCAGATT |
| MKI67 | 64 | 5 | CGTCTCCTGCTGCCAGTTACACTTGC | GGGATGTATTGAAGGAGGAT | GGGATGTATTGAAGGAGGAT | GGGATGTATTGAAGGAGGATTTTGGGATGTATTGAAGGAGGATTTTCGTCTCCTGCTGCCAGTTACACTTGC |
| MKI67 | 65 | 5 | CGGTGTCTTCTAGTTCTGGTGATGAT | ATAGGAAATGGTGGTAGTGT | ATAGGAAATGGTGGTAGTGT | ATAGGAAATGGTGGTAGTGTTTTATAGGAAATGGTGGTAGTGTTTTCGGTGTCTTCTAGTTCTGGTGATGAT |
| MKI67 | 66 | 5 | TGTCCTGGGCCGTCTCTTTGAGCTTG | GGGATGTATTGAAGGAGGAT | GGGATGTATTGAAGGAGGAT | GGGATGTATTGAAGGAGGATTTTGGGATGTATTGAAGGAGGATTTTTGTCCTGGGCCGTCTCTTTGAGCTTG |
| MKI67 | 67 | 5 | GCTTTCGTGCCTTTGCCCTCACCTAC | ATAGGAAATGGTGGTAGTGT | ATAGGAAATGGTGGTAGTGT | ATAGGAAATGGTGGTAGTGTTTTATAGGAAATGGTGGTAGTGTTTTGCTTTCGTGCCTTTGCCCTCACCTAC |
| MKI67 | 68 | 5 | TCTCCTGCTGCCAATTACATCTTCTG | GGGATGTATTGAAGGAGGAT | GGGATGTATTGAAGGAGGAT | GGGATGTATTGAAGGAGGATTTTGGGATGTATTGAAGGAGGATTTTTCTCCTGCTGCCAATTACATCTTCTG |
| MKI67 | 69 | 5 | TTGTGATTTTACAGGGTCTCTGGTGC | ATAGGAAATGGTGGTAGTGT | ATAGGAAATGGTGGTAGTGT | ATAGGAAATGGTGGTAGTGTTTTATAGGAAATGGTGGTAGTGTTTTTTGTGATTTTACAGGGTCTCTGGTGC |
| MKI67 | 70 | 5 | TGACGCTTCCATCTTTGCCACCTCCC | GGGATGTATTGAAGGAGGAT | GGGATGTATTGAAGGAGGAT | GGGATGTATTGAAGGAGGATTTTGGGATGTATTGAAGGAGGATTTTTGACGCTTCCATCTTTGCCACCTCCC |
| MKI67 | 71 | 5 | ACGGGTTCGGATGATTTGCCTCTTGC | ATAGGAAATGGTGGTAGTGT | ATAGGAAATGGTGGTAGTGT | ATAGGAAATGGTGGTAGTGTTTTATAGGAAATGGTGGTAGTGTTTTACGGGTTCGGATGATTTGCCTCTTGC |
| MKI67 | 72 | 5 | CTTTGTTGGTTTTCATGTCGTTGCTG | GGGATGTATTGAAGGAGGAT | GGGATGTATTGAAGGAGGAT | GGGATGTATTGAAGGAGGATTTTGGGATGTATTGAAGGAGGATTTTCTTTGTTGGTTTTCATGTCGTTGCTG |
| MKI67 | 73 | 5 | TCTTTCTGCTAATACAAAGACCTCAGTTA | ATAGGAAATGGTGGTAGTGT | ATAGGAAATGGTGGTAGTGT | ATAGGAAATGGTGGTAGTGTTTTATAGGAAATGGTGGTAGTGTTTTTCTTTCTGCTAATACAAAGACCTCAGTTA |
| MKI67 | 74 | 5 | CTGGATTCTGAATGTCCATCTCTGGG | GGGATGTATTGAAGGAGGAT | GGGATGTATTGAAGGAGGAT | GGGATGTATTGAAGGAGGATTTTGGGATGTATTGAAGGAGGATTTTCTGGATTCTGAATGTCCATCTCTGGG |
| MKI67 | 75 | 5 | AAGCACCTTTTGTTCTCAGTGACTTT | ATAGGAAATGGTGGTAGTGT | ATAGGAAATGGTGGTAGTGT | ATAGGAAATGGTGGTAGTGTTTTATAGGAAATGGTGGTAGTGTTTTAAGCACCTTTTGTTCTCAGTGACTTT |
| MKI67 | 76 | 5 | GGGAGCTCTCATTCTGTCTAGCAGAC | GGGATGTATTGAAGGAGGAT | GGGATGTATTGAAGGAGGAT | GGGATGTATTGAAGGAGGATTTTGGGATGTATTGAAGGAGGATTTTGGGAGCTCTCATTCTGTCTAGCAGAC |
| MKI67 | 77 | 5 | TTCTGCATGAGAACCTTCGCACTCTT | ATAGGAAATGGTGGTAGTGT | ATAGGAAATGGTGGTAGTGT | ATAGGAAATGGTGGTAGTGTTTTATAGGAAATGGTGGTAGTGTTTTTTCTGCATGAGAACCTTCGCACTCTT |
| MKI67 | 78 | 5 | CTTTCTTGATCTCAGGCACATGGAGT | GGGATGTATTGAAGGAGGAT | GGGATGTATTGAAGGAGGAT | GGGATGTATTGAAGGAGGATTTTGGGATGTATTGAAGGAGGATTTTCTTTCTTGATCTCAGGCACATGGAGT |
| MKI67 | 79 | 5 | TGGATTTTCTGCACACCTCTTGACAC | ATAGGAAATGGTGGTAGTGT | ATAGGAAATGGTGGTAGTGT | ATAGGAAATGGTGGTAGTGTTTTATAGGAAATGGTGGTAGTGTTTTTGGATTTTCTGCACACCTCTTGACAC |
| MKI67 | 80 | 5 | TTGACACACACATTGTCCTCAGCCT | GGGATGTATTGAAGGAGGAT | GGGATGTATTGAAGGAGGAT | GGGATGTATTGAAGGAGGATTTTGGGATGTATTGAAGGAGGATTTTTTGACACACACATTGTCCTCAGCCT |
| BRCA1 | 1 | 6 | TGTAGAAGTCTTTTGGCACGGTTTC | TGTGATGGAAGTTAGAGGGT | TGTGATGGAAGTTAGAGGGT | TGTGATGGAAGTTAGAGGGTTTTTGTGATGGAAGTTAGAGGGTTTTTGTAGAAGTCTTTTGGCACGGTTTC |
| BRCA1 | 2 | 6 | GTTAGAGAGTTGGACACTGAGACTG | GGGATGTATTGAAGGAGGAT | GGGATGTATTGAAGGAGGAT | GGGATGTATTGAAGGAGGATTTTGGGATGTATTGAAGGAGGATTTTGTTAGAGAGTTGGACACTGAGACTG |
| BRCA1 | 3 | 6 | ATACTTTTCTGGATGCCTCTCAGCT | TGTGATGGAAGTTAGAGGGT | TGTGATGGAAGTTAGAGGGT | TGTGATGGAAGTTAGAGGGTTTTTGTGATGGAAGTTAGAGGGTTTTATACTTTTCTGGATGCCTCTCAGCT |

| | | | | | | |
|---|---|---|---|---|---|---|
| BRCA1 | 4 | 6 | CCACATGCAAGTTTGAAACAGAACTAC | GGGATGTATTGAAGGAGGAT | GGGATGTATTGAAGGAGGAT | GGGATGTATTGAAGGAGGATTTTGGGATGTATTGAAGGAGGATTTTCCACATGCAAGTTTGAAACAGAACTAC |
| BRCA1 | 5 | 6 | AGTAATAAACTGCTGTTCTCATGCTGTA | TGTGATGGAAGTTAGAGGGT | TGTGATGGAAGTTAGAGGGT | TGTGATGGAAGTTAGAGGGTTTTTGTGATGGAAGTTAGAGGGTTTTAGTAATAAACTGCTGTTCTCATGCTGTA |
| BRCA1 | 6 | 6 | CAGCCTTTTCTACATTCATTCTGTCTTT | GGGATGTATTGAAGGAGGAT | GGGATGTATTGAAGGAGGAT | GGGATGTATTGAAGGAGGATTTTGGGATGTATTGAAGGAGGATTTTCAGCCTTTTCTACATTCATTCTGTCTTT |
| BRCA1 | 7 | 6 | GCCCATCTGTTATGTTGGCTCCTTG | TGTGATGGAAGTTAGAGGGT | TGTGATGGAAGTTAGAGGGT | TGTGATGGAAGTTAGAGGGTTTTTGTGATGGAAGTTAGAGGGTTTTGCCCATCTGTTATGTTGGCTCCTTG |
| BRCA1 | 8 | 6 | CCGCCTATCATTACATGTTTCCTTACT | GGGATGTATTGAAGGAGGAT | GGGATGTATTGAAGGAGGAT | GGGATGTATTGAAGGAGGATTTTGGGATGTATTGAAGGAGGATTTTCCGCCTATCATTACATGTTTCCTTACT |
| BRCA1 | 9 | 6 | CTGCTATTTAGTGTTATCCAAGGAACATC | TGTGATGGAAGTTAGAGGGT | TGTGATGGAAGTTAGAGGGT | TGTGATGGAAGTTAGAGGGTTTTTGTGATGGAAGTTAGAGGGTTTTCTGCTATTTAGTGTTATCCAAGGAACATC |
| BRCA1 | 10 | 6 | TCTGGAAAACCACTCATTAACTTTCTG | GGGATGTATTGAAGGAGGAT | GGGATGTATTGAAGGAGGAT | GGGATGTATTGAAGGAGGATTTTGGGATGTATTGAAGGAGGATTTTTCTGGAAAACCACTCATTAACTTTCTG |
| BRCA1 | 11 | 6 | ACTTTGGCATTTGATTCAGACTCCC | TGTGATGGAAGTTAGAGGGT | TGTGATGGAAGTTAGAGGGT | TGTGATGGAAGTTAGAGGGTTTTTGTGATGGAAGTTAGAGGGTTTTACTTTGGCATTTGATTCAGACTCCC |
| BRCA1 | 12 | 6 | TCTACCTCATTTAGAACGTCCAATACATC | GGGATGTATTGAAGGAGGAT | GGGATGTATTGAAGGAGGAT | GGGATGTATTGAAGGAGGATTTTGGGATGTATTGAAGGAGGATTTTTCTACCTCATTTAGAACGTCCAATACATC |
| BRCA1 | 13 | 6 | TTTTCAGTTACATGGCTTAAGTTGGG | TGTGATGGAAGTTAGAGGGT | TGTGATGGAAGTTAGAGGGT | TGTGATGGAAGTTAGAGGGTTTTTGTGATGGAAGTTAGAGGGTTTTTTTTCAGTTACATGGCTTAAGTTGGG |
| BRCA1 | 14 | 6 | TGGCTCAGTAACAAATGCTCCTATAAT | GGGATGTATTGAAGGAGGAT | GGGATGTATTGAAGGAGGAT | GGGATGTATTGAAGGAGGATTTTGGGATGTATTGAAGGAGGATTTTTGGCTCAGTAACAAATGCTCCTATAAT |
| BRCA1 | 15 | 6 | TCCTCAGGATGAAGGCCTGATGTAG | TGTGATGGAAGTTAGAGGGT | TGTGATGGAAGTTAGAGGGT | TGTGATGGAAGTTAGAGGGTTTTTGTGATGGAAGTTAGAGGGTTTTTCCTCAGGATGAAGGCCTGATGTAG |
| BRCA1 | 16 | 6 | GAACTGCCAAATCTGCTTTCTTGAT | GGGATGTATTGAAGGAGGAT | GGGATGTATTGAAGGAGGAT | GGGATGTATTGAAGGAGGATTTTGGGATGTATTGAAGGAGGATTTTGAACTGCCAAATCTGCTTTCTTGAT |
| BRCA1 | 17 | 6 | ATCACTTGACCATTCTGCTCCGTTT | TGTGATGGAAGTTAGAGGGT | TGTGATGGAAGTTAGAGGGT | TGTGATGGAAGTTAGAGGGTTTTTGTGATGGAAGTTAGAGGGTTTTATCACTTGACCATTCTGCTCCGTTT |
| BRCA1 | 18 | 6 | TCGAGTGATTCTATTGGGTTAGGATTTT | GGGATGTATTGAAGGAGGAT | GGGATGTATTGAAGGAGGAT | GGGATGTATTGAAGGAGGATTTTGGGATGTATTGAAGGAGGATTTTTCGAGTGATTCTATTGGGTTAGGATTTT |
| BRCA1 | 19 | 6 | GTTCAAGCGCATGAATATGCCTGGT | TGTGATGGAAGTTAGAGGGT | TGTGATGGAAGTTAGAGGGT | TGTGATGGAAGTTAGAGGGTTTTTGTGATGGAAGTTAGAGGGTTTTGTTCAAGCGCATGAATATGCCTGGT |
| BRCA1 | 20 | 6 | AGGTGGGCTTAGATTTCTACTGACT | GGGATGTATTGAAGGAGGAT | GGGATGTATTGAAGGAGGAT | GGGATGTATTGAAGGAGGATTTTGGGATGTATTGAAGGAGGATTTTAGGTGGGCTTAGATTTCTACTGACT |
| BRCA1 | 21 | 6 | CTGACTGGCATTTGGTTGTACTTTT | TGTGATGGAAGTTAGAGGGT | TGTGATGGAAGTTAGAGGGT | TGTGATGGAAGTTAGAGGGTTTTTGTGATGGAAGTTAGAGGGTTTTCTGACTGGCATTTGGTTGTACTTTT |
| BRCA1 | 22 | 6 | CTTCCATGAGTTGTAGGTTTCTGCT | GGGATGTATTGAAGGAGGAT | GGGATGTATTGAAGGAGGAT | GGGATGTATTGAAGGAGGATTTTGGGATGTATTGAAGGAGGATTTTCTTCCATGAGTTGTAGGTTTCTGCT |
| BRCA1 | 23 | 6 | TTTACTTGTCTGTTCATTTGGCTTGT | TGTGATGGAAGTTAGAGGGT | TGTGATGGAAGTTAGAGGGT | TGTGATGGAAGTTAGAGGGTTTTTGTGATGGAAGTTAGAGGGTTTTTTTACTTGTCTGTTCATTTGGCTTGT |
| BRCA1 | 24 | 6 | AGCTCTGGGAAAGTATCGCTGTCAT | GGGATGTATTGAAGGAGGAT | GGGATGTATTGAAGGAGGAT | GGGATGTATTGAAGGAGGATTTTGGGATGTATTGAAGGAGGATTTTAGCTCTGGGAAAGTATCGCTGTCAT |
| BRCA1 | 25 | 6 | TGGAAGGCTAGGATTGACAAATTCTTT | TGTGATGGAAGTTAGAGGGT | TGTGATGGAAGTTAGAGGGT | TGTGATGGAAGTTAGAGGGTTTTTGTGATGGAAGTTAGAGGGTTTTTGGAAGGCTAGGATTGACAAATTCTTT |
| BRCA1 | 26 | 6 | GGTCTTCAGCATTATTAGACACTTTAACTG | GGGATGTATTGAAGGAGGAT | GGGATGTATTGAAGGAGGAT | GGGATGTATTGAAGGAGGATTTTGGGATGTATTGAAGGAGGATTTTGGTCTTCAGCATTATTAGACACTTTAACTG |
| BRCA1 | 27 | 6 | GCTACTCTCTACAGATCTTTCAGTTTGC | TGTGATGGAAGTTAGAGGGT | TGTGATGGAAGTTAGAGGGT | TGTGATGGAAGTTAGAGGGTTTTTGTGATGGAAGTTAGAGGGTTTTGCTACTCTCTACAGATCTTTCAGTTTGC |
| BRCA1 | 28 | 6 | CCATAATCAGTACCAGGTACCAATGAAA | GGGATGTATTGAAGGAGGAT | GGGATGTATTGAAGGAGGAT | GGGATGTATTGAAGGAGGATTTTGGGATGTATTGAAGGAGGATTTTCCATAATCAGTACCAGGTACCAATGAAA |
| BRCA1 | 29 | 6 | AACAACCATGAATTAGTCCCTTGGG | TGTGATGGAAGTTAGAGGGT | TGTGATGGAAGTTAGAGGGT | TGTGATGGAAGTTAGAGGGTTTTTGTGATGGAAGTTAGAGGGTTTTAACAACCATGAATTAGTCCCTTGGG |
| BRCA1 | 30 | 6 | TGGATACTTAAAGCCTTCTGTGTCATT | GGGATGTATTGAAGGAGGAT | GGGATGTATTGAAGGAGGAT | GGGATGTATTGAAGGAGGATTTTGGGATGTATTGAAGGAGGATTTTTGGATACTTAAAGCCTTCTGTGTCATT |
| BRCA1 | 31 | 6 | GCGCTTTGAAACCTTGAATGTATTCTG | TGTGATGGAAGTTAGAGGGT | TGTGATGGAAGTTAGAGGGT | TGTGATGGAAGTTAGAGGGTTTTTGTGATGGAAGTTAGAGGGTTTTGCGCTTTGAAACCTTGAATGTATTCTG |
| BRCA1 | 32 | 6 | CCTGGATTTGAAAACGGAGCAAATG | GGGATGTATTGAAGGAGGAT | GGGATGTATTGAAGGAGGAT | GGGATGTATTGAAGGAGGATTTTGGGATGTATTGAAGGAGGATTTTCCTGGATTTGAAAACGGAGCAAATG |
| BRCA1 | 33 | 6 | TTGTTTCTTTAAGGACCCAGAGTGG | TGTGATGGAAGTTAGAGGGT | TGTGATGGAAGTTAGAGGGT | TGTGATGGAAGTTAGAGGGTTTTTGTGATGGAAGTTAGAGGGTTTTTTGTTTCTTTAAGGACCCAGAGTGG |
| BRCA1 | 34 | 6 | AGCCTGCAGTGATATTAACTGTCTG | GGGATGTATTGAAGGAGGAT | GGGATGTATTGAAGGAGGAT | GGGATGTATTGAAGGAGGATTTTGGGATGTATTGAAGGAGGATTTTAGCCTGCAGTGATATTAACTGTCTG |
| BRCA1 | 35 | 6 | GCCTCCTTTGATACTACATTTGGCATTAT | TGTGATGGAAGTTAGAGGGT | TGTGATGGAAGTTAGAGGGT | TGTGATGGAAGTTAGAGGGTTTTTGTGATGGAAGTTAGAGGGTTTTGCCTCCTTTGATACTACATTTGGCATTAT |
| BRCA1 | 36 | 6 | CCTCTGAACTGAGATGATAGACAAAACC | GGGATGTATTGAAGGAGGAT | GGGATGTATTGAAGGAGGAT | GGGATGTATTGAAGGAGGATTTTGGGATGTATTGAAGGAGGATTTTCCTCTGAACTGAGATGATAGACAAAACC |
| BRCA1 | 37 | 6 | GGTGGTATACGATATGGGTTTTGTAAAAG | TGTGATGGAAGTTAGAGGGT | TGTGATGGAAGTTAGAGGGT | TGTGATGGAAGTTAGAGGGTTTTTGTGATGGAAGTTAGAGGGTTTTGGTGGTATACGATATGGGTTTTGTAAAAG |
| BRCA1 | 38 | 6 | CCATTTCTCTTTCAGGTGACATTGAA | GGGATGTATTGAAGGAGGAT | GGGATGTATTGAAGGAGGAT | GGGATGTATTGAAGGAGGATTTTGGGATGTATTGAAGGAGGATTTTCCATTTCTCTTTCAGGTGACATTGAA |
| BRCA1 | 39 | 6 | CACTTCATTAGTACTGGAACCTACTTCAT | TGTGATGGAAGTTAGAGGGT | TGTGATGGAAGTTAGAGGGT | TGTGATGGAAGTTAGAGGGTTTTTGTGATGGAAGTTAGAGGGTTTTCACTTCATTAGTACTGGAACCTACTTCAT |
| BRCA1 | 40 | 6 | CTGTTTCTACCTAGTTCTGCTTGAATG | GGGATGTATTGAAGGAGGAT | GGGATGTATTGAAGGAGGAT | GGGATGTATTGAAGGAGGATTTTGGGATGTATTGAAGGAGGATTTTCTGTTTCTACCTAGTTCTGCTTGAATG |
| BRCA1 | 41 | 6 | ACTTTGTTTATAGACCTCAGGTTGCA | TGTGATGGAAGTTAGAGGGT | TGTGATGGAAGTTAGAGGGT | TGTGATGGAAGTTAGAGGGTTTTTGTGATGGAAGTTAGAGGGTTTTACTTTGTTTATAGACCTCAGGTTGCA |
| BRCA1 | 42 | 6 | ATTTCAGGATGCTTACAATTACTTCCAG | GGGATGTATTGAAGGAGGAT | GGGATGTATTGAAGGAGGAT | GGGATGTATTGAAGGAGGATTTTGGGATGTATTGAAGGAGGATTTTATTTCAGGATGCTTACAATTACTTCCAG |
| BRCA1 | 43 | 6 | GAACAAACCTGAGATGCATGACTAC | TGTGATGGAAGTTAGAGGGT | TGTGATGGAAGTTAGAGGGT | TGTGATGGAAGTTAGAGGGTTTTTGTGATGGAAGTTAGAGGGTTTTGAACAAACCTGAGATGCATGACTAC |
| BRCA1 | 44 | 6 | CATCATCTAACAGGTCATCAGGTGT | GGGATGTATTGAAGGAGGAT | GGGATGTATTGAAGGAGGAT | GGGATGTATTGAAGGAGGATTTTGGGATGTATTGAAGGAGGATTTTCATCATCTAACAGGTCATCAGGTGT |
| BRCA1 | 45 | 6 | TCCTTTCTGGACGCTTTTGCTAAAA | TGTGATGGAAGTTAGAGGGT | TGTGATGGAAGTTAGAGGGT | TGTGATGGAAGTTAGAGGGTTTTTGTGATGGAAGTTAGAGGGTTTTTCCTTTCTGGACGCTTTTGCTAAAA |
| BRCA1 | 46 | 6 | GTGAAAGGGCTAGGACTCCTGCTAA | GGGATGTATTGAAGGAGGAT | GGGATGTATTGAAGGAGGAT | GGGATGTATTGAAGGAGGATTTTGGGATGTATTGAAGGAGGATTTTGTGAAAGGGCTAGGACTCCTGCTAA |
| BRCA1 | 47 | 6 | TTCTGAGGACTCTAATTTCTTGGCC | TGTGATGGAAGTTAGAGGGT | TGTGATGGAAGTTAGAGGGT | TGTGATGGAAGTTAGAGGGTTTTTGTGATGGAAGTTAGAGGGTTTTTTCTGAGGACTCTAATTTCTTGGCC |
| BRCA1 | 48 | 6 | GAAGCTCTTCATCCTCACTAGATAAGT | GGGATGTATTGAAGGAGGAT | GGGATGTATTGAAGGAGGAT | GGGATGTATTGAAGGAGGATTTTGGGATGTATTGAAGGAGGATTTTGAAGCTCTTCATCCTCACTAGATAAGT |
| BRCA1 | 49 | 6 | TCTTAGACAGACACTCGGTAGCAAC | TGTGATGGAAGTTAGAGGGT | TGTGATGGAAGTTAGAGGGT | TGTGATGGAAGTTAGAGGGTTTTTGTGATGGAAGTTAGAGGGTTTTTCTTAGACAGACACTCGGTAGCAAC |
| BRCA1 | 50 | 6 | GTTACTGCAGTCATTTAAGCTATTCTTCA | GGGATGTATTGAAGGAGGAT | GGGATGTATTGAAGGAGGAT | GGGATGTATTGAAGGAGGATTTTGGGATGTATTGAAGGAGGATTTTGTTACTGCAGTCATTTAAGCTATTCTTCA |
| BRCA1 | 51 | 6 | AGAACATTTTGTTTCCTCACTAAGGTG | TGTGATGGAAGTTAGAGGGT | TGTGATGGAAGTTAGAGGGT | TGTGATGGAAGTTAGAGGGTTTTTGTGATGGAAGTTAGAGGGTTTTAGAACATTTTGTTTCCTCACTAAGGTG |
| BRCA1 | 52 | 6 | TCACTGCACTGTGAAGAAAACAAGC | GGGATGTATTGAAGGAGGAT | GGGATGTATTGAAGGAGGAT | GGGATGTATTGAAGGAGGATTTTGGGATGTATTGAAGGAGGATTTTTCACTGCACTGTGAAGAAAACAAGC |
| BRCA1 | 53 | 6 | GTTTGGAAGAACCAATCAAGAAAGGAT | TGTGATGGAAGTTAGAGGGT | TGTGATGGAAGTTAGAGGGT | TGTGATGGAAGTTAGAGGGTTTTTGTGATGGAAGTTAGAGGGTTTTGTTTGGAAGAACCAATCAAGAAAGGAT |
| BRCA1 | 54 | 6 | TCCCTGGCTTTCAGACTGATGCCTC | GGGATGTATTGAAGGAGGAT | GGGATGTATTGAAGGAGGAT | GGGATGTATTGAAGGAGGATTTTGGGATGTATTGAAGGAGGATTTTTCCCTGGCTTTCAGACTGATGCCTC |
| BRCA1 | 55 | 6 | CATTTCCTGCTGGAGCTTTATCAGG | TGTGATGGAAGTTAGAGGGT | TGTGATGGAAGTTAGAGGGT | TGTGATGGAAGTTAGAGGGTTTTTGTGATGGAAGTTAGAGGGTTTTCATTTCCTGCTGGAGCTTTATCAGG |
| BRCA1 | 56 | 6 | TGCTGTTCTAACACAGCTTCTAGTT | GGGATGTATTGAAGGAGGAT | GGGATGTATTGAAGGAGGAT | GGGATGTATTGAAGGAGGATTTTGGGATGTATTGAAGGAGGATTTTTGCTGTTCTAACACAGCTTCTAGTT |
| BRCA1 | 57 | 6 | CTCAAGGGCAGAAGAGTCACTTATG | TGTGATGGAAGTTAGAGGGT | TGTGATGGAAGTTAGAGGGT | TGTGATGGAAGTTAGAGGGTTTTTGTGATGGAAGTTAGAGGGTTTTCTCAAGGGCAGAAGAGTCACTTATG |
| BRCA1 | 58 | 6 | AGGGTATTCACTACTTTTCTGTGAAGTTA | GGGATGTATTGAAGGAGGAT | GGGATGTATTGAAGGAGGAT | GGGATGTATTGAAGGAGGATTTTGGGATGTATTGAAGGAGGATTTTAGGGTATTCACTACTTTTCTGTGAAGTTA |
| BRCA1 | 59 | 6 | AACTATCTGCAGACACCTCAAACTT | TGTGATGGAAGTTAGAGGGT | TGTGATGGAAGTTAGAGGGT | TGTGATGGAAGTTAGAGGGTTTTTGTGATGGAAGTTAGAGGGTTTTAACTATCTGCAGACACCTCAAACTT |
| BRCA1 | 60 | 6 | ACCTTTCCACTCCTGGTTCTTTATTTT | GGGATGTATTGAAGGAGGAT | GGGATGTATTGAAGGAGGAT | GGGATGTATTGAAGGAGGATTTTGGGATGTATTGAAGGAGGATTTTACCTTTCCACTCCTGGTTCTTTATTTT |
| BRCA1 | 61 | 6 | CTCTTGAGATGGGTAGTTTCTATTCTGA | TGTGATGGAAGTTAGAGGGT | TGTGATGGAAGTTAGAGGGT | TGTGATGGAAGTTAGAGGGTTTTTGTGATGGAAGTTAGAGGGTTTTCTCTTGAGATGGGTAGTTTCTATTCTGA |
| BRCA1 | 62 | 6 | TCCTCCACATCAACAACCTTAATGA | GGGATGTATTGAAGGAGGAT | GGGATGTATTGAAGGAGGAT | GGGATGTATTGAAGGAGGATTTTGGGATGTATTGAAGGAGGATTTTTCCTCCACATCAACAACCTTAATGA |
| BRCA1 | 63 | 6 | CCTTGGCAAGTAAGATGTTTCCGTC | TGTGATGGAAGTTAGAGGGT | TGTGATGGAAGTTAGAGGGT | TGTGATGGAAGTTAGAGGGTTTTTGTGATGGAAGTTAGAGGGTTTTCCTTGGCAAGTAAGATGTTTCCGTC |
| BRCA1 | 64 | 6 | GCTGATTCCAGATTCCAGGTAAGGG | GGGATGTATTGAAGGAGGAT | GGGATGTATTGAAGGAGGAT | GGGATGTATTGAAGGAGGATTTTGGGATGTATTGAAGGAGGATTTTGCTGATTCCAGATTCCAGGTAAGGG |
| BRCA1 | 65 | 6 | GTTGCCAACACGAGCTGACTCTGGG | TGTGATGGAAGTTAGAGGGT | TGTGATGGAAGTTAGAGGGT | TGTGATGGAAGTTAGAGGGTTTTTGTGATGGAAGTTAGAGGGTTTTGTTGCCAACACGAGCTGACTCTGGG |
| BRCA1 | 66 | 6 | ACTTTCAATGCAGAGGTTGAAGATG | GGGATGTATTGAAGGAGGAT | GGGATGTATTGAAGGAGGAT | GGGATGTATTGAAGGAGGATTTTGGGATGTATTGAAGGAGGATTTTACTTTCAATGCAGAGGTTGAAGATG |
| BRCA1 | 67 | 6 | ATCAGTAGTATGAGCAGCAGCTGGA | TGTGATGGAAGTTAGAGGGT | TGTGATGGAAGTTAGAGGGT | TGTGATGGAAGTTAGAGGGTTTTTGTGATGGAAGTTAGAGGGTTTTATCAGTAGTATGAGCAGCAGCTGGA |
| BRCA1 | 68 | 6 | CACTTTCTTCCATTGCATTATACCCAG | GGGATGTATTGAAGGAGGAT | GGGATGTATTGAAGGAGGAT | GGGATGTATTGAAGGAGGATTTTGGGATGTATTGAAGGAGGATTTTCACTTTCTTCCATTGCATTATACCCAG |
| BRCA1 | 69 | 6 | CACCCACTCTCGGGTCACCACAGGT | TGTGATGGAAGTTAGAGGGT | TGTGATGGAAGTTAGAGGGT | TGTGATGGAAGTTAGAGGGTTTTTGTGATGGAAGTTAGAGGGTTTTCACCCACTCTCGGGTCACCACAGGT |
| BRCA1 | 70 | 6 | GGCTGCAGTCAGTAGTGGCTGTGGG | GGGATGTATTGAAGGAGGAT | GGGATGTATTGAAGGAGGAT | GGGATGTATTGAAGGAGGATTTTGGGATGTATTGAAGGAGGATTTTGGCTGCAGTCAGTAGTGGCTGTGGG |
| BRCA1 | 71 | 6 | CAGATTTCCAAGGGAGACTTCAAGC | TGTGATGGAAGTTAGAGGGT | TGTGATGGAAGTTAGAGGGT | TGTGATGGAAGTTAGAGGGTTTTTGTGATGGAAGTTAGAGGGTTTTCAGATTTCCAAGGGAGACTTCAAGC |
| BRCA1 | 72 | 6 | TCAATTGGTGGCGTTTAAATGGTTTT | GGGATGTATTGAAGGAGGAT | GGGATGTATTGAAGGAGGAT | GGGATGTATTGAAGGAGGATTTTGGGATGTATTGAAGGAGGATTTTTCAATTGGTGGCGTTTAAATGGTTTT |
| BRCA1 | 73 | 6 | TGAGGCCAAGCCACTCTGTGCTTCC | TGTGATGGAAGTTAGAGGGT | TGTGATGGAAGTTAGAGGGT | TGTGATGGAAGTTAGAGGGTTTTTGTGATGGAAGTTAGAGGGTTTTTGAGGCCAAGCCACTCTGTGCTTCC |
| BRCA1 | 74 | 6 | AGAAGTAAACTTAGGGAAACCAGCTATT | GGGATGTATTGAAGGAGGAT | GGGATGTATTGAAGGAGGAT | GGGATGTATTGAAGGAGGATTTTGGGATGTATTGAAGGAGGATTTTAGAAGTAAACTTAGGGAAACCAGCTATT |
| BRCA1 | 75 | 6 | CCCAAGGACTATTCTGACTTTAAGTCA | TGTGATGGAAGTTAGAGGGT | TGTGATGGAAGTTAGAGGGT | TGTGATGGAAGTTAGAGGGTTTTTGTGATGGAAGTTAGAGGGTTTTCCCAAGGACTATTCTGACTTTAAGTCA |
| BRCA1 | 76 | 6 | CCAATGTTCCACTCCAACATTTGAGA | GGGATGTATTGAAGGAGGAT | GGGATGTATTGAAGGAGGAT | GGGATGTATTGAAGGAGGATTTTGGGATGTATTGAAGGAGGATTTTCCAATGTTCCACTCCAACATTTGAGA |
| BRCA1 | 77 | 6 | GGAATGGATTATATACCAGAGCTACAACA | TGTGATGGAAGTTAGAGGGT | TGTGATGGAAGTTAGAGGGT | TGTGATGGAAGTTAGAGGGTTTTTGTGATGGAAGTTAGAGGGTTTTGGAATGGATTATATACCAGAGCTACAACA |
| BRCA1 | 78 | 6 | ATCACCTCAAAGAAAGCAACAGCTT | GGGATGTATTGAAGGAGGAT | GGGATGTATTGAAGGAGGAT | GGGATGTATTGAAGGAGGATTTTGGGATGTATTGAAGGAGGATTTTATCACCTCAAAGAAAGCAACAGCTT |
| BRCA1 | 79 | 6 | ACAGTCCTGGATAATGGGTTTATGAAAA | TGTGATGGAAGTTAGAGGGT | TGTGATGGAAGTTAGAGGGT | TGTGATGGAAGTTAGAGGGTTTTTGTGATGGAAGTTAGAGGGTTTTACAGTCCTGGATAATGGGTTTATGAAAA |
| BRCA1 | 80 | 6 | GTCTTCACTGCCCTTGCACACTGGG | GGGATGTATTGAAGGAGGAT | GGGATGTATTGAAGGAGGAT | GGGATGTATTGAAGGAGGATTTTGGGATGTATTGAAGGAGGATTTTGTCTTCACTGCCCTTGCACACTGGG |
| NCOA2 | 1 | 7 | TCACATTCTCTGACACAAACACAAC | TGTGATGGAAGTTAGAGGGT | TGTGATGGAAGTTAGAGGGT | TGTGATGGAAGTTAGAGGGTTTTTGTGATGGAAGTTAGAGGGTTTTTCACATTCTCTGACACAAACACAAC |
| NCOA2 | 2 | 7 | TCAGCTCTTCTTGGTTATACCTTAGATAC | ATAGGAAATGGTGGTAGTGT | ATAGGAAATGGTGGTAGTGT | ATAGGAAATGGTGGTAGTGTTTTATAGGAAATGGTGGTAGTGTTTTTCAGCTCTTCTTGGTTATACCTTAGATAC |
| NCOA2 | 3 | 7 | CGACAATTGAAGGTATGGCTGTTCC | TGTGATGGAAGTTAGAGGGT | TGTGATGGAAGTTAGAGGGT | TGTGATGGAAGTTAGAGGGTTTTTGTGATGGAAGTTAGAGGGTTTTCGACAATTGAAGGTATGGCTGTTCC |
| NCOA2 | 4 | 7 | GCGAAGCACTGCATAGTTTCATATT | ATAGGAAATGGTGGTAGTGT | ATAGGAAATGGTGGTAGTGT | ATAGGAAATGGTGGTAGTGTTTTATAGGAAATGGTGGTAGTGTTTTGCGAAGCACTGCATAGTTTCATATT |
| NCOA2 | 5 | 7 | CAAGAGTGCCATCAGACAAGGAAAA | TGTGATGGAAGTTAGAGGGT | TGTGATGGAAGTTAGAGGGT | TGTGATGGAAGTTAGAGGGTTTTTGTGATGGAAGTTAGAGGGTTTTCAAGAGTGCCATCAGACAAGGAAAA |
| NCOA2 | 6 | 7 | GATGAGTTTGCTCTTCGTTTGTGCA | ATAGGAAATGGTGGTAGTGT | ATAGGAAATGGTGGTAGTGT | ATAGGAAATGGTGGTAGTGTTTTATAGGAAATGGTGGTAGTGTTTTGATGAGTTTGCTCTTCGTTTGTGCA |
| NCOA2 | 7 | 7 | ATTGCTACTGAGGGTCATGTCCTGA | TGTGATGGAAGTTAGAGGGT | TGTGATGGAAGTTAGAGGGT | TGTGATGGAAGTTAGAGGGTTTTTGTGATGGAAGTTAGAGGGTTTTATTGCTACTGAGGGTCATGTCCTGA |
| NCOA2 | 8 | 7 | TCCTTTGGGCCATTTATGGGAAAAT | ATAGGAAATGGTGGTAGTGT | ATAGGAAATGGTGGTAGTGT | ATAGGAAATGGTGGTAGTGTTTTATAGGAAATGGTGGTAGTGTTTTTCCTTTGGGCCATTTATGGGAAAAT |
| NCOA2 | 9 | 7 | TCATGCCAGGGCTGCTTTGTGAGGG | TGTGATGGAAGTTAGAGGGT | TGTGATGGAAGTTAGAGGGT | TGTGATGGAAGTTAGAGGGTTTTTGTGATGGAAGTTAGAGGGTTTTTCATGCCAGGGCTGCTTTGTGAGGG |
| NCOA2 | 10 | 7 | TGAAAGCATGGAGGTGGGCTGTCCT | ATAGGAAATGGTGGTAGTGT | ATAGGAAATGGTGGTAGTGT | ATAGGAAATGGTGGTAGTGTTTTATAGGAAATGGTGGTAGTGTTTTTGAAAGCATGGAGGTGGGCTGTCCT |
| NCOA2 | 11 | 7 | GGAGCTGTTGGTATAACTATGGCTAT | TGTGATGGAAGTTAGAGGGT | TGTGATGGAAGTTAGAGGGT | TGTGATGGAAGTTAGAGGGTTTTTGTGATGGAAGTTAGAGGGTTTTGGAGCTGTTGGTATAACTATGGCTAT |
| NCOA2 | 12 | 7 | GCCAACGATGACCCTAATGAGACCC | ATAGGAAATGGTGGTAGTGT | ATAGGAAATGGTGGTAGTGT | ATAGGAAATGGTGGTAGTGTTTTATAGGAAATGGTGGTAGTGTTTTGCCAACGATGACCCTAATGAGACCC |
| NCOA2 | 13 | 7 | CAAGCTTCCCATCTTGCTGAGTGGG | TGTGATGGAAGTTAGAGGGT | TGTGATGGAAGTTAGAGGGT | TGTGATGGAAGTTAGAGGGTTTTTGTGATGGAAGTTAGAGGGTTTTCAAGCTTCCCATCTTGCTGAGTGGG |
| NCOA2 | 14 | 7 | CCAGTTGTACCTTCAGAGGGCTCCC | ATAGGAAATGGTGGTAGTGT | ATAGGAAATGGTGGTAGTGT | ATAGGAAATGGTGGTAGTGTTTTATAGGAAATGGTGGTAGTGTTTTCCAGTTGTACCTTCAGAGGGCTCCC |
| NCOA2 | 15 | 7 | CCGTCAGCTCTCTCACTGCTCACGG | TGTGATGGAAGTTAGAGGGT | TGTGATGGAAGTTAGAGGGT | TGTGATGGAAGTTAGAGGGTTTTTGTGATGGAAGTTAGAGGGTTTTCCGTCAGCTCTCTCACTGCTCACGG |
| NCOA2 | 16 | 7 | GCCCTTTGCTGTCATGCAGTCTGCT | ATAGGAAATGGTGGTAGTGT | ATAGGAAATGGTGGTAGTGT | ATAGGAAATGGTGGTAGTGTTTTATAGGAAATGGTGGTAGTGTTTTGCCCTTTGCTGTCATGCAGTCTGCT |
| NCOA2 | 17 | 7 | ACCCAGAACCAGGCAAGCTACCTGT | TGTGATGGAAGTTAGAGGGT | TGTGATGGAAGTTAGAGGGT | TGTGATGGAAGTTAGAGGGTTTTTGTGATGGAAGTTAGAGGGTTTTACCCAGAACCAGGCAAGCTACCTGT |
| NCOA2 | 18 | 7 | CTTCTCCTTGAGCGAGGTTCCATGT | ATAGGAAATGGTGGTAGTGT | ATAGGAAATGGTGGTAGTGT | ATAGGAAATGGTGGTAGTGTTTTATAGGAAATGGTGGTAGTGTTTTCTTCTCCTTGAGCGAGGTTCCATGT |
| NCOA2 | 19 | 7 | TGTTAACTTGGCCAAGTCCACAGGG | TGTGATGGAAGTTAGAGGGT | TGTGATGGAAGTTAGAGGGT | TGTGATGGAAGTTAGAGGGTTTTTGTGATGGAAGTTAGAGGGTTTTTGTTAACTTGGCCAAGTCCACAGGG |
| NCOA2 | 20 | 7 | TGGCTCAGGTCTTTGCCTGTGGCTT | ATAGGAAATGGTGGTAGTGT | ATAGGAAATGGTGGTAGTGT | ATAGGAAATGGTGGTAGTGTTTTATAGGAAATGGTGGTAGTGTTTTTGGCTCAGGTCTTTGCCTGTGGCTT |
| NCOA2 | 21 | 7 | GCGAAGTAGTGCATTCTCTTTCTTCT | TGTGATGGAAGTTAGAGGGT | TGTGATGGAAGTTAGAGGGT | TGTGATGGAAGTTAGAGGGTTTTTGTGATGGAAGTTAGAGGGTTTTGCGAAGTAGTGCATTCTCTTTCTTCT |
| NCOA2 | 22 | 7 | TACTGTCCAGTCTCTCAAGTTTGGG | ATAGGAAATGGTGGTAGTGT | ATAGGAAATGGTGGTAGTGT | ATAGGAAATGGTGGTAGTGTTTTATAGGAAATGGTGGTAGTGTTTTTACTGTCCAGTCTCTCAAGTTTGGG |
| NCOA2 | 23 | 7 | GCTCATCTCCTCCTTCTCAGTTTTC | TGTGATGGAAGTTAGAGGGT | TGTGATGGAAGTTAGAGGGT | TGTGATGGAAGTTAGAGGGTTTTTGTGATGGAAGTTAGAGGGTTTTGCTCATCTCCTCCTTCTCAGTTTTC |
| NCOA2 | 24 | 7 | CTATTCTGCAAATCATCCAAAATCTCCT | ATAGGAAATGGTGGTAGTGT | ATAGGAAATGGTGGTAGTGT | ATAGGAAATGGTGGTAGTGTTTTATAGGAAATGGTGGTAGTGTTTTCTATTCTGCAAATCATCCAAAATCTCCT |
| NCOA2 | 25 | 7 | TTGCTTGTCAACTGATCCAGCAGGG | TGTGATGGAAGTTAGAGGGT | TGTGATGGAAGTTAGAGGGT | TGTGATGGAAGTTAGAGGGTTTTTGTGATGGAAGTTAGAGGGTTTTTTGCTTGTCAACTGATCCAGCAGGG |
| NCOA2 | 26 | 7 | GTGAGTTGCATGAGGTCATTGATGA | ATAGGAAATGGTGGTAGTGT | ATAGGAAATGGTGGTAGTGT | ATAGGAAATGGTGGTAGTGTTTTATAGGAAATGGTGGTAGTGTTTTGTGAGTTGCATGAGGTCATTGATGA |
| NCOA2 | 27 | 7 | TCTGGTTTGGCAATAACCTGCCCAG | TGTGATGGAAGTTAGAGGGT | TGTGATGGAAGTTAGAGGGT | TGTGATGGAAGTTAGAGGGTTTTTGTGATGGAAGTTAGAGGGTTTTTCTGGTTTGGCAATAACCTGCCCAG |
| NCOA2 | 28 | 7 | GCTTTGCAATGTGATGTCAAGTGGT | ATAGGAAATGGTGGTAGTGT | ATAGGAAATGGTGGTAGTGT | ATAGGAAATGGTGGTAGTGTTTTATAGGAAATGGTGGTAGTGTTTTGCTTTGCAATGTGATGTCAAGTGGT |
| NCOA2 | 29 | 7 | TCCCTTGATTACCCATCATTCCTGG | TGTGATGGAAGTTAGAGGGT | TGTGATGGAAGTTAGAGGGT | TGTGATGGAAGTTAGAGGGTTTTTGTGATGGAAGTTAGAGGGTTTTTCCCTTGATTACCCATCATTCCTGG |
| NCOA2 | 30 | 7 | TGACTCTCACAGCCGAACTCTGCGG | ATAGGAAATGGTGGTAGTGT | ATAGGAAATGGTGGTAGTGT | ATAGGAAATGGTGGTAGTGTTTTATAGGAAATGGTGGTAGTGTTTTTGACTCTCACAGCCGAACTCTGCGG |
| NCOA2 | 31 | 7 | TTCCGAATCATACCTCCTTGGACTG | TGTGATGGAAGTTAGAGGGT | TGTGATGGAAGTTAGAGGGT | TGTGATGGAAGTTAGAGGGTTTTTGTGATGGAAGTTAGAGGGTTTTTTCCGAATCATACCTCCTTGGACTG |
| NCOA2 | 32 | 7 | CTGAAGCGTCTGTCTTTGGCCAGGC | ATAGGAAATGGTGGTAGTGT | ATAGGAAATGGTGGTAGTGT | ATAGGAAATGGTGGTAGTGTTTTATAGGAAATGGTGGTAGTGTTTTCTGAAGCGTCTGTCTTTGGCCAGGC |
| NCOA2 | 33 | 7 | TAAGGCTCTATCAATCTCCTCCAGG | TGTGATGGAAGTTAGAGGGT | TGTGATGGAAGTTAGAGGGT | TGTGATGGAAGTTAGAGGGTTTTTGTGATGGAAGTTAGAGGGTTTTTAAGGCTCTATCAATCTCCTCCAGG |
| NCOA2 | 34 | 7 | CCTTCTGCTCCAGCATGATGTTGGA | ATAGGAAATGGTGGTAGTGT | ATAGGAAATGGTGGTAGTGT | ATAGGAAATGGTGGTAGTGTTTTATAGGAAATGGTGGTAGTGTTTTCCTTCTGCTCCAGCATGATGTTGGA |
| NCOA2 | 35 | 7 | GAATAGCTACCCTGGGCCATTTGTG | TGTGATGGAAGTTAGAGGGT | TGTGATGGAAGTTAGAGGGT | TGTGATGGAAGTTAGAGGGTTTTTGTGATGGAAGTTAGAGGGTTTTGAATAGCTACCCTGGGCCATTTGTG |

| | | | | | | |
|---|---|---|---|---|---|---|
| NCOA2 | 36 | 7 | TGGTGTGAAAGTTTGGATCTTGCAT | ATAGGAATGGTGGTAGTGT | ATAGGAATGGTGGTAGTGT | ATAGGAATGGTGGTAGTGTTTTATAGGAATGGTGGTAGTGTTTTTGGTGTGAAAGTTTGGATCTTGCAT |
| NCOA2 | 37 | 7 | AAGATGCTGGTTCAGGATTTCCCTC | TGTGATGGAAGTTAGAGGGT | TGTGATGGAAGTTAGAGGGT | TGTGATGGAAGTTAGAGGGTTTTTGTGATGGAAGTTAGAGGGTTTTAAGATGCTGGTTCAGGATTTCCCTC |
| NCOA2 | 38 | 7 | TGCTGTTGCTGATGCATTTGTCTCT | ATAGGAATGGTGGTAGTGT | ATAGGAATGGTGGTAGTGT | ATAGGAATGGTGGTAGTGTTTTATAGGAATGGTGGTAGTGTTTTTGCTGTTGCTGATGCATTTGTCTCT |
| NCOA2 | 39 | 7 | CATGCTTGGTGTCATATTCAACCCT | TGTGATGGAAGTTAGAGGGT | TGTGATGGAAGTTAGAGGGT | TGTGATGGAAGTTAGAGGGTTTTTGTGATGGAAGTTAGAGGGTTTTCATGCTTGGTGTCATATTCAACCCT |
| NCOA2 | 40 | 7 | ATAGTTGCTGGCATACCACTAGGAG | ATAGGAATGGTGGTAGTGT | ATAGGAATGGTGGTAGTGT | ATAGGAATGGTGGTAGTGTTTTATAGGAATGGTGGTAGTGTTTTATAGTTGCTGGCATACCACTAGGAG |
| NCOA2 | 41 | 7 | GGACTCTGTGTATGTGCCATTCGGG | TGTGATGGAAGTTAGAGGGT | TGTGATGGAAGTTAGAGGGT | TGTGATGGAAGTTAGAGGGTTTTTGTGATGGAAGTTAGAGGGTTTTGGACTCTGTGTATGTGCCATTCGGG |
| NCOA2 | 42 | 7 | CTGGGTTGGCCTGAGACTGTTGCAT | ATAGGAATGGTGGTAGTGT | ATAGGAATGGTGGTAGTGT | ATAGGAATGGTGGTAGTGTTTTATAGGAATGGTGGTAGTGTTTTCTGGGTTGGCCTGAGACTGTTGCAT |
| NCOA2 | 43 | 7 | CTGCTCATGCCACCTGTGTTGGTCG | TGTGATGGAAGTTAGAGGGT | TGTGATGGAAGTTAGAGGGT | TGTGATGGAAGTTAGAGGGTTTTTGTGATGGAAGTTAGAGGGTTTTCTGCTCATGCCACCTGTGTTGGTCG |
| NCOA2 | 44 | 7 | TGCTGATCTGTCCTGTCATCTGGTT | ATAGGAATGGTGGTAGTGT | ATAGGAATGGTGGTAGTGT | ATAGGAATGGTGGTAGTGTTTTATAGGAATGGTGGTAGTGTTTTTGCTGATCTGTCCTGTCATCTGGTT |
| NCOA2 | 45 | 7 | ACTGGAAGTGTTTTGAGCAAGTGAG | TGTGATGGAAGTTAGAGGGT | TGTGATGGAAGTTAGAGGGT | TGTGATGGAAGTTAGAGGGTTTTTGTGATGGAAGTTAGAGGGTTTTACTGGAAGTGTTTTGAGCAAGTGAG |
| NCOA2 | 46 | 7 | GGGTTGAAACAAATAGACACAGCTCT | ATAGGAATGGTGGTAGTGT | ATAGGAATGGTGGTAGTGT | ATAGGAATGGTGGTAGTGTTTTATAGGAATGGTGGTAGTGTTTTGGGTTGAAACAAATAGACACAGCTCT |
| NCOA2 | 47 | 7 | CTGTCTGGATGCGAGCTTCAGACTG | TGTGATGGAAGTTAGAGGGT | TGTGATGGAAGTTAGAGGGT | TGTGATGGAAGTTAGAGGGTTTTTGTGATGGAAGTTAGAGGGTTTTCTGTCTGGATGCGAGCTTCAGACTG |
| NCOA2 | 48 | 7 | AAGGTGAATGCAGTGAACAGACTGA | ATAGGAATGGTGGTAGTGT | ATAGGAATGGTGGTAGTGT | ATAGGAATGGTGGTAGTGTTTTATAGGAATGGTGGTAGTGTTTTAAGGTGAATGCAGTGAACAGACTGA |
| NCOA2 | 49 | 7 | TGACATGGGTATGAAATTTGCCTGT | TGTGATGGAAGTTAGAGGGT | TGTGATGGAAGTTAGAGGGT | TGTGATGGAAGTTAGAGGGTTTTTGTGATGGAAGTTAGAGGGTTTTTGACATGGGTATGAAATTTGCCTGT |
| NCOA2 | 50 | 7 | ACAAGAGCATGGTTCTCCTTAAATACATA | ATAGGAATGGTGGTAGTGT | ATAGGAATGGTGGTAGTGT | ATAGGAATGGTGGTAGTGTTTTATAGGAATGGTGGTAGTGTTTTACAAGAGCATGGTTCTCCTTAAATACATA |
| NCOA2 | 51 | 7 | CCCAAATTCAGGCTTTAGCTTAAAACAT | TGTGATGGAAGTTAGAGGGT | TGTGATGGAAGTTAGAGGGT | TGTGATGGAAGTTAGAGGGTTTTTGTGATGGAAGTTAGAGGGTTTTCCCAAATTCAGGCTTTAGCTTAAAACAT |
| NCOA2 | 52 | 7 | GCTGTCCACGGTGTCTGTCCTGCTT | ATAGGAATGGTGGTAGTGT | ATAGGAATGGTGGTAGTGT | ATAGGAATGGTGGTAGTGTTTTATAGGAATGGTGGTAGTGTTTTGCTGTCCACGGTGTCTGTCCTGCTT |
| NCOA2 | 53 | 7 | AGAGATACGACTGTGACTTTGTTGG | TGTGATGGAAGTTAGAGGGT | TGTGATGGAAGTTAGAGGGT | TGTGATGGAAGTTAGAGGGTTTTTGTGATGGAAGTTAGAGGGTTTTAGAGATACGACTGTGACTTTGTTGG |
| NCOA2 | 54 | 7 | TCTTTCCAACATGGTCTTTAGAGCT | ATAGGAATGGTGGTAGTGT | ATAGGAATGGTGGTAGTGT | ATAGGAATGGTGGTAGTGTTTTATAGGAATGGTGGTAGTGTTTTTCTTTCCAACATGGTCTTTAGAGCT |
| NCOA2 | 55 | 7 | AATATTTGCTAATGTTAACAGCCCTCTC | TGTGATGGAAGTTAGAGGGT | TGTGATGGAAGTTAGAGGGT | TGTGATGGAAGTTAGAGGGTTTTTGTGATGGAAGTTAGAGGGTTTTAATATTTGCTAATGTTAACAGCCCTCTC |
| NCOA2 | 56 | 7 | AGGTGAACCAGTTTGGTTTTAACAAA | ATAGGAATGGTGGTAGTGT | ATAGGAATGGTGGTAGTGT | ATAGGAATGGTGGTAGTGTTTTATAGGAATGGTGGTAGTGTTTTAGGTGAACCAGTTTGGTTTTAACAAA |
| NCOA2 | 57 | 7 | CTGACAGGGAAGAAGAGATGCTGAT | TGTGATGGAAGTTAGAGGGT | TGTGATGGAAGTTAGAGGGT | TGTGATGGAAGTTAGAGGGTTTTTGTGATGGAAGTTAGAGGGTTTTCTGACAGGGAAGAAGAGATGCTGAT |
| NCOA2 | 58 | 7 | GGCTTACCTGGTTTCTGTAGCTTATATT | ATAGGAATGGTGGTAGTGT | ATAGGAATGGTGGTAGTGT | ATAGGAATGGTGGTAGTGTTTTATAGGAATGGTGGTAGTGTTTTGGCTTACCTGGTTTCTGTAGCTTATATT |
| NCOA2 | 59 | 7 | GCTCGATTGGTATCAAGCCTTAACTTT | TGTGATGGAAGTTAGAGGGT | TGTGATGGAAGTTAGAGGGT | TGTGATGGAAGTTAGAGGGTTTTTGTGATGGAAGTTAGAGGGTTTTGCTCGATTGGTATCAAGCCTTAACTTT |
| NCOA2 | 60 | 7 | GCTATCAATTTAAACACACTGGCGTTTT | ATAGGAATGGTGGTAGTGT | ATAGGAATGGTGGTAGTGT | ATAGGAATGGTGGTAGTGTTTTATAGGAATGGTGGTAGTGTTTTGCTATCAATTTAAACACACTGGCGTTTT |
| NCOA2 | 61 | 7 | TTCGGCTTAGTCAGCACTGCTAAAA | TGTGATGGAAGTTAGAGGGT | TGTGATGGAAGTTAGAGGGT | TGTGATGGAAGTTAGAGGGTTTTTGTGATGGAAGTTAGAGGGTTTTTTCGGCTTAGTCAGCACTGCTAAAA |
| NCOA2 | 62 | 7 | CCTGCTTAGTAACATGTAGTCTTTCTAAGG | ATAGGAATGGTGGTAGTGT | ATAGGAATGGTGGTAGTGT | ATAGGAATGGTGGTAGTGTTTTATAGGAATGGTGGTAGTGTTTTCCTGCTTAGTAACATGTAGTCTTTCTAAGG |
| NCOA2 | 63 | 7 | CTTTGGGCTATATCTCTTCCTATATTTCCA | TGTGATGGAAGTTAGAGGGT | TGTGATGGAAGTTAGAGGGT | TGTGATGGAAGTTAGAGGGTTTTTGTGATGGAAGTTAGAGGGTTTTCTTTGGGCTATATCTCTTCCTATATTTCCA |
| NCOA2 | 64 | 7 | GCAACATTTTCAGACTGTCTACTTACCAT | ATAGGAATGGTGGTAGTGT | ATAGGAATGGTGGTAGTGT | ATAGGAATGGTGGTAGTGTTTTATAGGAATGGTGGTAGTGTTTTGCAACATTTTCAGACTGTCTACTTACCAT |
| NCOA2 | 65 | 7 | ACTGGTGGCCACTGAGAAGTGTTGT | TGTGATGGAAGTTAGAGGGT | TGTGATGGAAGTTAGAGGGT | TGTGATGGAAGTTAGAGGGTTTTTGTGATGGAAGTTAGAGGGTTTTACTGGTGGCCACTGAGAAGTGTTGT |
| NCOA2 | 66 | 7 | GGAGAAAAGTAGAGCCTTTTGAGGG | ATAGGAATGGTGGTAGTGT | ATAGGAATGGTGGTAGTGT | ATAGGAATGGTGGTAGTGTTTTATAGGAATGGTGGTAGTGTTTTGGAGAAAAGTAGAGCCTTTTGAGGG |
| NCOA2 | 67 | 7 | TTGTATGCCACATGAGCCTCAGCCC | TGTGATGGAAGTTAGAGGGT | TGTGATGGAAGTTAGAGGGT | TGTGATGGAAGTTAGAGGGTTTTTGTGATGGAAGTTAGAGGGTTTTTTGTATGCCACATGAGCCTCAGCCC |
| NCOA2 | 68 | 7 | GTGCTGTGTGAAGTTAAAAGTCATTCTA | ATAGGAATGGTGGTAGTGT | ATAGGAATGGTGGTAGTGT | ATAGGAATGGTGGTAGTGTTTTATAGGAATGGTGGTAGTGTTTTGTGCTGTGTGAAGTTAAAAGTCATTCTA |
| NCOA2 | 69 | 7 | AGATAGTTTTGGACGGTGTTGTAGTG | TGTGATGGAAGTTAGAGGGT | TGTGATGGAAGTTAGAGGGT | TGTGATGGAAGTTAGAGGGTTTTTGTGATGGAAGTTAGAGGGTTTTAGATAGTTTTGGACGGTGTTGTAGTG |
| NCOA2 | 70 | 7 | AAATACAGCGGGCAGATGGATAAAA | ATAGGAATGGTGGTAGTGT | ATAGGAATGGTGGTAGTGT | ATAGGAATGGTGGTAGTGTTTTATAGGAATGGTGGTAGTGTTTTAAATACAGCGGGCAGATGGATAAAA |
| NCOA2 | 71 | 7 | CCTTCAAAACAGCAATGATCACGAA | TGTGATGGAAGTTAGAGGGT | TGTGATGGAAGTTAGAGGGT | TGTGATGGAAGTTAGAGGGTTTTTGTGATGGAAGTTAGAGGGTTTTCCTTCAAAACAGCAATGATCACGAA |
| NCOA2 | 72 | 7 | ACCTGATTGAAACCGTAAGAGAATTTATC | ATAGGAATGGTGGTAGTGT | ATAGGAATGGTGGTAGTGT | ATAGGAATGGTGGTAGTGTTTTATAGGAATGGTGGTAGTGTTTTACCTGATTGAAACCGTAAGAGAATTTATC |
| NCOA2 | 73 | 7 | CAGGCCATAAGAGATACACATAGGG | TGTGATGGAAGTTAGAGGGT | TGTGATGGAAGTTAGAGGGT | TGTGATGGAAGTTAGAGGGTTTTTGTGATGGAAGTTAGAGGGTTTTCAGGCCATAAGAGATACACATAGGG |
| NCOA2 | 74 | 7 | TGCAAGTGGATCTGTGCCACCCATT | ATAGGAATGGTGGTAGTGT | ATAGGAATGGTGGTAGTGT | ATAGGAATGGTGGTAGTGTTTTATAGGAATGGTGGTAGTGTTTTTGCAAGTGGATCTGTGCCACCCATT |
| NCOA2 | 75 | 7 | TTGTGCAGAGGATGCTTTGCTCTTA | TGTGATGGAAGTTAGAGGGT | TGTGATGGAAGTTAGAGGGT | TGTGATGGAAGTTAGAGGGTTTTTGTGATGGAAGTTAGAGGGTTTTTTGTGCAGAGGATGCTTTGCTCTTA |
| NCOA2 | 76 | 7 | GTCTATAAAGAAACACCAGGGAGAAAATTC | ATAGGAATGGTGGTAGTGT | ATAGGAATGGTGGTAGTGT | ATAGGAATGGTGGTAGTGTTTTATAGGAATGGTGGTAGTGTTTTGTCTATAAAGAAACACCAGGGAGAAAATTC |
| NCOA2 | 77 | 7 | GGACAGCTATGTAGGAATATACAAGACT | TGTGATGGAAGTTAGAGGGT | TGTGATGGAAGTTAGAGGGT | TGTGATGGAAGTTAGAGGGTTTTTGTGATGGAAGTTAGAGGGTTTTGGACAGCTATGTAGGAATATACAAGACT |
| NCOA2 | 78 | 7 | AGCTTTCTCCTGAAGAACCATAGTTAA | ATAGGAATGGTGGTAGTGT | ATAGGAATGGTGGTAGTGT | ATAGGAATGGTGGTAGTGTTTTATAGGAATGGTGGTAGTGTTTTAGCTTTCTCCTGAAGAACCATAGTTAA |
| NCOA2 | 79 | 7 | AATACAAACATCAATGCAGGTGAACA | TGTGATGGAAGTTAGAGGGT | TGTGATGGAAGTTAGAGGGT | TGTGATGGAAGTTAGAGGGTTTTTGTGATGGAAGTTAGAGGGTTTTAATACAAACATCAATGCAGGTGAACA |
| NCOA2 | 80 | 7 | CTACCTTTGCATAAACTGCTCACAC | ATAGGAATGGTGGTAGTGT | ATAGGAATGGTGGTAGTGT | ATAGGAATGGTGGTAGTGTTTTATAGGAATGGTGGTAGTGTTTTCTACCTTTGCATAAACTGCTCACAC |
| BRCA2 | 1 | 8 | GTGGAGTTTTAAATAGGTTTGGTTCGT | TGTGGAGGGATTGAAGGATA | TGTGGAGGGATTGAAGGATA | TGTGGAGGGATTGAAGGATATTTTGTGGAGGGATTGAAGGATATTTGTGGAGTTTTAAATAGGTTTGGTTCGT |
| BRCA2 | 2 | 8 | AGCCAGCTGATTATAAGATGGTTTCC | AGAGTGAGTAGTAGTGGAGT | AGAGTGAGTAGTAGTGGAGT | AGAGTGAGTAGTAGTGGAGTTTTAGAGTGAGTAGTAGTGGAGTTTTAGCCAGCTGATTATAAGATGGTTTCC |
| BRCA2 | 3 | 8 | TGGTTCCACTTCAGATACAAATGAGTA | TGTGGAGGGATTGAAGGATA | TGTGGAGGGATTGAAGGATA | TGTGGAGGGATTGAAGGATATTTTGTGGAGGGATTGAAGGATATTTTGGTTCCACTTCAGATACAAATGAGTA |
| BRCA2 | 4 | 8 | GGCTTCTGATTTGCTACATTTGAATCTAA | AGAGTGAGTAGTAGTGGAGT | AGAGTGAGTAGTAGTGGAGT | AGAGTGAGTAGTAGTGGAGTTTTAGAGTGAGTAGTAGTGGAGTTTTGGCTTCTGATTTGCTACATTTGAATCTAA |
| BRCA2 | 5 | 8 | TTCACAGGCCAAAGACGGTACAACTT | TGTGGAGGGATTGAAGGATA | TGTGGAGGGATTGAAGGATA | TGTGGAGGGATTGAAGGATATTTTGTGGAGGGATTGAAGGATATTTTTCACAGGCCAAAGACGGTACAACTT |
| BRCA2 | 6 | 8 | ATTTAGACCTGAAAGGGTTAGTTGAGA | AGAGTGAGTAGTAGTGGAGT | AGAGTGAGTAGTAGTGGAGT | AGAGTGAGTAGTAGTGGAGTTTTAGAGTGAGTAGTAGTGGAGTTTTATTTAGACCTGAAAGGGTTAGTTGAGA |
| BRCA2 | 7 | 8 | GTGGCAAAGAATTCTCTGAAGTAAGAA | TGTGGAGGGATTGAAGGATA | TGTGGAGGGATTGAAGGATA | TGTGGAGGGATTGAAGGATATTTTGTGGAGGGATTGAAGGATATTTGTGGCAAAGAATTCTCTGAAGTAAGAA |
| BRCA2 | 8 | 8 | GCTTCTCTGATTTTGGTAGGCTAGAAA | AGAGTGAGTAGTAGTGGAGT | AGAGTGAGTAGTAGTGGAGT | AGAGTGAGTAGTAGTGGAGTTTTAGAGTGAGTAGTAGTGGAGTTTTGCTTCTCTGATTTTGGTAGGCTAGAAA |
| BRCA2 | 9 | 8 | ATATTGCCTGCTTTACTGCAAGAATG | TGTGGAGGGATTGAAGGATA | TGTGGAGGGATTGAAGGATA | TGTGGAGGGATTGAAGGATATTTTGTGGAGGGATTGAAGGATATTTATATTGCCTGCTTTACTGCAAGAATG |
| BRCA2 | 10 | 8 | AGTTTGGATCAGTCATATGACCTGAAAA | AGAGTGAGTAGTAGTGGAGT | AGAGTGAGTAGTAGTGGAGT | AGAGTGAGTAGTAGTGGAGTTTTAGAGTGAGTAGTAGTGGAGTTTTAGTTTGGATCAGTCATATGACCTGAAAA |
| BRCA2 | 11 | 8 | CCAGCTTCCATTATCAATTAAATTTGGAC | TGTGGAGGGATTGAAGGATA | TGTGGAGGGATTGAAGGATA | TGTGGAGGGATTGAAGGATATTTTGTGGAGGGATTGAAGGATATTTCCAGCTTCCATTATCAATTAAATTTGGAC |
| BRCA2 | 12 | 8 | GGGCTGAACAGTTAATTAGTTCTGATTTT | AGAGTGAGTAGTAGTGGAGT | AGAGTGAGTAGTAGTGGAGT | AGAGTGAGTAGTAGTGGAGTTTTAGAGTGAGTAGTAGTGGAGTTTTGGGCTGAACAGTTAATTAGTTCTGATTTT |
| BRCA2 | 13 | 8 | TTGAATGTTGTACTGGGTGACATGCT | TGTGGAGGGATTGAAGGATA | TGTGGAGGGATTGAAGGATA | TGTGGAGGGATTGAAGGATATTTTGTGGAGGGATTGAAGGATATTTTTGAATGTTGTACTGGGTGACATGCT |
| BRCA2 | 14 | 8 | GCTGGCATTTTCATGATCATATAAAAGAC | AGAGTGAGTAGTAGTGGAGT | AGAGTGAGTAGTAGTGGAGT | AGAGTGAGTAGTAGTGGAGTTTTAGAGTGAGTAGTAGTGGAGTTTTGCTGGCATTTTCATGATCATATAAAAGAC |
| BRCA2 | 15 | 8 | TTGCCTCTAGAAATCATGACTAGGTTT | TGTGGAGGGATTGAAGGATA | TGTGGAGGGATTGAAGGATA | TGTGGAGGGATTGAAGGATATTTTGTGGAGGGATTGAAGGATATTTTTGCCTCTAGAAATCATGACTAGGTTT |
| BRCA2 | 16 | 8 | CTTTGAGCTTGTCTGACATTTTGTATGAT | AGAGTGAGTAGTAGTGGAGT | AGAGTGAGTAGTAGTGGAGT | AGAGTGAGTAGTAGTGGAGTTTTAGAGTGAGTAGTAGTGGAGTTTTCTTTGAGCTTGTCTGACATTTTGTATGAT |
| BRCA2 | 17 | 8 | CTTGAAGGTGATGCTACTCTCATGTA | TGTGGAGGGATTGAAGGATA | TGTGGAGGGATTGAAGGATA | TGTGGAGGGATTGAAGGATATTTTGTGGAGGGATTGAAGGATATTTCTTGAAGGTGATGCTACTCTCATGTA |
| BRCA2 | 18 | 8 | CCTTTCATTAGCTACTTGGAAGACAAAAT | AGAGTGAGTAGTAGTGGAGT | AGAGTGAGTAGTAGTGGAGT | AGAGTGAGTAGTAGTGGAGTTTTAGAGTGAGTAGTAGTGGAGTTTTCCTTTCATTAGCTACTTGGAAGACAAAAT |
| BRCA2 | 19 | 8 | TTGAAAATGGGTTCGTTTACACAAGT | TGTGGAGGGATTGAAGGATA | TGTGGAGGGATTGAAGGATA | TGTGGAGGGATTGAAGGATATTTTGTGGAGGGATTGAAGGATATTTTTGAAAATGGGTTCGTTTACACAAGT |
| BRCA2 | 20 | 8 | TGTGTCTCCATATAAAACCATGGTAGAG | AGAGTGAGTAGTAGTGGAGT | AGAGTGAGTAGTAGTGGAGT | AGAGTGAGTAGTAGTGGAGTTTTAGAGTGAGTAGTAGTGGAGTTTTTGTGTCTCCATATAAAACCATGGTAGAG |
| BRCA2 | 21 | 8 | CCTGCTTGGAAAATAACATCTGAGGG | TGTGGAGGGATTGAAGGATA | TGTGGAGGGATTGAAGGATA | TGTGGAGGGATTGAAGGATATTTTGTGGAGGGATTGAAGGATATTTCCTGCTTGGAAAATAACATCTGAGGG |
| BRCA2 | 22 | 8 | CCTTTTGGCTAGGTGTTAAATTATGGTT | AGAGTGAGTAGTAGTGGAGT | AGAGTGAGTAGTAGTGGAGT | AGAGTGAGTAGTAGTGGAGTTTTAGAGTGAGTAGTAGTGGAGTTTTCCTTTTGGCTAGGTGTTAAATTATGGTT |
| BRCA2 | 23 | 8 | CTTCTGCAATATGTAGCTTGGTTTTCTAA | TGTGGAGGGATTGAAGGATA | TGTGGAGGGATTGAAGGATA | TGTGGAGGGATTGAAGGATATTTTGTGGAGGGATTGAAGGATATTTCTTCTGCAATATGTAGCTTGGTTTTCTAA |
| BRCA2 | 24 | 8 | ATCTGGTTTTCAGGCACTTCAAATGT | AGAGTGAGTAGTAGTGGAGT | AGAGTGAGTAGTAGTGGAGT | AGAGTGAGTAGTAGTGGAGTTTTAGAGTGAGTAGTAGTGGAGTTTTATCTGGTTTTCAGGCACTTCAAATGT |
| BRCA2 | 25 | 8 | GCATTCATTATGACATGAAGATCAGCATC | TGTGGAGGGATTGAAGGATA | TGTGGAGGGATTGAAGGATA | TGTGGAGGGATTGAAGGATATTTTGTGGAGGGATTGAAGGATATTTGCATTCATTATGACATGAAGATCAGCATC |
| BRCA2 | 26 | 8 | TGCTGCTGTCTACCTGACCAATCGAT | AGAGTGAGTAGTAGTGGAGT | AGAGTGAGTAGTAGTGGAGT | AGAGTGAGTAGTAGTGGAGTTTTAGAGTGAGTAGTAGTGGAGTTTTTGCTGCTGTCTACCTGACCAATCGAT |
| BRCA2 | 27 | 8 | CACTGCCATCAAATTCTAAGTTATGAGAA | TGTGGAGGGATTGAAGGATA | TGTGGAGGGATTGAAGGATA | TGTGGAGGGATTGAAGGATATTTTGTGGAGGGATTGAAGGATATTTCACTGCCATCAAATTCTAAGTTATGAGAA |
| BRCA2 | 28 | 8 | CAGTAAATAGCAAGTCCGTTTCATCTT | AGAGTGAGTAGTAGTGGAGT | AGAGTGAGTAGTAGTGGAGT | AGAGTGAGTAGTAGTGGAGTTTTAGAGTGAGTAGTAGTGGAGTTTTCAGTAAATAGCAAGTCCGTTTCATCTT |
| BRCA2 | 29 | 8 | ATCTGAGTGTTTCCCTCCTTCATAAA | TGTGGAGGGATTGAAGGATA | TGTGGAGGGATTGAAGGATA | TGTGGAGGGATTGAAGGATATTTTGTGGAGGGATTGAAGGATATTTATCTGAGTGTTTCCCTCCTTCATAAA |
| BRCA2 | 30 | 8 | TTCTTGAGCTTTCGCAACTTCCAAAA | AGAGTGAGTAGTAGTGGAGT | AGAGTGAGTAGTAGTGGAGT | AGAGTGAGTAGTAGTGGAGTTTTAGAGTGAGTAGTAGTGGAGTTTTTTCTTGAGCTTTCGCAACTTCCAAAA |
| BRCA2 | 31 | 8 | GCAATTCTTCTGGTTTCTGATCAAAGAA | TGTGGAGGGATTGAAGGATA | TGTGGAGGGATTGAAGGATA | TGTGGAGGGATTGAAGGATATTTTGTGGAGGGATTGAAGGATATTTGCAATTCTTCTGGTTTCTGATCAAAGAA |
| BRCA2 | 32 | 8 | TCTGTTTCCTCATAACTTAGAATGTCCA | AGAGTGAGTAGTAGTGGAGT | AGAGTGAGTAGTAGTGGAGT | AGAGTGAGTAGTAGTGGAGTTTTAGAGTGAGTAGTAGTGGAGTTTTTCTGTTTCCTCATAACTTAGAATGTCCA |
| BRCA2 | 33 | 8 | CTTTTCATCACGTTCGGGTTGTCCCT | TGTGGAGGGATTGAAGGATA | TGTGGAGGGATTGAAGGATA | TGTGGAGGGATTGAAGGATATTTTGTGGAGGGATTGAAGGATATTTCTTTTCATCACGTTCGGGTTGTCCCT |
| BRCA2 | 34 | 8 | GCTGTATGAAAACCCAATAGAGTAGGTT | AGAGTGAGTAGTAGTGGAGT | AGAGTGAGTAGTAGTGGAGT | AGAGTGAGTAGTAGTGGAGTTTTAGAGTGAGTAGTAGTGGAGTTTTGCTGTATGAAAACCCAATAGAGTAGGTT |
| BRCA2 | 35 | 8 | AGGGTCTTTGCCCATTGATGGCTAAA | TGTGGAGGGATTGAAGGATA | TGTGGAGGGATTGAAGGATA | TGTGGAGGGATTGAAGGATATTTTGTGGAGGGATTGAAGGATATTTAGGGTCTTTGCCCATTGATGGCTAAA |
| BRCA2 | 36 | 8 | CCACAGTCTCAATAGAAACAAGGTTTT | AGAGTGAGTAGTAGTGGAGT | AGAGTGAGTAGTAGTGGAGT | AGAGTGAGTAGTAGTGGAGTTTTAGAGTGAGTAGTAGTGGAGTTTTCCACAGTCTCAATAGAAACAAGGTTTT |
| BRCA2 | 37 | 8 | GGCTGAATTTTCAATGACTGAATAAGGG | TGTGGAGGGATTGAAGGATA | TGTGGAGGGATTGAAGGATA | TGTGGAGGGATTGAAGGATATTTTGTGGAGGGATTGAAGGATATTTGGCTGAATTTTCAATGACTGAATAAGGG |
| BRCA2 | 38 | 8 | GTTCCTCAACGCAAATATCTTCATTTACA | AGAGTGAGTAGTAGTGGAGT | AGAGTGAGTAGTAGTGGAGT | AGAGTGAGTAGTAGTGGAGTTTTAGAGTGAGTAGTAGTGGAGTTTTGTTCCTCAACGCAAATATCTTCATTTACA |
| BRCA2 | 39 | 8 | ACGATTTTACCACTGGCTATCCTAAAT | TGTGGAGGGATTGAAGGATA | TGTGGAGGGATTGAAGGATA | TGTGGAGGGATTGAAGGATATTTTGTGGAGGGATTGAAGGATATTTACGATTTTACCACTGGCTATCCTAAAT |
| BRCA2 | 40 | 8 | CCTCTGAATCATCCAATGCCTCGTAA | AGAGTGAGTAGTAGTGGAGT | AGAGTGAGTAGTAGTGGAGT | AGAGTGAGTAGTAGTGGAGTTTTAGAGTGAGTAGTAGTGGAGTTTTCCTCTGAATCATCCAATGCCTCGTAA |
| BRCA2 | 41 | 8 | CTTTCTCCAATCCAGACATATTTTGGTTA | TGTGGAGGGATTGAAGGATA | TGTGGAGGGATTGAAGGATA | TGTGGAGGGATTGAAGGATATTTTGTGGAGGGATTGAAGGATATTTCTTTCTCCAATCCAGACATATTTTGGTTA |
| BRCA2 | 42 | 8 | GTTTCCAAACTAACATCACAAGGTGA | AGAGTGAGTAGTAGTGGAGT | AGAGTGAGTAGTAGTGGAGT | AGAGTGAGTAGTAGTGGAGTTTTAGAGTGAGTAGTAGTGGAGTTTTGTTTCCAAACTAACATCACAAGGTGA |
| BRCA2 | 43 | 8 | GACTTGCTTGGTACTATCTTCTATTTCAG | TGTGGAGGGATTGAAGGATA | TGTGGAGGGATTGAAGGATA | TGTGGAGGGATTGAAGGATATTTTGTGGAGGGATTGAAGGATATTTGACTTGCTTGGTACTATCTTCTATTTCAG |
| BRCA2 | 44 | 8 | GCTGGTCTGAATGTTCGTTACTTTTAAA | AGAGTGAGTAGTAGTGGAGT | AGAGTGAGTAGTAGTGGAGT | AGAGTGAGTAGTAGTGGAGTTTTAGAGTGAGTAGTAGTGGAGTTTTGCTGGTCTGAATGTTCGTTACTTTTAAA |
| BRCA2 | 45 | 8 | AACTTGCTTTCCACTTGCTGTACTAA | TGTGGAGGGATTGAAGGATA | TGTGGAGGGATTGAAGGATA | TGTGGAGGGATTGAAGGATATTTTGTGGAGGGATTGAAGGATATTTAACTTGCTTTCCACTTGCTGTACTAA |
| BRCA2 | 46 | 8 | GTGAAGACTATGCTCAGTTCTGATTAAATC | AGAGTGAGTAGTAGTGGAGT | AGAGTGAGTAGTAGTGGAGT | AGAGTGAGTAGTAGTGGAGTTTTAGAGTGAGTAGTAGTGGAGTTTTGTGAAGACTATGCTCAGTTCTGATTAAATC |
| BRCA2 | 47 | 8 | GGTTTCTCTTATCAACACGAGGAAGTA | TGTGGAGGGATTGAAGGATA | TGTGGAGGGATTGAAGGATA | TGTGGAGGGATTGAAGGATATTTTGTGGAGGGATTGAAGGATATTTGGTTTCTCTTATCAACACGAGGAAGTA |
| BRCA2 | 48 | 8 | TTCCATTTCTGAGTTTACACAGTGCT | AGAGTGAGTAGTAGTGGAGT | AGAGTGAGTAGTAGTGGAGT | AGAGTGAGTAGTAGTGGAGTTTTAGAGTGAGTAGTAGTGGAGTTTTTTCCATTTCTGAGTTTACACAGTGCT |
| BRCA2 | 49 | 8 | CACTTTGGTTCCTAATACCAACTGTTG | TGTGGAGGGATTGAAGGATA | TGTGGAGGGATTGAAGGATA | TGTGGAGGGATTGAAGGATATTTTGTGGAGGGATTGAAGGATATTTCACTTTGGTTCCTAATACCAACTGTTG |
| BRCA2 | 50 | 8 | GTTCATCATCTTCCATAAAAGCTTTAGCA | AGAGTGAGTAGTAGTGGAGT | AGAGTGAGTAGTAGTGGAGT | AGAGTGAGTAGTAGTGGAGTTTTAGAGTGAGTAGTAGTGGAGTTTTGTTCATCATCTTCCATAAAAGCTTTAGCA |
| BRCA2 | 51 | 8 | CATTTCCTCATTTTCGGGACATGTAAAA | TGTGGAGGGATTGAAGGATA | TGTGGAGGGATTGAAGGATA | TGTGGAGGGATTGAAGGATATTTTGTGGAGGGATTGAAGGATATTTCATTTCCTCATTTTCGGGACATGTAAAA |
| BRCA2 | 52 | 8 | CGGTAAAATTTGGATTCTGTATCTCTTGA | AGAGTGAGTAGTAGTGGAGT | AGAGTGAGTAGTAGTGGAGT | AGAGTGAGTAGTAGTGGAGTTTTAGAGTGAGTAGTAGTGGAGTTTTCGGTAAAATTTGGATTCTGTATCTCTTGA |
| BRCA2 | 53 | 8 | CCTGAAACTGCTAAATTGCTTGAAGAT | TGTGGAGGGATTGAAGGATA | TGTGGAGGGATTGAAGGATA | TGTGGAGGGATTGAAGGATATTTTGTGGAGGGATTGAAGGATATTTCCTGAAACTGCTAAATTGCTTGAAGAT |
| BRCA2 | 54 | 8 | TGCCTGTAGTAATCAAGTGTCTCATTTT | AGAGTGAGTAGTAGTGGAGT | AGAGTGAGTAGTAGTGGAGT | AGAGTGAGTAGTAGTGGAGTTTTAGAGTGAGTAGTAGTGGAGTTTTTGCCTGTAGTAATCAAGTGTCTCATTTT |
| BRCA2 | 55 | 8 | CACTGTTCAACTCTGTGAAAATGTGATT | TGTGGAGGGATTGAAGGATA | TGTGGAGGGATTGAAGGATA | TGTGGAGGGATTGAAGGATATTTTGTGGAGGGATTGAAGGATATTTCACTGTTCAACTCTGTGAAAATGTGATT |
| BRCA2 | 56 | 8 | ATCATCAGAGCCATGTCCATCAATGT | AGAGTGAGTAGTAGTGGAGT | AGAGTGAGTAGTAGTGGAGT | AGAGTGAGTAGTAGTGGAGTTTTAGAGTGAGTAGTAGTGGAGTTTTATCATCAGAGCCATGTCCATCAATGT |
| BRCA2 | 57 | 8 | ACAGACTGCCTGGCTGTGGAAAGACG | TGTGGAGGGATTGAAGGATA | TGTGGAGGGATTGAAGGATA | TGTGGAGGGATTGAAGGATATTTTGTGGAGGGATTGAAGGATATTTACAGACTGCCTGGCTGTGGAAAGACG |
| BRCA2 | 58 | 8 | ATCCACCATCAGCCAACTGTATTCCT | AGAGTGAGTAGTAGTGGAGT | AGAGTGAGTAGTAGTGGAGT | AGAGTGAGTAGTAGTGGAGTTTTAGAGTGAGTAGTAGTGGAGTTTTATCCACCATCAGCCAACTGTATTCCT |
| BRCA2 | 59 | 8 | GCTGTGTCATCCCTTTCCATTATCTTTT | TGTGGAGGGATTGAAGGATA | TGTGGAGGGATTGAAGGATA | TGTGGAGGGATTGAAGGATATTTTGTGGAGGGATTGAAGGATATTTGCTGTGTCATCCCTTTCCATTATCTTTT |
| BRCA2 | 60 | 8 | AGAAGTTTCAGATATATTTGCGCTCAATG | AGAGTGAGTAGTAGTGGAGT | AGAGTGAGTAGTAGTGGAGT | AGAGTGAGTAGTAGTGGAGTTTTAGAGTGAGTAGTAGTGGAGTTTTAGAAGTTTCAGATATATTTGCGCTCAATG |
| BRCA2 | 61 | 8 | ATCTAACTGGGCCTTAACAGCATACC | TGTGGAGGGATTGAAGGATA | TGTGGAGGGATTGAAGGATA | TGTGGAGGGATTGAAGGATATTTTGTGGAGGGATTGAAGGATATTTATCTAACTGGGCCTTAACAGCATACC |
| BRCA2 | 62 | 8 | GCCATTCTTTAAGACAGCTAAGAGGG | AGAGTGAGTAGTAGTGGAGT | AGAGTGAGTAGTAGTGGAGT | AGAGTGAGTAGTAGTGGAGTTTTAGAGTGAGTAGTAGTGGAGTTTTGCCATTCTTTAAGACAGCTAAGAGGG |
| BRCA2 | 63 | 8 | TCTAGGGTCAGGAAAGAATCCAAGTT | TGTGGAGGGATTGAAGGATA | TGTGGAGGGATTGAAGGATA | TGTGGAGGGATTGAAGGATATTTTGTGGAGGGATTGAAGGATATTTTCTAGGGTCAGGAAAGAATCCAAGTT |
| BRCA2 | 64 | 8 | AAAAGCGATGATAAGGGCAGAGGAAA | AGAGTGAGTAGTAGTGGAGT | AGAGTGAGTAGTAGTGGAGT | AGAGTGAGTAGTAGTGGAGTTTTAGAGTGAGTAGTAGTGGAGTTTTAAAAGCGATGATAAGGGCAGAGGAAA |
| BRCA2 | 65 | 8 | CAAAGCACGAACTTGCTGTCTTGTTA | TGTGGAGGGATTGAAGGATA | TGTGGAGGGATTGAAGGATA | TGTGGAGGGATTGAAGGATATTTTGTGGAGGGATTGAAGGATATTTCAAAGCACGAACTTGCTGTCTTGTTA |
| BRCA2 | 66 | 8 | CAAGGTAAGCTGGGTCTGCTGCATTC | AGAGTGAGTAGTAGTGGAGT | AGAGTGAGTAGTAGTGGAGT | AGAGTGAGTAGTAGTGGAGTTTTAGAGTGAGTAGTAGTGGAGTTTTCAAGGTAAGCTGGGTCTGCTGCATTC |
| BRCA2 | 67 | 8 | CATGGCCTTCCTAATTTCCAACTGGA | TGTGGAGGGATTGAAGGATA | TGTGGAGGGATTGAAGGATA | TGTGGAGGGATTGAAGGATATTTTGTGGAGGGATTGAAGGATATTTCATGGCCTTCCTAATTTCCAACTGGA |

| | | | | | | |
|---|---|---|---|---|---|---|
| BRCA2 | 68 | 8 | AAACCTTGTTCCTTTTGTTCAGCAGA | AGAGTGAGTAGTAGTGGAGT | AGAGTGAGTAGTAGTGGAGT | AGAGTGAGTAGTAGTGGAGTTTTAGAGTGAGTAGTAGTGGAGTTTTAAACCTTGTTCCTTTTGTTCAGCAGA |
| BRCA2 | 69 | 8 | GGTCTATCCAAAACTTTATTGCCAGTAAA | TGTGGAGGGATTGAAGGATA | TGTGGAGGGATTGAAGGATA | TGTGGAGGGATTGAAGGATATTTTGTGGAGGGATTGAAGGATATTTGGTCTATCCAAAACTTTATTGCCAGTAAA |
| BRCA2 | 70 | 8 | ATTTGGATTCTGGTCGCCACTGGAGG | AGAGTGAGTAGTAGTGGAGT | AGAGTGAGTAGTAGTGGAGT | AGAGTGAGTAGTAGTGGAGTTTTAGAGTGAGTAGTAGTGGAGTTTTATTTGGATTCTGGTCGCCACTGGAGG |
| BRCA2 | 71 | 8 | CTCTTTTGGACTAGCAGAAAACACAG | TGTGGAGGGATTGAAGGATA | TGTGGAGGGATTGAAGGATA | TGTGGAGGGATTGAAGGATATTTTGTGGAGGGATTGAAGGATATTTCTCTTTTGGACTAGCAGAAAACACAG |
| BRCA2 | 72 | 8 | GTTTTCTGCTTCATTGCAAAGTATGTC | AGAGTGAGTAGTAGTGGAGT | AGAGTGAGTAGTAGTGGAGT | AGAGTGAGTAGTAGTGGAGTTTTAGAGTGAGTAGTAGTGGAGTTTTGTTTTCTGCTTCATTGCAAAGTATGTC |
| BRCA2 | 73 | 8 | AGACAGGTGTGGAAACAGACTTCCTT | TGTGGAGGGATTGAAGGATA | TGTGGAGGGATTGAAGGATA | TGTGGAGGGATTGAAGGATATTTTGTGGAGGGATTGAAGGATATTTAGACAGGTGTGGAAACAGACTTCCTT |
| BRCA2 | 74 | 8 | CAGTTCTTTTGGTCATCAATCTCTTTCT | AGAGTGAGTAGTAGTGGAGT | AGAGTGAGTAGTAGTGGAGT | AGAGTGAGTAGTAGTGGAGTTTTAGAGTGAGTAGTAGTGGAGTTTTCAGTTCTTTTGGTCATCAATCTCTTTCT |
| BRCA2 | 75 | 8 | GCAGCCGGAGAAACAAATGTACAAAT | TGTGGAGGGATTGAAGGATA | TGTGGAGGGATTGAAGGATA | TGTGGAGGGATTGAAGGATATTTTGTGGAGGGATTGAAGGATATTTGCAGCCGGAGAAACAAATGTACAAAT |
| BRCA2 | 76 | 8 | CTTTATGGGTGTTTCGTATTTGGTGC | AGAGTGAGTAGTAGTGGAGT | AGAGTGAGTAGTAGTGGAGT | AGAGTGAGTAGTAGTGGAGTTTTAGAGTGAGTAGTAGTGGAGTTTTCTTTATGGGTGTTTCGTATTTGGTGC |
| BRCA2 | 77 | 8 | TTCTTCGTCAGCTATTGAATTACTTTCC | TGTGGAGGGATTGAAGGATA | TGTGGAGGGATTGAAGGATA | TGTGGAGGGATTGAAGGATATTTTGTGGAGGGATTGAAGGATATTTTTCTTCGTCAGCTATTGAATTACTTTCC |
| BRCA2 | 78 | 8 | ACAAAAGAGCTTGGGTATTTATCAATGC | AGAGTGAGTAGTAGTGGAGT | AGAGTGAGTAGTAGTGGAGT | AGAGTGAGTAGTAGTGGAGTTTTAGAGTGAGTAGTAGTGGAGTTTTACAAAAGAGCTTGGGTATTTATCAATGC |
| BRCA2 | 79 | 8 | GAGATGTAGTACAACGTCGTTTCAGT | TGTGGAGGGATTGAAGGATA | TGTGGAGGGATTGAAGGATA | TGTGGAGGGATTGAAGGATATTTTGTGGAGGGATTGAAGGATATTTGAGATGTAGTACAACGTCGTTTCAGT |
| BRCA2 | 80 | 8 | GGCCTGGGAACTCTCCTGTTCTTTGA | AGAGTGAGTAGTAGTGGAGT | AGAGTGAGTAGTAGTGGAGT | AGAGTGAGTAGTAGTGGAGTTTTAGAGTGAGTAGTAGTGGAGTTTTGGCCTGGGAACTCTCCTGTTCTTTGA |
| CENPF | 1 | 9 | CTTTTCAGGTTTGTACCCTCGGTTTT | TGTGGAGGGATTGAAGGATA | TGTGGAGGGATTGAAGGATA | TGTGGAGGGATTGAAGGATATTTTGTGGAGGGATTGAAGGATATTTCTTTTCAGGTTTGTACCCTCGGTTTT |
| CENPF | 2 | 9 | CCTGGAAATTCACTTGTGACTCCTTG | TGTGATGGAAGTTAGAGGGT | TGTGATGGAAGTTAGAGGGT | TGTGATGGAAGTTAGAGGGTTTTTGTGATGGAAGTTAGAGGGTTTTCCTGGAAATTCACTTGTGACTCCTTG |
| CENPF | 3 | 9 | GCATCTCTTTTCCCTATTTGCAAAGT | TGTGGAGGGATTGAAGGATA | TGTGGAGGGATTGAAGGATA | TGTGGAGGGATTGAAGGATATTTTGTGGAGGGATTGAAGGATATTTGCATCTCTTTTCCCTATTTGCAAAGT |
| CENPF | 4 | 9 | CTGCTAGAATTGTCAAAGAAACTGCTA | TGTGATGGAAGTTAGAGGGT | TGTGATGGAAGTTAGAGGGT | TGTGATGGAAGTTAGAGGGTTTTTGTGATGGAAGTTAGAGGGTTTTCTGCTAGAATTGTCAAAGAAACTGCTA |
| CENPF | 5 | 9 | GAAACTTATTCACTTGGCCTTTCATTTC | TGTGGAGGGATTGAAGGATA | TGTGGAGGGATTGAAGGATA | TGTGGAGGGATTGAAGGATATTTTGTGGAGGGATTGAAGGATATTTGAAACTTATTCACTTGGCCTTTCATTTC |
| CENPF | 6 | 9 | TCATCCCTACATTTGTTCAAAACTTTCTC | TGTGATGGAAGTTAGAGGGT | TGTGATGGAAGTTAGAGGGT | TGTGATGGAAGTTAGAGGGTTTTTGTGATGGAAGTTAGAGGGTTTTTCATCCCTACATTTGTTCAAAACTTTCTC |
| CENPF | 7 | 9 | TGTTTGGAAAGAACGCTGTTGACGGG | TGTGGAGGGATTGAAGGATA | TGTGGAGGGATTGAAGGATA | TGTGGAGGGATTGAAGGATATTTTGTGGAGGGATTGAAGGATATTTTGTTTGGAAAGAACGCTGTTGACGGG |
| CENPF | 8 | 9 | GCCTTCATCTGGATGCACTCCTGGTC | TGTGATGGAAGTTAGAGGGT | TGTGATGGAAGTTAGAGGGT | TGTGATGGAAGTTAGAGGGTTTTTGTGATGGAAGTTAGAGGGTTTTGCCTTCATCTGGATGCACTCCTGGTC |
| CENPF | 9 | 9 | GATCTGACTCGCCTGGAACGCCTGTT | TGTGGAGGGATTGAAGGATA | TGTGGAGGGATTGAAGGATA | TGTGGAGGGATTGAAGGATATTTTGTGGAGGGATTGAAGGATATTTGATCTGACTCGCCTGGAACGCCTGTT |
| CENPF | 10 | 9 | TGTTCTTGAGTTCTGCCTCCAGGTGG | TGTGATGGAAGTTAGAGGGT | TGTGATGGAAGTTAGAGGGT | TGTGATGGAAGTTAGAGGGTTTTTGTGATGGAAGTTAGAGGGTTTTTGTTCTTGAGTTCTGCCTCCAGGTGG |
| CENPF | 11 | 9 | CCAGATCAGCCACAGCAAGCTTCAGT | TGTGGAGGGATTGAAGGATA | TGTGGAGGGATTGAAGGATA | TGTGGAGGGATTGAAGGATATTTTGTGGAGGGATTGAAGGATATTTCCAGATCAGCCACAGCAAGCTTCAGT |
| CENPF | 12 | 9 | ACTCTTTCTCTGTCTTGCTTAACTTATCA | TGTGATGGAAGTTAGAGGGT | TGTGATGGAAGTTAGAGGGT | TGTGATGGAAGTTAGAGGGTTTTTGTGATGGAAGTTAGAGGGTTTTACTCTTTCTCTGTCTTGCTTAACTTATCA |
| CENPF | 13 | 9 | CGTTCTTACTCTCTCGTTGTATTCATGAC | TGTGGAGGGATTGAAGGATA | TGTGGAGGGATTGAAGGATA | TGTGGAGGGATTGAAGGATATTTTGTGGAGGGATTGAAGGATATTTCGTTCTTACTCTCTCGTTGTATTCATGAC |
| CENPF | 14 | 9 | CTGGGTCTCTACCTCCACTGACTTAC | TGTGATGGAAGTTAGAGGGT | TGTGATGGAAGTTAGAGGGT | TGTGATGGAAGTTAGAGGGTTTTTGTGATGGAAGTTAGAGGGTTTTCTGGGTCTCTACCTCCACTGACTTAC |
| CENPF | 15 | 9 | CATATTCGGCATGCAAGTCTTGGTAA | TGTGGAGGGATTGAAGGATA | TGTGGAGGGATTGAAGGATA | TGTGGAGGGATTGAAGGATATTTTGTGGAGGGATTGAAGGATATTTCATATTCGGCATGCAAGTCTTGGTAA |
| CENPF | 16 | 9 | GCTGATCAAAAGCCAAAAGACTTCTC | TGTGATGGAAGTTAGAGGGT | TGTGATGGAAGTTAGAGGGT | TGTGATGGAAGTTAGAGGGTTTTTGTGATGGAAGTTAGAGGGTTTTGCTGATCAAAAGCCAAAAGACTTCTC |
| CENPF | 17 | 9 | TCAACTCTATTTTGTAGGATGGCAGAA | TGTGGAGGGATTGAAGGATA | TGTGGAGGGATTGAAGGATA | TGTGGAGGGATTGAAGGATATTTTGTGGAGGGATTGAAGGATATTTTCAACTCTATTTTGTAGGATGGCAGAA |
| CENPF | 18 | 9 | TTTGCAGGTCTGAGTTCATCTGTTTT | TGTGATGGAAGTTAGAGGGT | TGTGATGGAAGTTAGAGGGT | TGTGATGGAAGTTAGAGGGTTTTTGTGATGGAAGTTAGAGGGTTTTTTTGCAGGTCTGAGTTCATCTGTTTT |
| CENPF | 19 | 9 | CTGTTCTGCTTTCATGAGATTTTCTTCTA | TGTGGAGGGATTGAAGGATA | TGTGGAGGGATTGAAGGATA | TGTGGAGGGATTGAAGGATATTTTGTGGAGGGATTGAAGGATATTTCTGTTCTGCTTTCATGAGATTTTCTTCTA |
| CENPF | 20 | 9 | AGTCTTCAAGTAGATGGTTGCTCTTTT | TGTGATGGAAGTTAGAGGGT | TGTGATGGAAGTTAGAGGGT | TGTGATGGAAGTTAGAGGGTTTTTGTGATGGAAGTTAGAGGGTTTTAGTCTTCAAGTAGATGGTTGCTCTTTT |
| CENPF | 21 | 9 | GCATTAATTTCCTTAAGAGTCCCATTCTC | TGTGGAGGGATTGAAGGATA | TGTGGAGGGATTGAAGGATA | TGTGGAGGGATTGAAGGATATTTTGTGGAGGGATTGAAGGATATTTGCATTAATTTCCTTAAGAGTCCCATTCTC |
| CENPF | 22 | 9 | CTCTCACTTTTCTGGATTAAGTTCATCTTC | TGTGATGGAAGTTAGAGGGT | TGTGATGGAAGTTAGAGGGT | TGTGATGGAAGTTAGAGGGTTTTTGTGATGGAAGTTAGAGGGTTTTCTCTCACTTTTCTGGATTAAGTTCATCTTC |
| CENPF | 23 | 9 | ACTAGTGCATTCATTTAGCAAGCATTC | TGTGGAGGGATTGAAGGATA | TGTGGAGGGATTGAAGGATA | TGTGGAGGGATTGAAGGATATTTTGTGGAGGGATTGAAGGATATTTACTAGTGCATTCATTTAGCAAGCATTC |
| CENPF | 24 | 9 | CTTCCTTTAGCTGTTCCAACTCATTTT | TGTGATGGAAGTTAGAGGGT | TGTGATGGAAGTTAGAGGGT | TGTGATGGAAGTTAGAGGGTTTTTGTGATGGAAGTTAGAGGGTTTTCTTCCTTTAGCTGTTCCAACTCATTTT |
| CENPF | 25 | 9 | CATCAGCTGTTCATTCTCTTGTAATAAGT | TGTGGAGGGATTGAAGGATA | TGTGGAGGGATTGAAGGATA | TGTGGAGGGATTGAAGGATATTTTGTGGAGGGATTGAAGGATATTTCATCAGCTGTTCATTCTCTTGTAATAAGT |
| CENPF | 26 | 9 | AGAGTTCCTAATTGGTTCTGATTCTAGAT | TGTGATGGAAGTTAGAGGGT | TGTGATGGAAGTTAGAGGGT | TGTGATGGAAGTTAGAGGGTTTTTGTGATGGAAGTTAGAGGGTTTTAGAGTTCCTAATTGGTTCTGATTCTAGAT |
| CENPF | 27 | 9 | CCAACTGCGCATTATAACTATCTAGAGAA | TGTGGAGGGATTGAAGGATA | TGTGGAGGGATTGAAGGATA | TGTGGAGGGATTGAAGGATATTTTGTGGAGGGATTGAAGGATATTTCCAACTGCGCATTATAACTATCTAGAGAA |
| CENPF | 28 | 9 | TCATGAGGCCCTGAAATATACTTTTCT | TGTGATGGAAGTTAGAGGGT | TGTGATGGAAGTTAGAGGGT | TGTGATGGAAGTTAGAGGGTTTTTGTGATGGAAGTTAGAGGGTTTTTCATGAGGCCCTGAAATATACTTTTCT |
| CENPF | 29 | 9 | CATTGTTGTTTGCAGAGAGCACTGAA | TGTGGAGGGATTGAAGGATA | TGTGGAGGGATTGAAGGATA | TGTGGAGGGATTGAAGGATATTTTGTGGAGGGATTGAAGGATATTTCATTGTTGTTTGCAGAGAGCACTGAA |
| CENPF | 30 | 9 | GAGTTCATACTTTTCAGCCTGCAGTA | TGTGATGGAAGTTAGAGGGT | TGTGATGGAAGTTAGAGGGT | TGTGATGGAAGTTAGAGGGTTTTTGTGATGGAAGTTAGAGGGTTTTGAGTTCATACTTTTCAGCCTGCAGTA |
| CENPF | 31 | 9 | CTCCTGGTATGTCTTCCACTAACTCA | TGTGGAGGGATTGAAGGATA | TGTGGAGGGATTGAAGGATA | TGTGGAGGGATTGAAGGATATTTTGTGGAGGGATTGAAGGATATTTCTCCTGGTATGTCTTCCACTAACTCA |
| CENPF | 32 | 9 | CTCGTCCAATGGAGCCAAAGACACAG | TGTGATGGAAGTTAGAGGGT | TGTGATGGAAGTTAGAGGGT | TGTGATGGAAGTTAGAGGGTTTTTGTGATGGAAGTTAGAGGGTTTTCTCGTCCAATGGAGCCAAAGACACAG |
| CENPF | 33 | 9 | CCAAATTTTCGGCCTTTAATGAGTCAA | TGTGGAGGGATTGAAGGATA | TGTGGAGGGATTGAAGGATA | TGTGGAGGGATTGAAGGATATTTTGTGGAGGGATTGAAGGATATTTCCAAATTTTCGGCCTTTAATGAGTCAA |
| CENPF | 34 | 9 | CTTGAAAGTTTCTCAGATTCGTTGACA | TGTGATGGAAGTTAGAGGGT | TGTGATGGAAGTTAGAGGGT | TGTGATGGAAGTTAGAGGGTTTTTGTGATGGAAGTTAGAGGGTTTTCTTGAAAGTTTCTCAGATTCGTTGACA |
| CENPF | 35 | 9 | TGCTAAGACTAGAGCTGTCAGGCACA | TGTGGAGGGATTGAAGGATA | TGTGGAGGGATTGAAGGATA | TGTGGAGGGATTGAAGGATATTTTGTGGAGGGATTGAAGGATATTTTGCTAAGACTAGAGCTGTCAGGCACA |
| CENPF | 36 | 9 | GGGTTTCTTTCCTGGTCAGATTCTCC | TGTGATGGAAGTTAGAGGGT | TGTGATGGAAGTTAGAGGGT | TGTGATGGAAGTTAGAGGGTTTTTGTGATGGAAGTTAGAGGGTTTTGGGTTTCTTTCCTGGTCAGATTCTCC |
| CENPF | 37 | 9 | CAAGCTCTTGCCTTTCAGAACTTAATA | TGTGGAGGGATTGAAGGATA | TGTGGAGGGATTGAAGGATA | TGTGGAGGGATTGAAGGATATTTTGTGGAGGGATTGAAGGATATTTCAAGCTCTTGCCTTTCAGAACTTAATA |
| CENPF | 38 | 9 | TTCTGACAAATACTGCTTCCTAAGGC | TGTGATGGAAGTTAGAGGGT | TGTGATGGAAGTTAGAGGGT | TGTGATGGAAGTTAGAGGGTTTTTGTGATGGAAGTTAGAGGGTTTTTTCTGACAAATACTGCTTCCTAAGGC |
| CENPF | 39 | 9 | CAGTTGTTCCGTCTGTTTCTTTTCTG | TGTGGAGGGATTGAAGGATA | TGTGGAGGGATTGAAGGATA | TGTGGAGGGATTGAAGGATATTTTGTGGAGGGATTGAAGGATATTTCAGTTGTTCCGTCTGTTTCTTTTCTG |
| CENPF | 40 | 9 | ACCGAGAACTTAAGTCCAGACCTTGT | TGTGATGGAAGTTAGAGGGT | TGTGATGGAAGTTAGAGGGT | TGTGATGGAAGTTAGAGGGTTTTTGTGATGGAAGTTAGAGGGTTTTACCGAGAACTTAAGTCCAGACCTTGT |
| CENPF | 41 | 9 | TGGTGTTCTTTCTGTAGTTTCTGAAGTAT | TGTGGAGGGATTGAAGGATA | TGTGGAGGGATTGAAGGATA | TGTGGAGGGATTGAAGGATATTTTGTGGAGGGATTGAAGGATATTTTGGTGTTCTTTCTGTAGTTTCTGAAGTAT |
| CENPF | 42 | 9 | CTGGAGGCTCATAATTGGTATCTGGG | TGTGATGGAAGTTAGAGGGT | TGTGATGGAAGTTAGAGGGT | TGTGATGGAAGTTAGAGGGTTTTTGTGATGGAAGTTAGAGGGTTTTCTGGAGGCTCATAATTGGTATCTGGG |
| CENPF | 43 | 9 | GCATTAGGACCAGAAAATGACAATTCAG | TGTGGAGGGATTGAAGGATA | TGTGGAGGGATTGAAGGATA | TGTGGAGGGATTGAAGGATATTTTGTGGAGGGATTGAAGGATATTTGCATTAGGACCAGAAAATGACAATTCAG |
| CENPF | 44 | 9 | TCTCATTTGATGTCTCTTTTACCCGC | TGTGATGGAAGTTAGAGGGT | TGTGATGGAAGTTAGAGGGT | TGTGATGGAAGTTAGAGGGTTTTTGTGATGGAAGTTAGAGGGTTTTTCTCATTTGATGTCTCTTTTACCCGC |
| CENPF | 45 | 9 | TGAGGCCTTCAGAAGTTTCTACTCTC | TGTGGAGGGATTGAAGGATA | TGTGGAGGGATTGAAGGATA | TGTGGAGGGATTGAAGGATATTTTGTGGAGGGATTGAAGGATATTTTGAGGCCTTCAGAAGTTTCTACTCTC |
| CENPF | 46 | 9 | ACGTGATGATTTATCTGCATGCATTTC | TGTGATGGAAGTTAGAGGGT | TGTGATGGAAGTTAGAGGGT | TGTGATGGAAGTTAGAGGGTTTTTGTGATGGAAGTTAGAGGGTTTTACGTGATGATTTATCTGCATGCATTTC |
| CENPF | 47 | 9 | CTTGTGACCACTGAGAGTTCTTCTTC | TGTGGAGGGATTGAAGGATA | TGTGGAGGGATTGAAGGATA | TGTGGAGGGATTGAAGGATATTTTGTGGAGGGATTGAAGGATATTTCTTGTGACCACTGAGAGTTCTTCTTC |
| CENPF | 48 | 9 | CTAATTCTCCACGAAGCTGGTTTCTC | TGTGATGGAAGTTAGAGGGT | TGTGATGGAAGTTAGAGGGT | TGTGATGGAAGTTAGAGGGTTTTTGTGATGGAAGTTAGAGGGTTTTCTAATTCTCCACGAAGCTGGTTTCTC |
| CENPF | 49 | 9 | GTTTTGTCTTTTAACAGCTCACTCACAT | TGTGGAGGGATTGAAGGATA | TGTGGAGGGATTGAAGGATA | TGTGGAGGGATTGAAGGATATTTTGTGGAGGGATTGAAGGATATTTGTTTTGTCTTTTAACAGCTCACTCACAT |
| CENPF | 50 | 9 | GATTCCTTGACAAGCAATTCTTTCTCTTT | TGTGATGGAAGTTAGAGGGT | TGTGATGGAAGTTAGAGGGT | TGTGATGGAAGTTAGAGGGTTTTTGTGATGGAAGTTAGAGGGTTTTGATTCCTTGACAAGCAATTCTTTCTCTTT |
| CENPF | 51 | 9 | TTTCTCGATGCCTCTTCTCAGCTGAT | TGTGGAGGGATTGAAGGATA | TGTGGAGGGATTGAAGGATA | TGTGGAGGGATTGAAGGATATTTTGTGGAGGGATTGAAGGATATTTTTTCTCGATGCCTCTTCTCAGCTGAT |
| CENPF | 52 | 9 | TTTTCATCGGCCTCAATGCGAACTCT | TGTGATGGAAGTTAGAGGGT | TGTGATGGAAGTTAGAGGGT | TGTGATGGAAGTTAGAGGGTTTTTGTGATGGAAGTTAGAGGGTTTTTTTTCATCGGCCTCAATGCGAACTCT |
| CENPF | 53 | 9 | ATTCCCTTTCAAGGTTCTCAACTTTATC | TGTGGAGGGATTGAAGGATA | TGTGGAGGGATTGAAGGATA | TGTGGAGGGATTGAAGGATATTTTGTGGAGGGATTGAAGGATATTTATTCCCTTTCAAGGTTCTCAACTTTATC |
| CENPF | 54 | 9 | TAGCTCCTGGTTTTCTTCTGACATCT | TGTGATGGAAGTTAGAGGGT | TGTGATGGAAGTTAGAGGGT | TGTGATGGAAGTTAGAGGGTTTTTGTGATGGAAGTTAGAGGGTTTTTAGCTCCTGGTTTTCTTCTGACATCT |
| CENPF | 55 | 9 | CTGGCCATCTCTTCTATTTGTGTTTT | TGTGGAGGGATTGAAGGATA | TGTGGAGGGATTGAAGGATA | TGTGGAGGGATTGAAGGATATTTTGTGGAGGGATTGAAGGATATTTCTGGCCATCTCTTCTATTTGTGTTTT |
| CENPF | 56 | 9 | CTTGCTCCTTTTCTTCTAACAGACTTTT | TGTGATGGAAGTTAGAGGGT | TGTGATGGAAGTTAGAGGGT | TGTGATGGAAGTTAGAGGGTTTTTGTGATGGAAGTTAGAGGGTTTTCTTGCTCCTTTTCTTCTAACAGACTTTT |
| CENPF | 57 | 9 | TGTTCTGTGGCCTTCATAATTTCTTG | TGTGGAGGGATTGAAGGATA | TGTGGAGGGATTGAAGGATA | TGTGGAGGGATTGAAGGATATTTTGTGGAGGGATTGAAGGATATTTTGTTCTGTGGCCTTCATAATTTCTTG |
| CENPF | 58 | 9 | GCTCTTCCTCTATTGGTGGGTCTAGA | TGTGATGGAAGTTAGAGGGT | TGTGATGGAAGTTAGAGGGT | TGTGATGGAAGTTAGAGGGTTTTTGTGATGGAAGTTAGAGGGTTTTGCTCTTCCTCTATTGGTGGGTCTAGA |
| CENPF | 59 | 9 | CCTTCAGTTGTTGTAAGACACAGAGC | TGTGGAGGGATTGAAGGATA | TGTGGAGGGATTGAAGGATA | TGTGGAGGGATTGAAGGATATTTTGTGGAGGGATTGAAGGATATTTCCTTCAGTTGTTGTAAGACACAGAGC |
| CENPF | 60 | 9 | CCTTAAGTAAATCTGCATGATGCTCAC | TGTGATGGAAGTTAGAGGGT | TGTGATGGAAGTTAGAGGGT | TGTGATGGAAGTTAGAGGGTTTTTGTGATGGAAGTTAGAGGGTTTTCCTTAAGTAAATCTGCATGATGCTCAC |
| CENPF | 61 | 9 | GCATGCTCTTGGTTTGTCCTGGCTAT | TGTGGAGGGATTGAAGGATA | TGTGGAGGGATTGAAGGATA | TGTGGAGGGATTGAAGGATATTTTGTGGAGGGATTGAAGGATATTTGCATGCTCTTGGTTTGTCCTGGCTAT |
| CENPF | 62 | 9 | CTAATTCAGATATTCGCTCTTGCTCTTTT | TGTGATGGAAGTTAGAGGGT | TGTGATGGAAGTTAGAGGGT | TGTGATGGAAGTTAGAGGGTTTTTGTGATGGAAGTTAGAGGGTTTTCTAATTCAGATATTCGCTCTTGCTCTTTT |
| CENPF | 63 | 9 | CTCTCTCATTGAGCTCTTTTAATTGTGT | TGTGGAGGGATTGAAGGATA | TGTGGAGGGATTGAAGGATA | TGTGGAGGGATTGAAGGATATTTTGTGGAGGGATTGAAGGATATTTCTCTCTCATTGAGCTCTTTTAATTGTGT |
| CENPF | 64 | 9 | CTACTAAGATTCTGCTCTTTGGCCTT | TGTGATGGAAGTTAGAGGGT | TGTGATGGAAGTTAGAGGGT | TGTGATGGAAGTTAGAGGGTTTTTGTGATGGAAGTTAGAGGGTTTTCTACTAAGATTCTGCTCTTTGGCCTT |
| CENPF | 65 | 9 | GCTGCTCTTGGTCTTGAATTTGATTTT | TGTGGAGGGATTGAAGGATA | TGTGGAGGGATTGAAGGATA | TGTGGAGGGATTGAAGGATATTTTGTGGAGGGATTGAAGGATATTTGCTGCTCTTGGTCTTGAATTTGATTTT |
| CENPF | 66 | 9 | TTTGGATTGTAGCACTTGGATCTTCT | TGTGATGGAAGTTAGAGGGT | TGTGATGGAAGTTAGAGGGT | TGTGATGGAAGTTAGAGGGTTTTTGTGATGGAAGTTAGAGGGTTTTTTTGGATTGTAGCACTTGGATCTTCT |
| CENPF | 67 | 9 | CATTCTCTAGATTCTTGTAAGAACTCTGC | TGTGGAGGGATTGAAGGATA | TGTGGAGGGATTGAAGGATA | TGTGGAGGGATTGAAGGATATTTTGTGGAGGGATTGAAGGATATTTCATTCTCTAGATTCTTGTAAGAACTCTGC |
| CENPF | 68 | 9 | CAGTTTCCTTTGCAGTCATTTTGTTTAC | TGTGATGGAAGTTAGAGGGT | TGTGATGGAAGTTAGAGGGT | TGTGATGGAAGTTAGAGGGTTTTTGTGATGGAAGTTAGAGGGTTTTCAGTTTCCTTTGCAGTCATTTTGTTTAC |
| CENPF | 69 | 9 | TTTCTAGTGTGAGCTCCTTCAATTGA | TGTGGAGGGATTGAAGGATA | TGTGGAGGGATTGAAGGATA | TGTGGAGGGATTGAAGGATATTTTGTGGAGGGATTGAAGGATATTTTTTCTAGTGTGAGCTCCTTCAATTGA |
| CENPF | 70 | 9 | GCAATCTAGGCTCTTCTTCAATTCAC | TGTGATGGAAGTTAGAGGGT | TGTGATGGAAGTTAGAGGGT | TGTGATGGAAGTTAGAGGGTTTTTGTGATGGAAGTTAGAGGGTTTTGCAATCTAGGCTCTTCTTCAATTCAC |
| CENPF | 71 | 9 | GTTTGTTTGTGTCCAAAAGCAAAGCC | TGTGGAGGGATTGAAGGATA | TGTGGAGGGATTGAAGGATA | TGTGGAGGGATTGAAGGATATTTTGTGGAGGGATTGAAGGATATTTGTTTGTTTGTGTCCAAAAGCAAAGCC |
| CENPF | 72 | 9 | CTCGGTATGTCTGGATTTCTACTTCAT | TGTGATGGAAGTTAGAGGGT | TGTGATGGAAGTTAGAGGGT | TGTGATGGAAGTTAGAGGGTTTTTGTGATGGAAGTTAGAGGGTTTTCTCGGTATGTCTGGATTTCTACTTCAT |
| CENPF | 73 | 9 | GCTTGAAGTTGAGAGAGTTCTTTCTG | TGTGGAGGGATTGAAGGATA | TGTGGAGGGATTGAAGGATA | TGTGGAGGGATTGAAGGATATTTTGTGGAGGGATTGAAGGATATTTGCTTGAAGTTGAGAGAGTTCTTTCTG |
| CENPF | 74 | 9 | TGTTAATTCATCGACCTTGGTATCCA | TGTGATGGAAGTTAGAGGGT | TGTGATGGAAGTTAGAGGGT | TGTGATGGAAGTTAGAGGGTTTTTGTGATGGAAGTTAGAGGGTTTTTGTTAATTCATCGACCTTGGTATCCA |
| CENPF | 75 | 9 | TGTGTCTCTAACATCTCTTTAGCTTTCT | TGTGGAGGGATTGAAGGATA | TGTGGAGGGATTGAAGGATA | TGTGGAGGGATTGAAGGATATTTTGTGGAGGGATTGAAGGATATTTTGTGTCTCTAACATCTCTTTAGCTTTCT |
| CENPF | 76 | 9 | TAGCAAAGGAGACCCTCGGGAATCTT | TGTGATGGAAGTTAGAGGGT | TGTGATGGAAGTTAGAGGGT | TGTGATGGAAGTTAGAGGGTTTTTGTGATGGAAGTTAGAGGGTTTTTAGCAAAGGAGACCCTCGGGAATCTT |
| CENPF | 77 | 9 | TTTGCCTCTTGCCTGAAGCTTTATTT | TGTGGAGGGATTGAAGGATA | TGTGGAGGGATTGAAGGATA | TGTGGAGGGATTGAAGGATATTTTGTGGAGGGATTGAAGGATATTTTTTGCCTCTTGCCTGAAGCTTTATTT |
| CENPF | 78 | 9 | TCTACCATTCTCCCATATTCCACTGG | TGTGATGGAAGTTAGAGGGT | TGTGATGGAAGTTAGAGGGT | TGTGATGGAAGTTAGAGGGTTTTTGTGATGGAAGTTAGAGGGTTTTTCTACCATTCTCCCATATTCCACTGG |
| CENPF | 79 | 9 | GCTGGTCCGAGTTGCCATGGTTGTTC | TGTGGAGGGATTGAAGGATA | TGTGGAGGGATTGAAGGATA | TGTGGAGGGATTGAAGGATATTTTGTGGAGGGATTGAAGGATATTTGCTGGTCCGAGTTGCCATGGTTGTTC |
| CENPF | 80 | 9 | GGATAGCGCTAACTTCTGTGCAGCCA | TGTGATGGAAGTTAGAGGGT | TGTGATGGAAGTTAGAGGGT | TGTGATGGAAGTTAGAGGGTTTTTGTGATGGAAGTTAGAGGGTTTTGGATAGCGCTAACTTCTGTGCAGCCA |
| CKAP5 | 1 | 10 | CACTGCATTGGAATCAGTGACAAATTTT | TGTGGAGGGATTGAAGGATA | TGTGGAGGGATTGAAGGATA | TGTGGAGGGATTGAAGGATATTTTGTGGAGGGATTGAAGGATATTTCACTGCATTGGAATCAGTGACAAATTTT |
| CKAP5 | 2 | 10 | AGTGCAGCTTCTAATCCTTTCAATTG | GGGATGTATTGAAGGAGGAT | GGGATGTATTGAAGGAGGAT | GGGATGTATTGAAGGAGGATTTTGGGATGTATTGAAGGAGGATTTTAGTGCAGCTTCTAATCCTTTCAATTG |
| CKAP5 | 3 | 10 | GCTTTAGGTTGATTGAACACCTTACTTAC | TGTGGAGGGATTGAAGGATA | TGTGGAGGGATTGAAGGATA | TGTGGAGGGATTGAAGGATATTTTGTGGAGGGATTGAAGGATATTTGCTTTAGGTTGATTGAACACCTTACTTAC |
| CKAP5 | 4 | 10 | AGATCTCTATGCCCAGCTCCTTGGCT | GGGATGTATTGAAGGAGGAT | GGGATGTATTGAAGGAGGAT | GGGATGTATTGAAGGAGGATTTTGGGATGTATTGAAGGAGGATTTTAGATCTCTATGCCCAGCTCCTTGGCT |
| CKAP5 | 5 | 10 | CAAGCCTTTCAGGAGCTCTTCTTGAA | TGTGGAGGGATTGAAGGATA | TGTGGAGGGATTGAAGGATA | TGTGGAGGGATTGAAGGATATTTTGTGGAGGGATTGAAGGATATTTCAAGCCTTTCAGGAGCTCTTCTTGAA |
| CKAP5 | 6 | 10 | GGCCACTATGATCTTGGGATTCTTATT | GGGATGTATTGAAGGAGGAT | GGGATGTATTGAAGGAGGAT | GGGATGTATTGAAGGAGGATTTTGGGATGTATTGAAGGAGGATTTTGGCCACTATGATCTTGGGATTCTTATT |
| CKAP5 | 7 | 10 | GCTTCATCTCGAACAGCCTTCTCTCG | TGTGGAGGGATTGAAGGATA | TGTGGAGGGATTGAAGGATA | TGTGGAGGGATTGAAGGATATTTTGTGGAGGGATTGAAGGATATTTGCTTCATCTCGAACAGCCTTCTCTCG |
| CKAP5 | 8 | 10 | CCATCTGTAAATCTCCACAGCAATTAGT | GGGATGTATTGAAGGAGGAT | GGGATGTATTGAAGGAGGAT | GGGATGTATTGAAGGAGGATTTTGGGATGTATTGAAGGAGGATTTTCCATCTGTAAATCTCCACAGCAATTAGT |
| CKAP5 | 9 | 10 | TTGGGAACGAAGAAATCGAGTAGGTC | TGTGGAGGGATTGAAGGATA | TGTGGAGGGATTGAAGGATA | TGTGGAGGGATTGAAGGATATTTTGTGGAGGGATTGAAGGATATTTTTGGGAACGAAGAAATCGAGTAGGTC |
| CKAP5 | 10 | 10 | TATTTGTGGCACCTCATCACCATCAT | GGGATGTATTGAAGGAGGAT | GGGATGTATTGAAGGAGGAT | GGGATGTATTGAAGGAGGATTTTGGGATGTATTGAAGGAGGATTTTTATTTGTGGCACCTCATCACCATCAT |
| CKAP5 | 11 | 10 | GTCATAAAAGTCTTTGGGAAGTTTGGAAA | TGTGGAGGGATTGAAGGATA | TGTGGAGGGATTGAAGGATA | TGTGGAGGGATTGAAGGATATTTTGTGGAGGGATTGAAGGATATTTGTCATAAAAGTCTTTGGGAAGTTTGGAAA |
| CKAP5 | 12 | 10 | GGGCCTCTTTTCTCTCTTGCCATTTT | GGGATGTATTGAAGGAGGAT | GGGATGTATTGAAGGAGGAT | GGGATGTATTGAAGGAGGATTTTGGGATGTATTGAAGGAGGATTTTGGGCCTCTTTTCTCTCTTGCCATTTT |
| CKAP5 | 13 | 10 | TGCCAAAGCCACCAACATGACATTGG | TGTGGAGGGATTGAAGGATA | TGTGGAGGGATTGAAGGATA | TGTGGAGGGATTGAAGGATATTTTGTGGAGGGATTGAAGGATATTTTGCCAAAGCCACCAACATGACATTGG |
| CKAP5 | 14 | 10 | CCAACAGCCAGGCCAGTAAGACATTT | GGGATGTATTGAAGGAGGAT | GGGATGTATTGAAGGAGGAT | GGGATGTATTGAAGGAGGATTTTGGGATGTATTGAAGGAGGATTTTCCAACAGCCAGGCCAGTAAGACATTT |
| CKAP5 | 15 | 10 | TTGTACCACTTGAGGTTTCTTCTCTT | TGTGGAGGGATTGAAGGATA | TGTGGAGGGATTGAAGGATA | TGTGGAGGGATTGAAGGATATTTTGTGGAGGGATTGAAGGATATTTTTGTACCACTTGAGGTTTCTTCTCTT |
| CKAP5 | 16 | 10 | AAGATTGCATCAATTGCCTCCTGCAG | GGGATGTATTGAAGGAGGAT | GGGATGTATTGAAGGAGGAT | GGGATGTATTGAAGGAGGATTTTGGGATGTATTGAAGGAGGATTTTAAGATTGCATCAATTGCCTCCTGCAG |
| CKAP5 | 17 | 10 | AGTGGCGGAAACTTCTTGCAATAAAA | TGTGGAGGGATTGAAGGATA | TGTGGAGGGATTGAAGGATA | TGTGGAGGGATTGAAGGATATTTTGTGGAGGGATTGAAGGATATTTAGTGGCGGAAACTTCTTGCAATAAAA |
| CKAP5 | 18 | 10 | CAAGCTCTTTGGCAGGGTAGAAGCAG | GGGATGTATTGAAGGAGGAT | GGGATGTATTGAAGGAGGAT | GGGATGTATTGAAGGAGGATTTTGGGATGTATTGAAGGAGGATTTTCAAGCTCTTTGGCAGGGTAGAAGCAG |
| CKAP5 | 19 | 10 | TGACTTCAGGAGCAGAATCATTGATG | TGTGGAGGGATTGAAGGATA | TGTGGAGGGATTGAAGGATA | TGTGGAGGGATTGAAGGATATTTTGTGGAGGGATTGAAGGATATTTTGACTTCAGGAGCAGAATCATTGATG |

| | | | | | | |
|---|---|---|---|---|---|---|
| CKAP5 | 20 | 10 | AGTACCCAATGCTTCAAATGCGGCAT | GGGATGTATTGAAGGAGGAT | GGGATGTATTGAAGGAGGAT | GGGATGTATTGAAGGAGGATTTTGGGATGTATTGAAGGAGGATTTTAGTACCCAATGCTTCAAATGCGGCAT |
| CKAP5 | 21 | 10 | GTTTGTCCACATCAGCTAGGAATGGG | TGTGGAGGGATTGAAGGATA | TGTGGAGGGATTGAAGGATA | TGTGGAGGGATTGAAGGATATTTTGTGGAGGGATTGAAGGATATTTGTTTGTCCACATCAGCTAGGAATGGG |
| CKAP5 | 22 | 10 | CAGGCAGAGGTTTGAATTCCTTCTTA | GGGATGTATTGAAGGAGGAT | GGGATGTATTGAAGGAGGAT | GGGATGTATTGAAGGAGGATTTTGGGATGTATTGAAGGAGGATTTTCAGGCAGAGGTTTGAATTCCTTCTTA |
| CKAP5 | 23 | 10 | AGAGGTCCTGGTTTGGGTGCAGAAAT | TGTGGAGGGATTGAAGGATA | TGTGGAGGGATTGAAGGATA | TGTGGAGGGATTGAAGGATATTTTGTGGAGGGATTGAAGGATATTTAGAGGTCCTGGTTTGGGTGCAGAAAT |
| CKAP5 | 24 | 10 | GGTTCCAGTATTCCCTGCGCCTCCTG | GGGATGTATTGAAGGAGGAT | GGGATGTATTGAAGGAGGAT | GGGATGTATTGAAGGAGGATTTTGGGATGTATTGAAGGAGGATTTTGGTTCCAGTATTCCCTGCGCCTCCTG |
| CKAP5 | 25 | 10 | ACAAGCCAGCCTTTCTTTCCAGTTAC | TGTGGAGGGATTGAAGGATA | TGTGGAGGGATTGAAGGATA | TGTGGAGGGATTGAAGGATATTTTGTGGAGGGATTGAAGGATATTTACAAGCCAGCCTTTCTTTCCAGTTAC |
| CKAP5 | 26 | 10 | CCTGGCATGGCATTTCAGTTCGGTCC | GGGATGTATTGAAGGAGGAT | GGGATGTATTGAAGGAGGAT | GGGATGTATTGAAGGAGGATTTTGGGATGTATTGAAGGAGGATTTTCCTGGCATGGCATTTCAGTTCGGTCC |
| CKAP5 | 27 | 10 | ACAACCTGAGCTGACGTTTTGGAAAA | TGTGGAGGGATTGAAGGATA | TGTGGAGGGATTGAAGGATA | TGTGGAGGGATTGAAGGATATTTTGTGGAGGGATTGAAGGATATTTACAACCTGAGCTGACGTTTTGGAAAA |
| CKAP5 | 28 | 10 | CTCCAATCTTGTCCACAAGGCCATCT | GGGATGTATTGAAGGAGGAT | GGGATGTATTGAAGGAGGAT | GGGATGTATTGAAGGAGGATTTTGGGATGTATTGAAGGAGGATTTTCTCCAATCTTGTCCACAAGGCCATCT |
| CKAP5 | 29 | 10 | TTTGGGATTCTTTTGTGAGAAAGCCA | TGTGGAGGGATTGAAGGATA | TGTGGAGGGATTGAAGGATA | TGTGGAGGGATTGAAGGATATTTTGTGGAGGGATTGAAGGATATTTTTTGGGATTCTTTTGTGAGAAAGCCA |
| CKAP5 | 30 | 10 | TACAGATACATCACGCCAAGCAGGGT | GGGATGTATTGAAGGAGGAT | GGGATGTATTGAAGGAGGAT | GGGATGTATTGAAGGAGGATTTTGGGATGTATTGAAGGAGGATTTTTACAGATACATCACGCCAAGCAGGGT |
| CKAP5 | 31 | 10 | TCATCTGGTTCATCTCCATCTTCTCC | TGTGGAGGGATTGAAGGATA | TGTGGAGGGATTGAAGGATA | TGTGGAGGGATTGAAGGATATTTTGTGGAGGGATTGAAGGATATTTTCATCTGGTTCATCTCCATCTTCTCC |
| CKAP5 | 32 | 10 | GCAAAAGATCAACGACATCATTGCTC | GGGATGTATTGAAGGAGGAT | GGGATGTATTGAAGGAGGAT | GGGATGTATTGAAGGAGGATTTTGGGATGTATTGAAGGAGGATTTTGCAAAAGATCAACGACATCATTGCTC |
| CKAP5 | 33 | 10 | GGAAGTTCACCTATATTCGGTTGGATAAAT | TGTGGAGGGATTGAAGGATA | TGTGGAGGGATTGAAGGATA | TGTGGAGGGATTGAAGGATATTTTGTGGAGGGATTGAAGGATATTTGGAAGTTCACCTATATTCGGTTGGATAAAT |
| CKAP5 | 34 | 10 | AATCATTGAGTCGACCCTTCAAGGCA | GGGATGTATTGAAGGAGGAT | GGGATGTATTGAAGGAGGAT | GGGATGTATTGAAGGAGGATTTTGGGATGTATTGAAGGAGGATTTTAATCATTGAGTCGACCCTTCAAGGCA |
| CKAP5 | 35 | 10 | CCAAGCATTCACAGTCGCTAGGGCAG | TGTGGAGGGATTGAAGGATA | TGTGGAGGGATTGAAGGATA | TGTGGAGGGATTGAAGGATATTTTGTGGAGGGATTGAAGGATATTTCCAAGCATTCACAGTCGCTAGGGCAG |
| CKAP5 | 36 | 10 | AGCCATTCCTTCATGCCAGTCTGTTC | GGGATGTATTGAAGGAGGAT | GGGATGTATTGAAGGAGGAT | GGGATGTATTGAAGGAGGATTTTGGGATGTATTGAAGGAGGATTTTAGCCATTCCTTCATGCCAGTCTGTTC |
| CKAP5 | 37 | 10 | GGTAGTTTCTCAGCCAGCCAGCCCAG | TGTGGAGGGATTGAAGGATA | TGTGGAGGGATTGAAGGATA | TGTGGAGGGATTGAAGGATATTTTGTGGAGGGATTGAAGGATATTTGGTAGTTTCTCAGCCAGCCAGCCCAG |
| CKAP5 | 38 | 10 | GAACACAAAGGATAAGGTCTGTAGGG | GGGATGTATTGAAGGAGGAT | GGGATGTATTGAAGGAGGAT | GGGATGTATTGAAGGAGGATTTTGGGATGTATTGAAGGAGGATTTTGAACACAAAGGATAAGGTCTGTAGGG |
| CKAP5 | 39 | 10 | TCTTGGGCCTTCTTTCGCACATCTCC | TGTGGAGGGATTGAAGGATA | TGTGGAGGGATTGAAGGATA | TGTGGAGGGATTGAAGGATATTTTGTGGAGGGATTGAAGGATATTTTCTTGGGCCTTCTTTCGCACATCTCC |
| CKAP5 | 40 | 10 | CCTAAATGCATCATGAAGAATGGCAAG | GGGATGTATTGAAGGAGGAT | GGGATGTATTGAAGGAGGAT | GGGATGTATTGAAGGAGGATTTTGGGATGTATTGAAGGAGGATTTTCCTAAATGCATCATGAAGAATGGCAAG |
| CKAP5 | 41 | 10 | GCATGTTAACTTTGGCTTTCTCTAGC | TGTGGAGGGATTGAAGGATA | TGTGGAGGGATTGAAGGATA | TGTGGAGGGATTGAAGGATATTTTGTGGAGGGATTGAAGGATATTTGCATGTTAACTTTGGCTTTCTCTAGC |
| CKAP5 | 42 | 10 | TGCTTTAGTGGGTGGAGCAGGCTTGG | GGGATGTATTGAAGGAGGAT | GGGATGTATTGAAGGAGGAT | GGGATGTATTGAAGGAGGATTTTGGGATGTATTGAAGGAGGATTTTTGCTTTAGTGGGTGGAGCAGGCTTGG |
| CKAP5 | 43 | 10 | TGCTCTTTTCCATTTGGAACAACAAT | TGTGGAGGGATTGAAGGATA | TGTGGAGGGATTGAAGGATA | TGTGGAGGGATTGAAGGATATTTTGTGGAGGGATTGAAGGATATTTTGCTCTTTTCCATTTGGAACAACAAT |
| CKAP5 | 44 | 10 | GTTGCTCAATGTATTCATCCCGTGGG | GGGATGTATTGAAGGAGGAT | GGGATGTATTGAAGGAGGAT | GGGATGTATTGAAGGAGGATTTTGGGATGTATTGAAGGAGGATTTTGTTGCTCAATGTATTCATCCCGTGGG |
| CKAP5 | 45 | 10 | TGAGTGAAACATCTCATCTTGTAACCA | TGTGGAGGGATTGAAGGATA | TGTGGAGGGATTGAAGGATA | TGTGGAGGGATTGAAGGATATTTTGTGGAGGGATTGAAGGATATTTTGAGTGAAACATCTCATCTTGTAACCA |
| CKAP5 | 46 | 10 | GCAAGGGCTTTGTTATGATGCTGAAA | GGGATGTATTGAAGGAGGAT | GGGATGTATTGAAGGAGGAT | GGGATGTATTGAAGGAGGATTTTGGGATGTATTGAAGGAGGATTTTGCAAGGGCTTTGTTATGATGCTGAAA |
| CKAP5 | 47 | 10 | TCAGGACGCTTGTATTGGTGTCAAAA | TGTGGAGGGATTGAAGGATA | TGTGGAGGGATTGAAGGATA | TGTGGAGGGATTGAAGGATATTTTGTGGAGGGATTGAAGGATATTTTCAGGACGCTTGTATTGGTGTCAAAA |
| CKAP5 | 48 | 10 | CACTTAGCAAGGTGAAGAGCAATTTT | GGGATGTATTGAAGGAGGAT | GGGATGTATTGAAGGAGGAT | GGGATGTATTGAAGGAGGATTTTGGGATGTATTGAAGGAGGATTTTCACTTAGCAAGGTGAAGAGCAATTTT |
| CKAP5 | 49 | 10 | CGAACATCTTTACGAATGACATCCTTT | TGTGGAGGGATTGAAGGATA | TGTGGAGGGATTGAAGGATA | TGTGGAGGGATTGAAGGATATTTTGTGGAGGGATTGAAGGATATTTCGAACATCTTTACGAATGACATCCTTT |
| CKAP5 | 50 | 10 | GATAAAGGGAAACATCTTGCTAGCTG | GGGATGTATTGAAGGAGGAT | GGGATGTATTGAAGGAGGAT | GGGATGTATTGAAGGAGGATTTTGGGATGTATTGAAGGAGGATTTTGATAAAGGGAAACATCTTGCTAGCTG |
| CKAP5 | 51 | 10 | TGCGTACAGCATTGTCACGGTCTCCT | TGTGGAGGGATTGAAGGATA | TGTGGAGGGATTGAAGGATA | TGTGGAGGGATTGAAGGATATTTTGTGGAGGGATTGAAGGATATTTTGCGTACAGCATTGTCACGGTCTCCT |
| CKAP5 | 52 | 10 | TCCAATCAGTTTGAACACCTGATCCC | GGGATGTATTGAAGGAGGAT | GGGATGTATTGAAGGAGGAT | GGGATGTATTGAAGGAGGATTTTGGGATGTATTGAAGGAGGATTTTTCCAATCAGTTTGAACACCTGATCCC |
| CKAP5 | 53 | 10 | CTCTTTGCTGACCGCTTAATCCTCTC | TGTGGAGGGATTGAAGGATA | TGTGGAGGGATTGAAGGATA | TGTGGAGGGATTGAAGGATATTTTGTGGAGGGATTGAAGGATATTTCTCTTTGCTGACCGCTTAATCCTCTC |
| CKAP5 | 54 | 10 | TATGTTCTGTGCACGCTGAGGTTTCT | GGGATGTATTGAAGGAGGAT | GGGATGTATTGAAGGAGGAT | GGGATGTATTGAAGGAGGATTTTGGGATGTATTGAAGGAGGATTTTTATGTTCTGTGCACGCTGAGGTTTCT |
| CKAP5 | 55 | 10 | CCTCAGGATGCCCACTCATGCTTCGG | TGTGGAGGGATTGAAGGATA | TGTGGAGGGATTGAAGGATA | TGTGGAGGGATTGAAGGATATTTTGTGGAGGGATTGAAGGATATTTCCTCAGGATGCCCACTCATGCTTCGG |
| CKAP5 | 56 | 10 | CTGGAATTCTCGGCGGACCATCTGGG | GGGATGTATTGAAGGAGGAT | GGGATGTATTGAAGGAGGAT | GGGATGTATTGAAGGAGGATTTTGGGATGTATTGAAGGAGGATTTTCTGGAATTCTCGGCGGACCATCTGGG |
| CKAP5 | 57 | 10 | GTTCAGGAATAAGGACTGGCTCAAAA | TGTGGAGGGATTGAAGGATA | TGTGGAGGGATTGAAGGATA | TGTGGAGGGATTGAAGGATATTTTGTGGAGGGATTGAAGGATATTTGTTCAGGAATAAGGACTGGCTCAAAA |
| CKAP5 | 58 | 10 | GAAGTGTGGAGAAACAGCCCGGATCT | GGGATGTATTGAAGGAGGAT | GGGATGTATTGAAGGAGGAT | GGGATGTATTGAAGGAGGATTTTGGGATGTATTGAAGGAGGATTTTGAAGTGTGGAGAAACAGCCCGGATCT |
| CKAP5 | 59 | 10 | CTGGCTACTTGGGAGATAATGAAATTG | TGTGGAGGGATTGAAGGATA | TGTGGAGGGATTGAAGGATA | TGTGGAGGGATTGAAGGATATTTTGTGGAGGGATTGAAGGATATTTCTGGCTACTTGGGAGATAATGAAATTG |
| CKAP5 | 60 | 10 | GAGCTTGGATACTTGTGTTGATGTCA | GGGATGTATTGAAGGAGGAT | GGGATGTATTGAAGGAGGAT | GGGATGTATTGAAGGAGGATTTTGGGATGTATTGAAGGAGGATTTTGAGCTTGGATACTTGTGTTGATGTCA |
| CKAP5 | 61 | 10 | TGGCCGGACATGGCTTCAGCTTTGTC | TGTGGAGGGATTGAAGGATA | TGTGGAGGGATTGAAGGATA | TGTGGAGGGATTGAAGGATATTTTGTGGAGGGATTGAAGGATATTTTGGCCGGACATGGCTTCAGCTTTGTC |
| CKAP5 | 62 | 10 | GCATAAAAGTGGCTATCAGAAACTGATCA | GGGATGTATTGAAGGAGGAT | GGGATGTATTGAAGGAGGAT | GGGATGTATTGAAGGAGGATTTTGGGATGTATTGAAGGAGGATTTTGCATAAAAGTGGCTATCAGAAACTGATCA |
| CKAP5 | 63 | 10 | GATCTCGTCCTTCTCCAATTTCTCAT | TGTGGAGGGATTGAAGGATA | TGTGGAGGGATTGAAGGATA | TGTGGAGGGATTGAAGGATATTTTGTGGAGGGATTGAAGGATATTTGATCTCGTCCTTCTCCAATTTCTCAT |
| CKAP5 | 64 | 10 | GTTGCCAATGATACAGCTATACAACTT | GGGATGTATTGAAGGAGGAT | GGGATGTATTGAAGGAGGAT | GGGATGTATTGAAGGAGGATTTTGGGATGTATTGAAGGAGGATTTTGTTGCCAATGATACAGCTATACAACTT |
| CKAP5 | 65 | 10 | TCAAGATCTTCAATCCGAGAATCCAG | TGTGGAGGGATTGAAGGATA | TGTGGAGGGATTGAAGGATA | TGTGGAGGGATTGAAGGATATTTTGTGGAGGGATTGAAGGATATTTTCAAGATCTTCAATCCGAGAATCCAG |
| CKAP5 | 66 | 10 | TCACAGAGCGGATGACCTGTTGTCCT | GGGATGTATTGAAGGAGGAT | GGGATGTATTGAAGGAGGAT | GGGATGTATTGAAGGAGGATTTTGGGATGTATTGAAGGAGGATTTTTCACAGAGCGGATGACCTGTTGTCCT |
| CKAP5 | 67 | 10 | CTAGCAGGCTGTCTTGGAGCAAAACA | TGTGGAGGGATTGAAGGATA | TGTGGAGGGATTGAAGGATA | TGTGGAGGGATTGAAGGATATTTTGTGGAGGGATTGAAGGATATTTCTAGCAGGCTGTCTTGGAGCAAAACA |
| CKAP5 | 68 | 10 | CTCTGAGAATTTGGGAGAACTGGCTG | GGGATGTATTGAAGGAGGAT | GGGATGTATTGAAGGAGGAT | GGGATGTATTGAAGGAGGATTTTGGGATGTATTGAAGGAGGATTTTCTCTGAGAATTTGGGAGAACTGGCTG |
| CKAP5 | 69 | 10 | TTGCATTGCTTCAGTTTCTCTTTGGG | TGTGGAGGGATTGAAGGATA | TGTGGAGGGATTGAAGGATA | TGTGGAGGGATTGAAGGATATTTTGTGGAGGGATTGAAGGATATTTTTGCATTGCTTCAGTTTCTCTTTGGG |
| CKAP5 | 70 | 10 | GTCTTTAGGGTCCTTATGGGAAATTCA | GGGATGTATTGAAGGAGGAT | GGGATGTATTGAAGGAGGAT | GGGATGTATTGAAGGAGGATTTTGGGATGTATTGAAGGAGGATTTTGTCTTTAGGGTCCTTATGGGAAATTCA |
| CKAP5 | 71 | 10 | ATCATCGTTAGGTGGTCCAGGATCTT | TGTGGAGGGATTGAAGGATA | TGTGGAGGGATTGAAGGATA | TGTGGAGGGATTGAAGGATATTTTGTGGAGGGATTGAAGGATATTTATCATCGTTAGGTGGTCCAGGATCTT |
| CKAP5 | 72 | 10 | ATGGGCCTCCAGCTCAGACTCGTTTT | GGGATGTATTGAAGGAGGAT | GGGATGTATTGAAGGAGGAT | GGGATGTATTGAAGGAGGATTTTGGGATGTATTGAAGGAGGATTTTATGGGCCTCCAGCTCAGACTCGTTTT |
| CKAP5 | 73 | 10 | TTTCCTTATCAGACTTGCTCCCAGTC | TGTGGAGGGATTGAAGGATA | TGTGGAGGGATTGAAGGATA | TGTGGAGGGATTGAAGGATATTTTGTGGAGGGATTGAAGGATATTTTTTCCTTATCAGACTTGCTCCCAGTC |
| CKAP5 | 74 | 10 | AAATCATTCACTTTGGCCTTTGATGA | GGGATGTATTGAAGGAGGAT | GGGATGTATTGAAGGAGGAT | GGGATGTATTGAAGGAGGATTTTGGGATGTATTGAAGGAGGATTTTAAATCATTCACTTTGGCCTTTGATGA |
| CKAP5 | 75 | 10 | GCTCTGGAAGAACTGTGAGGAATTTT | TGTGGAGGGATTGAAGGATA | TGTGGAGGGATTGAAGGATA | TGTGGAGGGATTGAAGGATATTTTGTGGAGGGATTGAAGGATATTTGCTCTGGAAGAACTGTGAGGAATTTT |
| CKAP5 | 76 | 10 | TCAATCACCCGAAGGCCTCTTTCGAC | GGGATGTATTGAAGGAGGAT | GGGATGTATTGAAGGAGGAT | GGGATGTATTGAAGGAGGATTTTGGGATGTATTGAAGGAGGATTTTTCAATCACCCGAAGGCCTCTTTCGAC |
| CKAP5 | 77 | 10 | GCACACATGTGACTTCCATCTGAGGG | TGTGGAGGGATTGAAGGATA | TGTGGAGGGATTGAAGGATA | TGTGGAGGGATTGAAGGATATTTTGTGGAGGGATTGAAGGATATTTGCACACATGTGACTTCCATCTGAGGG |
| CKAP5 | 78 | 10 | AGCCTTTCCAAGTAGACAGATGGCCC | GGGATGTATTGAAGGAGGAT | GGGATGTATTGAAGGAGGAT | GGGATGTATTGAAGGAGGATTTTGGGATGTATTGAAGGAGGATTTTAGCCTTTCCAAGTAGACAGATGGCCC |
| CKAP5 | 79 | 10 | GCATGTCTGTGGAAGAGGCGACAGTA | TGTGGAGGGATTGAAGGATA | TGTGGAGGGATTGAAGGATA | TGTGGAGGGATTGAAGGATATTTTGTGGAGGGATTGAAGGATATTTGCATGTCTGTGGAAGAGGCGACAGTA |
| CKAP5 | 80 | 10 | CTCCCGGAGCTGAGAGAGTTTGCTGT | GGGATGTATTGAAGGAGGAT | GGGATGTATTGAAGGAGGAT | GGGATGTATTGAAGGAGGATTTTGGGATGTATTGAAGGAGGATTTTCTCCCGGAGCTGAGAGAGTTTGCTGT |
| NCOA | 1 | 11 | GCGATCCTTTCCTCTTATGTGAGTCT | TGTGGAGGGATTGAAGGATA | TGTGGAGGGATTGAAGGATA | TGTGGAGGGATTGAAGGATATTTTGTGGAGGGATTGAAGGATATTTGCGATCCTTTCCTCTTATGTGAGTCT |
| NCOA | 2 | 11 | TTTATTTTCTTGCTCCCTGCGCCTC | ATAGGAAATGGTGGTAGTGT | ATAGGAAATGGTGGTAGTGT | ATAGGAAATGGTGGTAGTGTTTTATAGGAAATGGTGGTAGTGTTTTTTTATTTTCTTGCTCCCTGCGCCTC |
| NCOA | 3 | 11 | ACTCAAGCTGTCAATGTCACTAATGT | TGTGGAGGGATTGAAGGATA | TGTGGAGGGATTGAAGGATA | TGTGGAGGGATTGAAGGATATTTTGTGGAGGGATTGAAGGATATTTACTCAAGCTGTCAATGTCACTAATGT |
| NCOA | 4 | 11 | CGACTGTTTTCTTCAAAATCTTGCATTTG | ATAGGAAATGGTGGTAGTGT | ATAGGAAATGGTGGTAGTGT | ATAGGAAATGGTGGTAGTGTTTTATAGGAAATGGTGGTAGTGTTTTCGACTGTTTTCTTCAAAATCTTGCATTTG |
| NCOA | 5 | 11 | TGTACATCGTCATCAGTTGTTGATTTC | TGTGGAGGGATTGAAGGATA | TGTGGAGGGATTGAAGGATA | TGTGGAGGGATTGAAGGATATTTTGTGGAGGGATTGAAGGATATTTTGTACATCGTCATCAGTTGTTGATTTC |
| NCOA | 6 | 11 | CCTTGACTACTTGATGAGATGTCTGAT | ATAGGAAATGGTGGTAGTGT | ATAGGAAATGGTGGTAGTGT | ATAGGAAATGGTGGTAGTGTTTTATAGGAAATGGTGGTAGTGTTTTCCTTGACTACTTGATGAGATGTCTGAT |
| NCOA | 7 | 11 | ATTCTCCCTTCACAGTTCACAACAA | TGTGGAGGGATTGAAGGATA | TGTGGAGGGATTGAAGGATA | TGTGGAGGGATTGAAGGATATTTTGTGGAGGGATTGAAGGATATTTATTCTCCCTTCACAGTTCACAACAA |
| NCOA | 8 | 11 | AGCTGGTTACATTCTCTGACACAAAT | ATAGGAAATGGTGGTAGTGT | ATAGGAAATGGTGGTAGTGT | ATAGGAAATGGTGGTAGTGTTTTATAGGAAATGGTGGTAGTGTTTTAGCTGGTTACATTCTCTGACACAAAT |
| NCOA | 9 | 11 | TTCTTCACAAATTCTGCATGATCCC | TGTGGAGGGATTGAAGGATA | TGTGGAGGGATTGAAGGATA | TGTGGAGGGATTGAAGGATATTTTGTGGAGGGATTGAAGGATATTTTTCTTCACAAATTCTGCATGATCCC |
| NCOA | 10 | 11 | TAGCATCCTGCAGTTAAAGGTATGG | ATAGGAAATGGTGGTAGTGT | ATAGGAAATGGTGGTAGTGT | ATAGGAAATGGTGGTAGTGTTTTATAGGAAATGGTGGTAGTGTTTTTAGCATCCTGCAGTTAAAGGTATGG |
| NCOA | 11 | 11 | TACTTCATAACGCTGGCAAGCTTCT | TGTGGAGGGATTGAAGGATA | TGTGGAGGGATTGAAGGATA | TGTGGAGGGATTGAAGGATATTTTGTGGAGGGATTGAAGGATATTTTACTTCATAACGCTGGCAAGCTTCT |
| NCOA | 12 | 11 | TTGGCTGTGACACAGTGAAACACTG | ATAGGAAATGGTGGTAGTGT | ATAGGAAATGGTGGTAGTGT | ATAGGAAATGGTGGTAGTGTTTTATAGGAAATGGTGGTAGTGTTTTTTGGCTGTGACACAGTGAAACACTG |
| NCOA | 13 | 11 | GTAGTATCTTGCTTGGTCATAAAGGATTC | TGTGGAGGGATTGAAGGATA | TGTGGAGGGATTGAAGGATA | TGTGGAGGGATTGAAGGATATTTTGTGGAGGGATTGAAGGATATTTGTAGTATCTTGCTTGGTCATAAAGGATTC |
| NCOA | 14 | 11 | TCTCAGGGAACTAGTATCAATAGAGATGAT | ATAGGAAATGGTGGTAGTGT | ATAGGAAATGGTGGTAGTGT | ATAGGAAATGGTGGTAGTGTTTTATAGGAAATGGTGGTAGTGTTTTTCTCAGGGAACTAGTATCAATAGAGATGAT |
| NCOA | 15 | 11 | GGTTCTCTGCCCTGAGGTTGGAAAA | TGTGGAGGGATTGAAGGATA | TGTGGAGGGATTGAAGGATA | TGTGGAGGGATTGAAGGATATTTTGTGGAGGGATTGAAGGATATTTGGTTCTCTGCCCTGAGGTTGGAAAA |
| NCOA | 16 | 11 | TTCTTGGAACAGCTGTCTGGCATAA | ATAGGAAATGGTGGTAGTGT | ATAGGAAATGGTGGTAGTGT | ATAGGAAATGGTGGTAGTGTTTTATAGGAAATGGTGGTAGTGTTTTTTCTTGGAACAGCTGTCTGGCATAA |
| NCOA | 17 | 11 | ATGTCTGGACTTTGAGGGTAGCAAA | TGTGGAGGGATTGAAGGATA | TGTGGAGGGATTGAAGGATA | TGTGGAGGGATTGAAGGATATTTTGTGGAGGGATTGAAGGATATTTATGTCTGGACTTTGAGGGTAGCAAA |
| NCOA | 18 | 11 | CGATGATATGAATTCCCATGATGAAAGGT | ATAGGAAATGGTGGTAGTGT | ATAGGAAATGGTGGTAGTGT | ATAGGAAATGGTGGTAGTGTTTTATAGGAAATGGTGGTAGTGTTTTCGATGATATGAATTCCCATGATGAAAGGT |
| NCOA | 19 | 11 | GGGAATTGACATTCCAGAATTAGTGTCA | TGTGGAGGGATTGAAGGATA | TGTGGAGGGATTGAAGGATA | TGTGGAGGGATTGAAGGATATTTTGTGGAGGGATTGAAGGATATTTGGGAATTGACATTCCAGAATTAGTGTCA |
| NCOA | 20 | 11 | GATACTAGGATTGACCGAGGGATTTAC | ATAGGAAATGGTGGTAGTGT | ATAGGAAATGGTGGTAGTGT | ATAGGAAATGGTGGTAGTGTTTTATAGGAAATGGTGGTAGTGTTTTGATACTAGGATTGACCGAGGGATTTAC |
| NCOA | 21 | 11 | ATGTTGCTGTTGGATGGTGGCAATG | TGTGGAGGGATTGAAGGATA | TGTGGAGGGATTGAAGGATA | TGTGGAGGGATTGAAGGATATTTTGTGGAGGGATTGAAGGATATTTATGTTGCTGTTGGATGGTGGCAATG |
| NCOA | 22 | 11 | CTGCTGGCGGTTTATTCTGGTGGAT | ATAGGAAATGGTGGTAGTGT | ATAGGAAATGGTGGTAGTGT | ATAGGAAATGGTGGTAGTGTTTTATAGGAAATGGTGGTAGTGTTTTCTGCTGGCGGTTTATTCTGGTGGAT |
| NCOA | 23 | 11 | AAACTTCCTTGGCTGTTGCTAGAAT | TGTGGAGGGATTGAAGGATA | TGTGGAGGGATTGAAGGATA | TGTGGAGGGATTGAAGGATATTTTGTGGAGGGATTGAAGGATATTTAAACTTCCTTGGCTGTTGCTAGAAT |
| NCOA | 24 | 11 | TACAATCTGACTTCCGGGTGAGCAT | ATAGGAAATGGTGGTAGTGT | ATAGGAAATGGTGGTAGTGT | ATAGGAAATGGTGGTAGTGTTTTATAGGAAATGGTGGTAGTGTTTTTACAATCTGACTTCCGGGTGAGCAT |
| NCOA | 25 | 11 | AAAGAGGGATTACTGCTCTGTGAAC | TGTGGAGGGATTGAAGGATA | TGTGGAGGGATTGAAGGATA | TGTGGAGGGATTGAAGGATATTTTGTGGAGGGATTGAAGGATATTTAAAGAGGGATTACTGCTCTGTGAAC |
| NCOA | 26 | 11 | TGTACCTTCCATAGGAGAATTATTGAGG | ATAGGAAATGGTGGTAGTGT | ATAGGAAATGGTGGTAGTGT | ATAGGAAATGGTGGTAGTGTTTTATAGGAAATGGTGGTAGTGTTTTTGTACCTTCCATAGGAGAATTATTGAGG |
| NCOA | 27 | 11 | GGAATTGTTTGGCATCCTGGGCCTT | TGTGGAGGGATTGAAGGATA | TGTGGAGGGATTGAAGGATA | TGTGGAGGGATTGAAGGATATTTTGTGGAGGGATTGAAGGATATTTGGAATTGTTTGGCATCCTGGGCCTT |
| NCOA | 28 | 11 | GGAGAGCTTAATGTCGAAATATTAGGAGG | ATAGGAAATGGTGGTAGTGT | ATAGGAAATGGTGGTAGTGT | ATAGGAAATGGTGGTAGTGTTTTATAGGAAATGGTGGTAGTGTTTTGGAGAGCTTAATGTCGAAATATTAGGAGG |
| NCOA | 29 | 11 | CTGGGATGTTTGAATAAGATCGGTTATT | TGTGGAGGGATTGAAGGATA | TGTGGAGGGATTGAAGGATA | TGTGGAGGGATTGAAGGATATTTTGTGGAGGGATTGAAGGATATTTCTGGGATGTTTGAATAAGATCGGTTATT |
| NCOA | 30 | 11 | GGTCCTTCATTCATACCCTGTAAAGATG | ATAGGAAATGGTGGTAGTGT | ATAGGAAATGGTGGTAGTGT | ATAGGAAATGGTGGTAGTGTTTTATAGGAAATGGTGGTAGTGTTTTGGTCCTTCATTCATACCCTGTAAAGATG |
| NCOA | 31 | 11 | GTGAGCTCATCTGCCTGAGGACTGG | TGTGGAGGGATTGAAGGATA | TGTGGAGGGATTGAAGGATA | TGTGGAGGGATTGAAGGATATTTTGTGGAGGGATTGAAGGATATTTGTGAGCTCATCTGCCTGAGGACTGG |
| NCOA | 32 | 11 | TGCTGGTTGTATATTTAATCTGCTAGGT | ATAGGAAATGGTGGTAGTGT | ATAGGAAATGGTGGTAGTGT | ATAGGAAATGGTGGTAGTGTTTTATAGGAAATGGTGGTAGTGTTTTTGCTGGTTGTATATTTAATCTGCTAGGT |
| NCOA | 33 | 11 | GAGCTGTTGTCAGATTGAATCATTTCATT | TGTGGAGGGATTGAAGGATA | TGTGGAGGGATTGAAGGATA | TGTGGAGGGATTGAAGGATATTTTGTGGAGGGATTGAAGGATATTTGAGCTGTTGTCAGATTGAATCATTTCATT |
| NCOA | 34 | 11 | CCCTGAATCCAGAGGTTTGCCATCA | ATAGGAAATGGTGGTAGTGT | ATAGGAAATGGTGGTAGTGT | ATAGGAAATGGTGGTAGTGTTTTATAGGAAATGGTGGTAGTGTTTTCCCTGAATCCAGAGGTTTGCCATCA |
| NCOA | 35 | 11 | TGTGACTGGTTTGAGAGTATTTACTGTC | TGTGGAGGGATTGAAGGATA | TGTGGAGGGATTGAAGGATA | TGTGGAGGGATTGAAGGATATTTTGTGGAGGGATTGAAGGATATTTTGTGACTGGTTTGAGAGTATTTACTGTC |
| NCOA | 36 | 11 | GCAGTTGTTGTCAAAAGCTGCACTA | ATAGGAAATGGTGGTAGTGT | ATAGGAAATGGTGGTAGTGT | ATAGGAAATGGTGGTAGTGTTTTATAGGAAATGGTGGTAGTGTTTTGCAGTTGTTGTCAAAAGCTGCACTA |
| NCOA | 37 | 11 | AAGACAGGACATCTTTGCAGCTTGT | TGTGGAGGGATTGAAGGATA | TGTGGAGGGATTGAAGGATA | TGTGGAGGGATTGAAGGATATTTTGTGGAGGGATTGAAGGATATTTAAGACAGGACATCTTTGCAGCTTGT |
| NCOA | 38 | 11 | AGAGGGACAAGAACCTCCTGAAGAG | ATAGGAAATGGTGGTAGTGT | ATAGGAAATGGTGGTAGTGT | ATAGGAAATGGTGGTAGTGTTTTATAGGAAATGGTGGTAGTGTTTTAGAGGGACAAGAACCTCCTGAAGAG |
| NCOA | 39 | 11 | CCCTCCTGTAAGAGCCGGTGTAGAA | TGTGGAGGGATTGAAGGATA | TGTGGAGGGATTGAAGGATA | TGTGGAGGGATTGAAGGATATTTTGTGGAGGGATTGAAGGATATTTCCCTCCTGTAAGAGCCGGTGTAGAA |
| NCOA | 40 | 11 | CAGACAAAGTGGTGATATCTGAGGG | ATAGGAAATGGTGGTAGTGT | ATAGGAAATGGTGGTAGTGT | ATAGGAAATGGTGGTAGTGTTTTATAGGAAATGGTGGTAGTGTTTTCAGACAAAGTGGTGATATCTGAGGG |
| NCOA | 41 | 11 | GTTTCCTTGTACCTGTCCAGTCACT | TGTGGAGGGATTGAAGGATA | TGTGGAGGGATTGAAGGATA | TGTGGAGGGATTGAAGGATATTTTGTGGAGGGATTGAAGGATATTTGTTTCCTTGTACCTGTCCAGTCACT |
| NCOA | 42 | 11 | GGAGCTGATGGTCTTTTGATTCTTTT | ATAGGAAATGGTGGTAGTGT | ATAGGAAATGGTGGTAGTGT | ATAGGAAATGGTGGTAGTGTTTTATAGGAAATGGTGGTAGTGTTTTGGAGCTGATGGTCTTTTGATTCTTTT |
| NCOA | 43 | 11 | CCACTTTCACCTTTACATCATCCAGG | TGTGGAGGGATTGAAGGATA | TGTGGAGGGATTGAAGGATA | TGTGGAGGGATTGAAGGATATTTTGTGGAGGGATTGAAGGATATTTCCACTTTCACCTTTACATCATCCAGG |
| NCOA | 44 | 11 | TGTATTACATGGATCCATCTGTTCTTTCT | ATAGGAAATGGTGGTAGTGT | ATAGGAAATGGTGGTAGTGT | ATAGGAAATGGTGGTAGTGTTTTATAGGAAATGGTGGTAGTGTTTTTGTATTACATGGATCCATCTGTTCTTTCT |
| NCOA | 45 | 11 | ACTGGCTCTGGGCCTCCAGTTTTAT | TGTGGAGGGATTGAAGGATA | TGTGGAGGGATTGAAGGATA | TGTGGAGGGATTGAAGGATATTTTGTGGAGGGATTGAAGGATATTTACTGGCTCTGGGCCTCCAGTTTTAT |
| NCOA | 46 | 11 | TGATCAAACTGGTCAAGGTCAGCTG | ATAGGAAATGGTGGTAGTGT | ATAGGAAATGGTGGTAGTGT | ATAGGAAATGGTGGTAGTGTTTTATAGGAAATGGTGGTAGTGTTTTTGATCAAACTGGTCAAGGTCAGCTG |
| NCOA | 47 | 11 | ACCGCACCATCCATCCTGTCTGTCT | TGTGGAGGGATTGAAGGATA | TGTGGAGGGATTGAAGGATA | TGTGGAGGGATTGAAGGATATTTTGTGGAGGGATTGAAGGATATTTACCGCACCATCCATCCTGTCTGTCT |
| NCOA | 48 | 11 | GGATCTCCGATTTGATGGTTACACTG | ATAGGAAATGGTGGTAGTGT | ATAGGAAATGGTGGTAGTGT | ATAGGAAATGGTGGTAGTGTTTTATAGGAAATGGTGGTAGTGTTTTGGATCTCCGATTTGATGGTTACACTG |
| NCOA | 49 | 11 | GATCGCCTGTTCCTGGTTGTCCAAA | TGTGGAGGGATTGAAGGATA | TGTGGAGGGATTGAAGGATA | TGTGGAGGGATTGAAGGATATTTTGTGGAGGGATTGAAGGATATTTGATCGCCTGTTCCTGGTTGTCCAAA |
| NCOA | 50 | 11 | GCTGTCACTGTATTATTTGTCCATGGAA | ATAGGAAATGGTGGTAGTGT | ATAGGAAATGGTGGTAGTGT | ATAGGAAATGGTGGTAGTGTTTTATAGGAAATGGTGGTAGTGTTTTGCTGTCACTGTATTATTTGTCCATGGAA |
| NCOA | 51 | 11 | CCAGCTGTTCAAGAAGAGCCTTCTC | TGTGGAGGGATTGAAGGATA | TGTGGAGGGATTGAAGGATA | TGTGGAGGGATTGAAGGATATTTTGTGGAGGGATTGAAGGATATTTCCAGCTGTTCAAGAAGAGCCTTCTC |

| | | | | | | |
|---|---|---|---|---|---|---|
| NCOA | 52 | 11 | GTTTCATCTTTGCCACTAAGGAAGG | ATAGGAAATGGTGGTAGTGT | ATAGGAAATGGTGGTAGTGT | ATAGGAAATGGTGGTAGTGTTTTATAGGAAATGGTGGTAGTGTTTTGTTTCATCTTTGCCACTAAGGAAGG |
| NCOA | 53 | 11 | ATCTCTCTGATAATACATCTAATCCACCC | TGTGGAGGGATTGAAGGATA | TGTGGAGGGATTGAAGGATA | TGTGGAGGGATTGAAGGATATTTTGTGGAGGGATTGAAGGATATTTATCTCTCTGATAATACATCTAATCCACCC |
| NCOA | 54 | 11 | AAAGGTGGCGTTGCTTGTTGTGGTG | ATAGGAAATGGTGGTAGTGT | ATAGGAAATGGTGGTAGTGT | ATAGGAAATGGTGGTAGTGTTTTATAGGAAATGGTGGTAGTGTTTTAAAGGTGGCGTTGCTTGTTGTGGTG |
| NCOA | 55 | 11 | TGCCTTGGAAAGGGCTAGGGAGATT | TGTGGAGGGATTGAAGGATA | TGTGGAGGGATTGAAGGATA | TGTGGAGGGATTGAAGGATATTTTGTGGAGGGATTGAAGGATATTTTGCCTTGGAAAGGGCTAGGGAGATT |
| NCOA | 56 | 11 | GTGTTACTTGAACAGGCATCGTCCC | ATAGGAAATGGTGGTAGTGT | ATAGGAAATGGTGGTAGTGT | ATAGGAAATGGTGGTAGTGTTTTATAGGAAATGGTGGTAGTGTTTTGTGTTACTTGAACAGGCATCGTCCC |
| NCOA | 57 | 11 | GCCGGAGGTCTGTTTAGAGTTTGCC | TGTGGAGGGATTGAAGGATA | TGTGGAGGGATTGAAGGATA | TGTGGAGGGATTGAAGGATATTTTGTGGAGGGATTGAAGGATATTTGCCGGAGGTCTGTTTAGAGTTTGCC |
| NCOA | 58 | 11 | TAGTTGAAGTCGAAGCTGGTTAGGT | ATAGGAAATGGTGGTAGTGT | ATAGGAAATGGTGGTAGTGT | ATAGGAAATGGTGGTAGTGTTTTATAGGAAATGGTGGTAGTGTTTTTAGTTGAAGTCGAAGCTGGTTAGGT |
| NCOA | 59 | 11 | AAGATAGCTTGCCGATTTTGGTGTA | TGTGGAGGGATTGAAGGATA | TGTGGAGGGATTGAAGGATA | TGTGGAGGGATTGAAGGATATTTTGTGGAGGGATTGAAGGATATTTAAGATAGCTTGCCGATTTTGGTGTA |
| NCOA | 60 | 11 | TGTTGCATGCCTGATCTCATATTGAT | ATAGGAAATGGTGGTAGTGT | ATAGGAAATGGTGGTAGTGT | ATAGGAAATGGTGGTAGTGTTTTATAGGAAATGGTGGTAGTGTTTTTGTTGCATGCCTGATCTCATATTGAT |
| NCOA | 61 | 11 | CTGTTGACTGTACAGTTCCCGCTGA | TGTGGAGGGATTGAAGGATA | TGTGGAGGGATTGAAGGATA | TGTGGAGGGATTGAAGGATATTTTGTGGAGGGATTGAAGGATATTTCTGTTGACTGTACAGTTCCCGCTGA |
| NCOA | 62 | 11 | GCTGCTGTATTAGCTGCCTCTGTCG | ATAGGAAATGGTGGTAGTGT | ATAGGAAATGGTGGTAGTGT | ATAGGAAATGGTGGTAGTGTTTTATAGGAAATGGTGGTAGTGTTTTGCTGCTGTATTAGCTGCCTCTGTCG |
| NCOA | 63 | 11 | AGATGAGGGAGGGAGGTTGTTCCCA | TGTGGAGGGATTGAAGGATA | TGTGGAGGGATTGAAGGATA | TGTGGAGGGATTGAAGGATATTTTGTGGAGGGATTGAAGGATATTTAGATGAGGGAGGGAGGTTGTTCCCA |
| NCOA | 64 | 11 | TGTGGAGCACCCTGAGGAAGACGGG | ATAGGAAATGGTGGTAGTGT | ATAGGAAATGGTGGTAGTGT | ATAGGAAATGGTGGTAGTGTTTTATAGGAAATGGTGGTAGTGTTTTTGTGGAGCACCCTGAGGAAGACGGG |
| NCOA | 65 | 11 | AAACGGGCTGGTAGAAGCAGGTGGG | TGTGGAGGGATTGAAGGATA | TGTGGAGGGATTGAAGGATA | TGTGGAGGGATTGAAGGATATTTTGTGGAGGGATTGAAGGATATTTAAACGGGCTGGTAGAAGCAGGTGGG |
| NCOA | 66 | 11 | ATGCTTCAGGATTTGCTGCTAGTTG | ATAGGAAATGGTGGTAGTGT | ATAGGAAATGGTGGTAGTGT | ATAGGAAATGGTGGTAGTGTTTTATAGGAAATGGTGGTAGTGTTTTATGCTTCAGGATTTGCTGCTAGTTG |
| NCOA | 67 | 11 | ACTGTCCTCCTATGTTTCCTGTCAT | TGTGGAGGGATTGAAGGATA | TGTGGAGGGATTGAAGGATA | TGTGGAGGGATTGAAGGATATTTTGTGGAGGGATTGAAGGATATTTACTGTCCTCCTATGTTTCCTGTCAT |
| NCOA | 68 | 11 | TGCATCTGAGGATTGATTCCAGTGC | ATAGGAAATGGTGGTAGTGT | ATAGGAAATGGTGGTAGTGT | ATAGGAAATGGTGGTAGTGTTTTATAGGAAATGGTGGTAGTGTTTTTGCATCTGAGGATTGATTCCAGTGC |
| NCOA | 69 | 11 | GGAGCAAAGTTGGCCTCACCTTGGG | TGTGGAGGGATTGAAGGATA | TGTGGAGGGATTGAAGGATA | TGTGGAGGGATTGAAGGATATTTTGTGGAGGGATTGAAGGATATTTGGAGCAAAGTTGGCCTCACCTTGGG |
| NCOA | 70 | 11 | CACCATGGAGCTCCCAGGGCTTAGA | ATAGGAAATGGTGGTAGTGT | ATAGGAAATGGTGGTAGTGT | ATAGGAAATGGTGGTAGTGTTTTATAGGAAATGGTGGTAGTGTTTTCACCATGGAGCTCCCAGGGCTTAGA |
| NCOA | 71 | 11 | GAGGCAGGTGGAGTTTGCTGGAGAA | TGTGGAGGGATTGAAGGATA | TGTGGAGGGATTGAAGGATA | TGTGGAGGGATTGAAGGATATTTTGTGGAGGGATTGAAGGATATTTGAGGCAGGTGGAGTTTGCTGGAGAA |
| NCOA | 72 | 11 | GGCCTTCATGTCTGGTGACTGATAC | ATAGGAAATGGTGGTAGTGT | ATAGGAAATGGTGGTAGTGT | ATAGGAAATGGTGGTAGTGTTTTATAGGAAATGGTGGTAGTGTTTTGGCCTTCATGTCTGGTGACTGATAC |
| NCOA | 73 | 11 | TGGTTCTGGACAGCTTGACTGAACA | TGTGGAGGGATTGAAGGATA | TGTGGAGGGATTGAAGGATA | TGTGGAGGGATTGAAGGATATTTTGTGGAGGGATTGAAGGATATTTTGGTTCTGGACAGCTTGACTGAACA |
| NCOA | 74 | 11 | GTATACTCCTGGCTGTGCAGGCGTG | ATAGGAAATGGTGGTAGTGT | ATAGGAAATGGTGGTAGTGT | ATAGGAAATGGTGGTAGTGTTTTATAGGAAATGGTGGTAGTGTTTTGTATACTCCTGGCTGTGCAGGCGTG |
| NCOA | 75 | 11 | TCATGTTCTGAACATTCGTATTTCCAC | TGTGGAGGGATTGAAGGATA | TGTGGAGGGATTGAAGGATA | TGTGGAGGGATTGAAGGATATTTTGTGGAGGGATTGAAGGATATTTTCATGTTCTGAACATTCGTATTTCCAC |
| NCOA | 76 | 11 | CTCATCTGCATCTGGGCCATCATTG | ATAGGAAATGGTGGTAGTGT | ATAGGAAATGGTGGTAGTGT | ATAGGAAATGGTGGTAGTGTTTTATAGGAAATGGTGGTAGTGTTTTCTCATCTGCATCTGGGCCATCATTG |
| NCOA | 77 | 11 | TTGCAGTAGAGGCCTGTGTGTCTCA | TGTGGAGGGATTGAAGGATA | TGTGGAGGGATTGAAGGATA | TGTGGAGGGATTGAAGGATATTTTGTGGAGGGATTGAAGGATATTTTTGCAGTAGAGGCCTGTGTGTCTCA |
| NCOA | 78 | 11 | TTTGAGAAGGTCAGTGGATGAGAGC | ATAGGAAATGGTGGTAGTGT | ATAGGAAATGGTGGTAGTGT | ATAGGAAATGGTGGTAGTGTTTTATAGGAAATGGTGGTAGTGTTTTTTTGAGAAGGTCAGTGGATGAGAGC |
| NCOA | 79 | 11 | CCAGGCTGGTTCAGGTAAGGGTCCC | TGTGGAGGGATTGAAGGATA | TGTGGAGGGATTGAAGGATA | TGTGGAGGGATTGAAGGATATTTTGTGGAGGGATTGAAGGATATTTCCAGGCTGGTTCAGGTAAGGGTCCC |
| NCOA | 80 | 11 | TGACGTGGGCTTTTGAGTTCCCAGT | ATAGGAAATGGTGGTAGTGT | ATAGGAAATGGTGGTAGTGT | ATAGGAAATGGTGGTAGTGTTTTATAGGAAATGGTGGTAGTGTTTTTGACGTGGGCTTTTGAGTTCCCAGT |

Phasor Transform Expressions

The first-order phasor transform of the lifetime intensity photon histogram $I(t)$:

$$S = \frac{\int_0^T I(t)\sin(\omega t)dt}{\int_0^T I(t)dt} \tag{1}$$

$$G = \frac{\int_0^T I(t)\cos(\omega t)\,dt}{\int_0^T I(t)\,dt} \tag{2}$$

Where $T$ is the period between excitation pulses (or modulation period), and $\omega = \frac{2\pi}{T}$ the pulsation frequency such that the period of trigonometric functions matches the excitation period $T$.

The first-order phasor transform of the spectral intensity photon histogram $I(\lambda)$:

$$S = \frac{\int_{\lambda_0}^{\lambda_f} I(\lambda)\sin(\omega\lambda - \omega\lambda_0)\,d\lambda}{\int_{\lambda_0}^{\lambda_f} I(\lambda)\,d\lambda} \tag{3}$$

$$G = \frac{\int_{\lambda_0}^{\lambda_f} I(\lambda)\cos(\omega\lambda - \omega\lambda_0)\,d\lambda}{\int_{\lambda_0}^{\lambda_f} I(\lambda)\,d\lambda} \tag{4}$$

Where $\lambda_0$ and $\lambda_f$ are the limits of the spectral band of the detector, and $\omega = \frac{2\pi}{\lambda_f - \lambda_0}$ the pulsation frequency such that the period of trigonometric functions matches the spectral bandwidth.

#### Figures

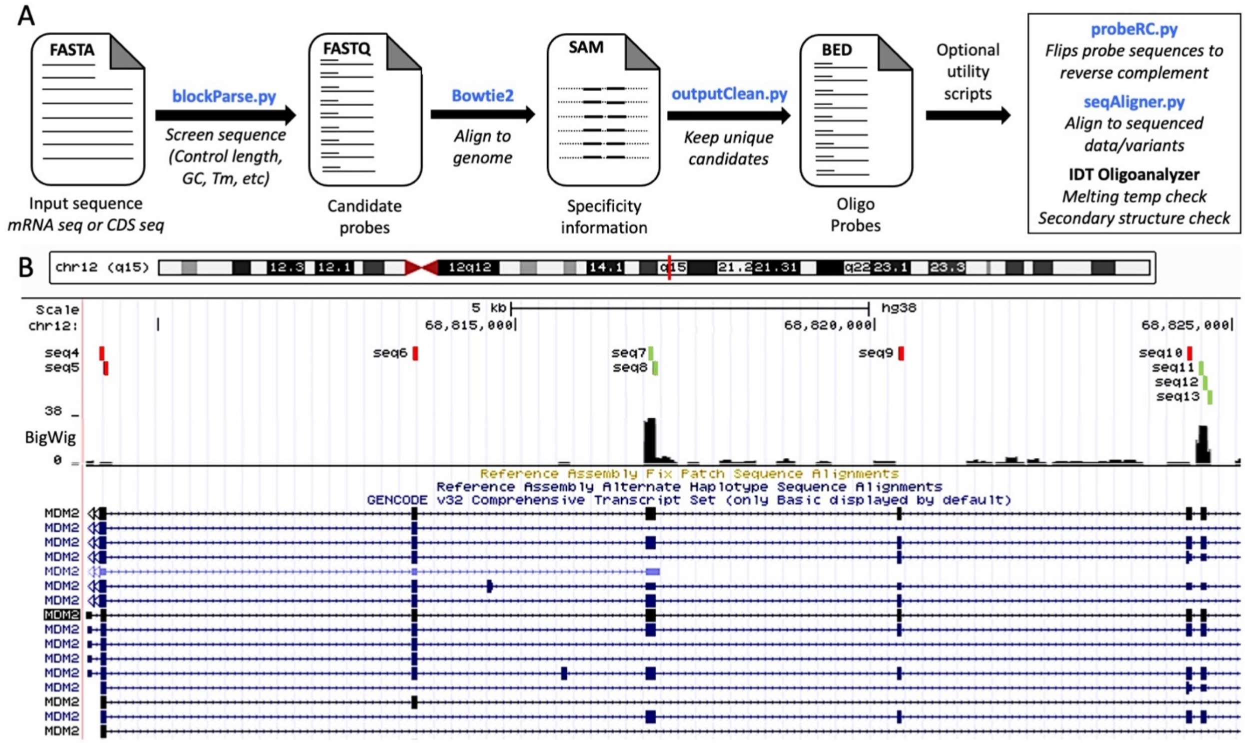


**Supplementary Figure 1. Schematic overview of probe design process. A)** A modified version of Oligominer was used to design primary probes. **B)** In the seqAligner.py script, the probes (labeled "seq#") go through BLAT and are aligned to the RNA sequencing data, as seen on the UCSC Genome Browser. The BigWig browser track shows a histogram of the read counts and probes are aligned and compared to the read counts of the region they overlap with. Probes that overlapped with more than 5% of the highest read count were used (green) and those that aligned with regions considered as "low read count" (5% or less) were removed from the final probe list (red).

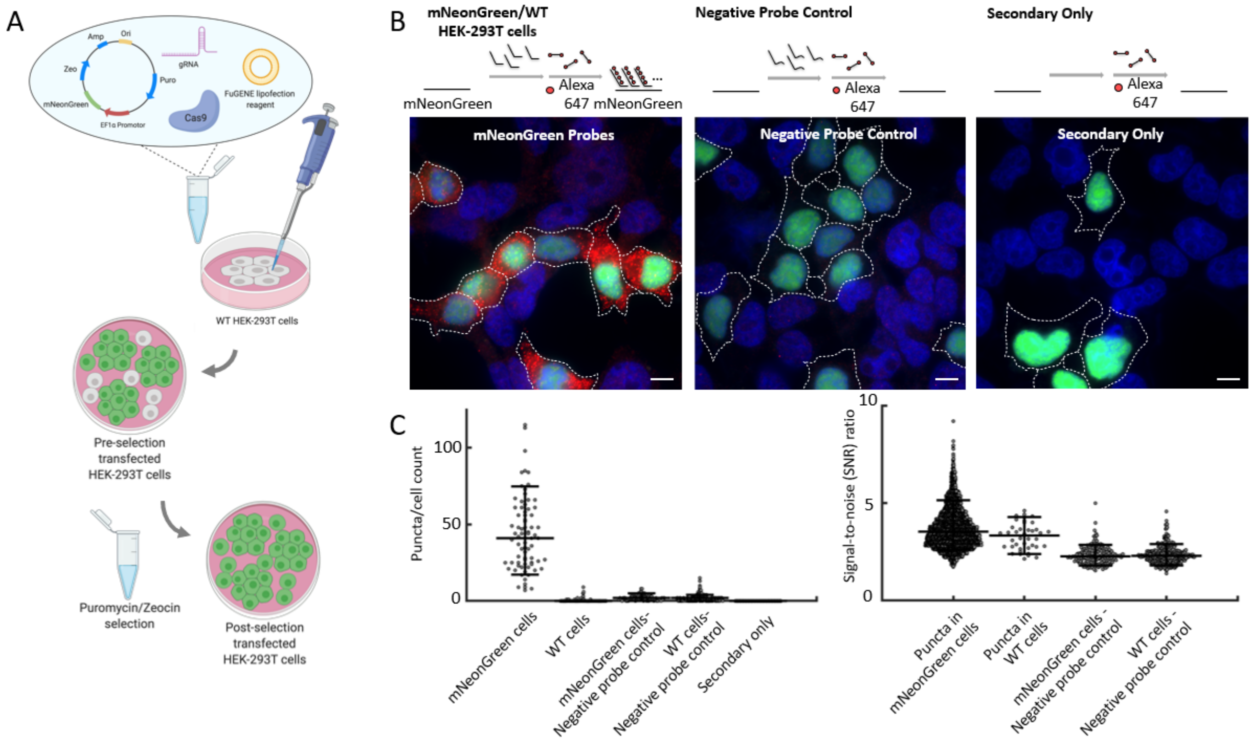


***Supplementary Figure 2. Validation of probe hybridization in mNeonGreen cells. A)*** *Schematic of engineering of mNeonGreen HEK293T-X cells. Engineered mNeonGreen plasmids were transfected into HEK293T-X cells with FuGENE HD Transfection Reagent. Three days after transfection, the cells were then selected with Puromycin and Zeocin.* ***B)*** *Schematic and representative images of each condition. The primary probes were designed to be complementary to mNeonGreen transcripts. A dopachrome tautomerase (DCT) primary probe negative control, which uses primary probes targeting sequences not present in the mNeonGreen HEK-293T-X cells but can still bind to secondary fluorescent oligonucleotides, was used to indicate any nonspecific binding which can occur with primary probe labeling. A negative control where only secondary probes were added but no primary probes were added was used as a reference for nonspecific binding from secondary probes alone. For each condition, the concentration of each primary probe (14 in total) was 1 nM and the secondary probe was 5 nM. Scale bar = 10 µm.* ***C)*** *Plots to quantify the detected puncta per cell and signal-to-noise (SNR) ratio under different conditions. Left, scatter plot showing puncta number per cell (n=755 cells). Right, signal-to-noise ratio (SNR). SNR=each signal intensity/the mean of background noise (n=3,860 puncta).*

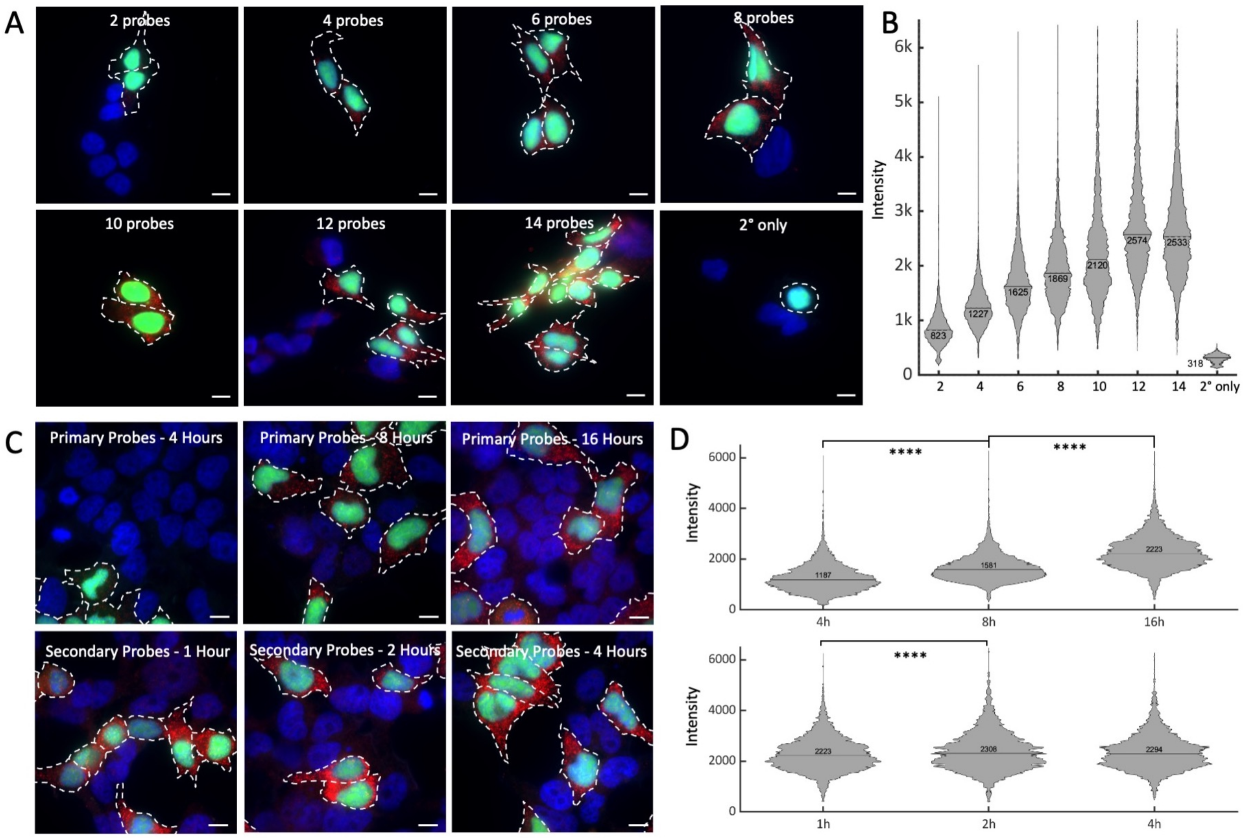


***Supplementary Figure 3. Optimization parameters of in situ hybridization conditions. A)*** *Representative images of mNeonGreen cells with different numbers of primary probes. Conditions included 2, 4, 6, 8, 10, 12, and 14 mNeonGreen primary probes to the HEK293T-X mNeonGreen and WT model. The concentration of each primary probe is constant (5 nM). Scale bars are 10 µm.* ***B)*** *Intensity distribution of detected puncta shows the effects of the number of primary probes on signal intensity (total n ≈ 64k puncta).* ***C)*** *Representative images of mNeonGreen cells with different incubation time of probes. The primary probes hybridization incubation times consisted of 4, 8, and 16 hours. For secondary fluorophore probes, incubation times tested were 1, 2, and 4 hours. Scale bars are 10 µm.* ***D)*** *Intensity distribution of detected puncta as a function of incubation time. Top, primary probe incubation time (total n ≈ 26k puncta). Bottom, secondary probe incubation time (total n ≈ 20k puncta). Pairwise t-test resulted in p-values <$10^4$ for conditions marked with ****.*

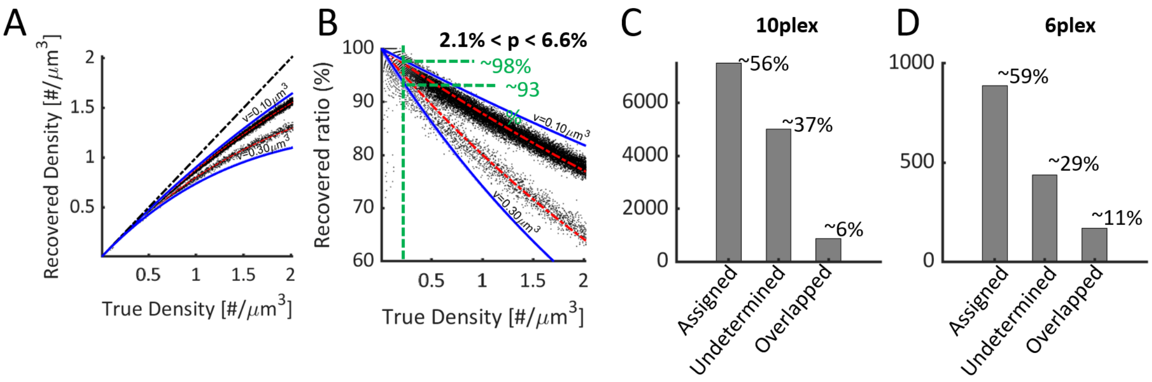


***Supplementary Figure 4. Overlapping and inconsistent signal simulations.*** *Simulations are run at different densities to generate 3D image stacks and puncta are detected using our image processing pipeline.* ***A)*** *Recovered density by our system as a function of the true density for the limit considered puncta volumes (blue) and simulation trends (top trend for 1plex bottom trend for 10-plex).* ***B)*** *Percentage of recovered puncta for the 1plex (top trend, depicting ratio of underestimated due to overlap) and 10-plex (bottom trend, depicting ratio of inconsistent signal due to overlap), again between the limit considered puncta volumes (blue).* ***C,D)*** *Number of puncta assigned to a particular gene, undetermined puncta and overlapped puncta for the 10-plex experiment (Figure 4 in the paper) and 6-plex experiment (Figure 5 in the paper).*

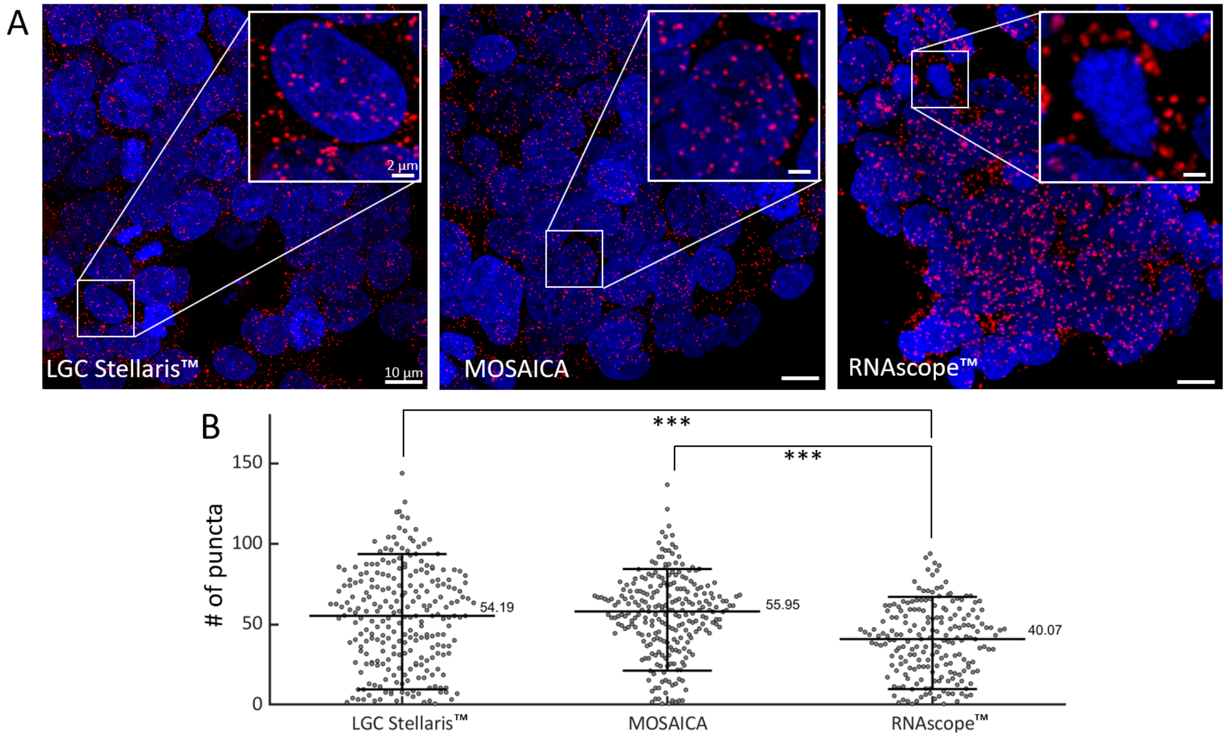


***Supplementary Figure 5.*** *Benchmarking MOSAICA against RNAscope$^{TM}$ and LGC Stellaris$^{TM}$.* ***A)*** *POLR2A gene expression on colorectal cancer SW480 cells following RNAscope$^{TM}$, LGC Stellaris$^{TM}$ and MOSAICA protocols. Scale bars are 10µm.* ***B)*** *Puncta counts per cell volume between three platforms. MOSAICA exhibited comparable puncta per cell count compared to benchmark LGC Stellaris$^{TM}$, whereas RNAscope$^{TM}$ was undercount. Pairwise t-test against null hypothesis that values belong to distributions of equal means were $p = 0.4$ (LGC Stellaris$^{TM}$ vs MOSAICA), $p = 3.4 \times 10^{-4}$ (LGC Stellaris$^{TM}$ vs RNAscope$^{TM}$ and $p = 7.8 \times 10^{-4}$ (MOSAICA vs RNAscope$^{TM}$). A sliding volume of 3000µm$^3$ was used throughout the image stacks and the number of puncta counts per volume was obtained. This number was then divided into the average number of cells per volume depending on the 3D segmentation of DAPI nuclei.*

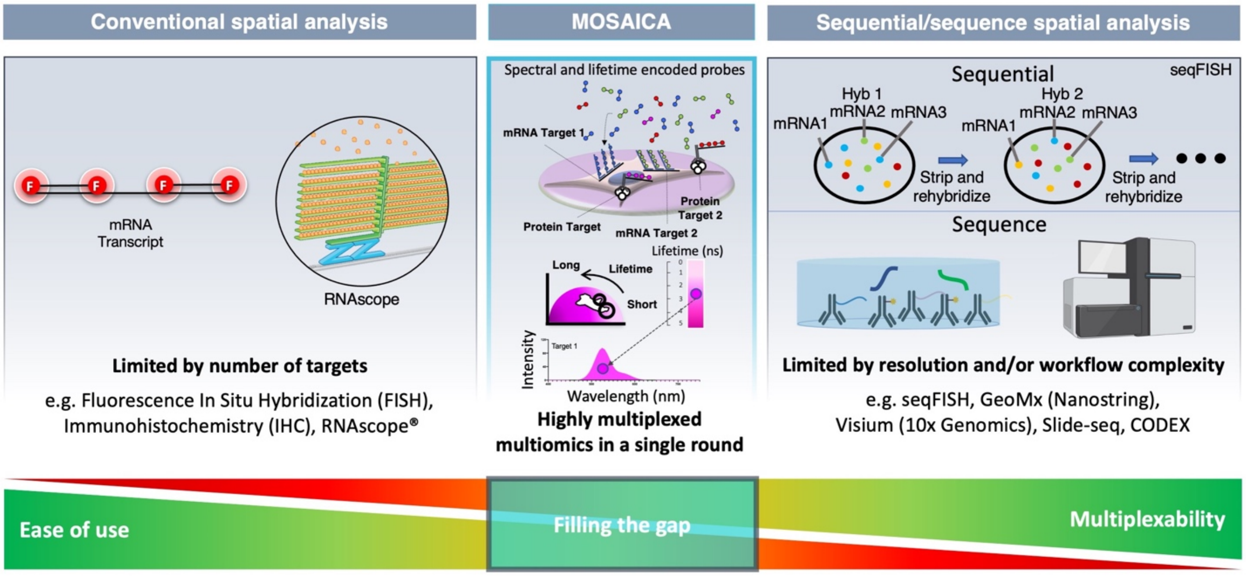


***Supplementary Figure 6.*** *Multiplexed spatial transcriptomics with MOSAICA, which is rapid, cost-effective, and easy-to-use, can fill a critical gap between conventional FISH and sequential- and sequencing-based techniques.*

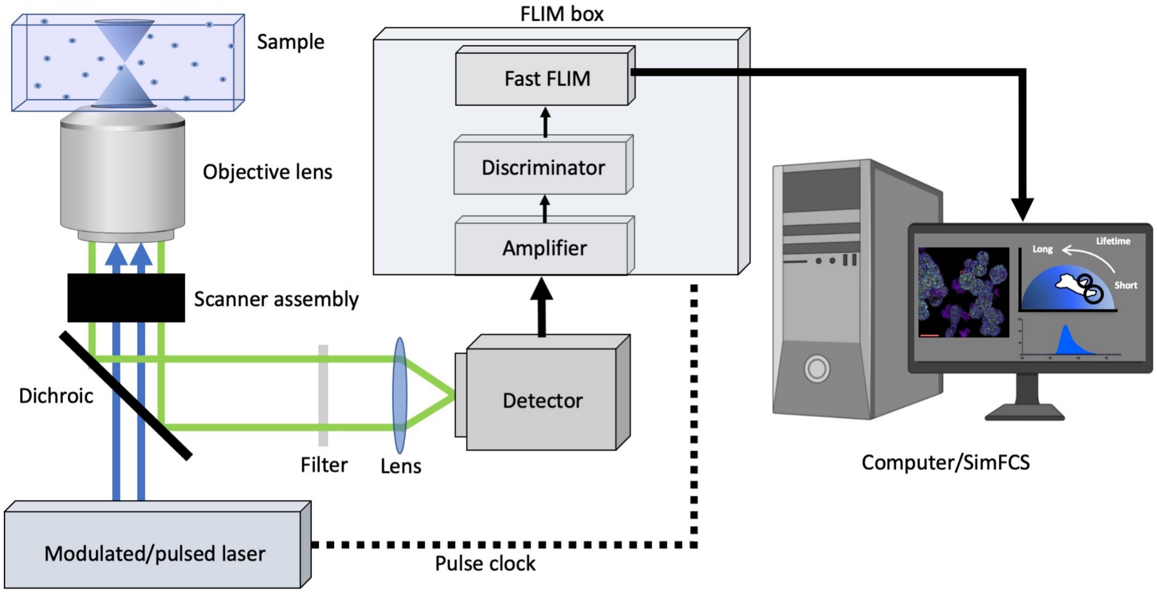


***Supplementary Figure 7. Generalized spectral-FLIM Microscopy setup****. A pulsed/modulated light source is used to illuminate the sample and the fluorescence of the sample is collected by a spectral detector (current resolution around 10 nm). The repetition rate can either be supplied by or delivered to the laser which is used by the electronics in the digital frequency domain to obtain a single photon arrival time using the heterodyne principle (current resolution around 50 ps)*

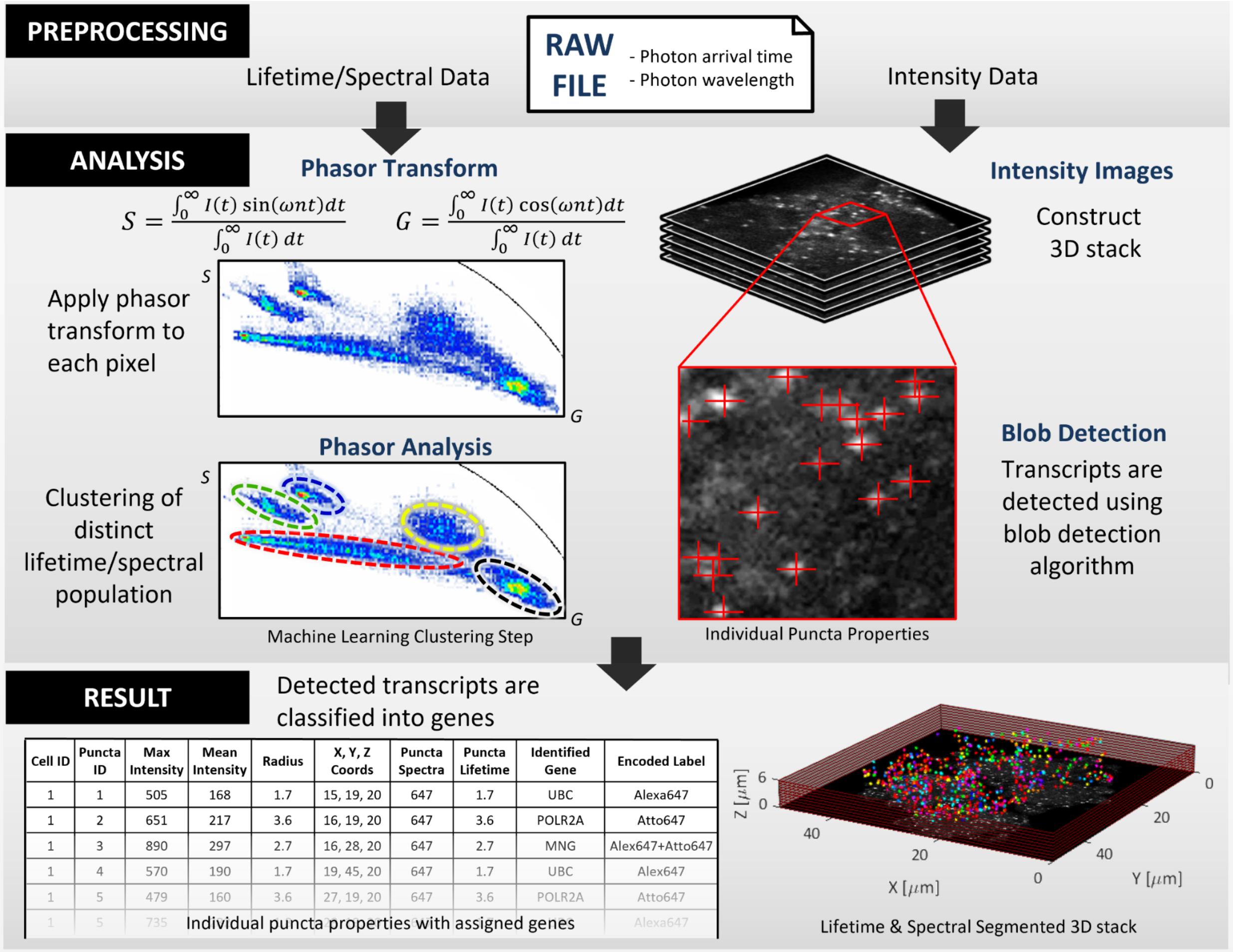


| Cell ID | Puncta ID | Max Intensity | Mean Intensity | Radius | X, Y, Z Coords | Puncta Spectra | Puncta Lifetime | Identified Gene | Encoded Label |
|---|---|---|---|---|---|---|---|---|---|
| 1 | 1 | 505 | 168 | 1.7 | 15, 19, 20 | 647 | 1.7 | UBC | Alexa647 |
| 1 | 2 | 651 | 217 | 3.6 | 16, 19, 20 | 647 | 3.6 | POLR2A | Atto647 |
| 1 | 3 | 890 | 297 | 2.7 | 16, 28, 20 | 647 | 2.7 | MNG | Alex647+Atto647 |
| 1 | 4 | 570 | 190 | 1.7 | 19, 45, 20 | 647 | 1.7 | UBC | Alex647 |
| 1 | 5 | 479 | 160 | 3.6 | 27, 19, 20 | 647 | 3.6 | POLR2A | Atto647 |
| 1 | 5 | 735 | [illegible] | [illegible] | [illegible] | [illegible] | [illegible] | [illegible] | Alexa647 |

***Supplementary Figure 8. Automated pipeline of the processing and analysis.*** *Raw data consists of a list of detected photons with their times of arrivals. Using the acquisition parameters, dwell time, number of pixels, number of repetitions per image etc. the image stacks are reconstructed. Knowing the laser frequency, a photon histogram for each voxel is built and the phasor transform is applied. The two custom made algorithms work in parallel, one identifying clusters in the phasor space, the other identifying puncta in the intensity space. The two then recombine to result in each transcript being identified, assigned to a particular gene and its morphological properties measured.*
